# Non-invasive assessment of normal and impaired iron homeostasis in living human brains

**DOI:** 10.1101/2022.05.02.490254

**Authors:** Shir Filo, Rona Shaharabani, Daniel Bar Hanin, Masha Adam, Eliel Ben-David, Hanan Schoffman, Nevo Margalit, Naomi Habib, Tal Shahar, Aviv Mezer

## Abstract

Strict iron regulation is essential for normal brain function. The iron homeostasis, determined by the milieu of available iron compounds, is impaired in aging, neurodegenerative diseases and cancer. However, non-invasive assessment of different molecular iron environments implicating brain tissue’s iron homeostasis remains a challenge. We present a novel magnetic resonance imaging (MRI) technology sensitive to the iron homeostasis of the living brain (the r1-r2* relaxivity). *In vitro*, our MRI approach reveals the distinct paramagnetic properties of ferritin, transferrin and ferrous iron. In the *in vivo* human brain, we validate our approach against ex vivo iron compounds quantification and gene expression. Our approach varies with the iron mobilization capacity across brain regions and in aging. It reveals brain tumors’ iron homeostasis, and enhances the distinction between tumor tissue and non-pathological tissue without contrast agents. Therefore, our approach may allow for non-invasive research and diagnosis of iron homeostasis in living human brains.

**Graphical abstract:** Non-invasive assessment of normal and impaired iron homeostasis in living human brains.

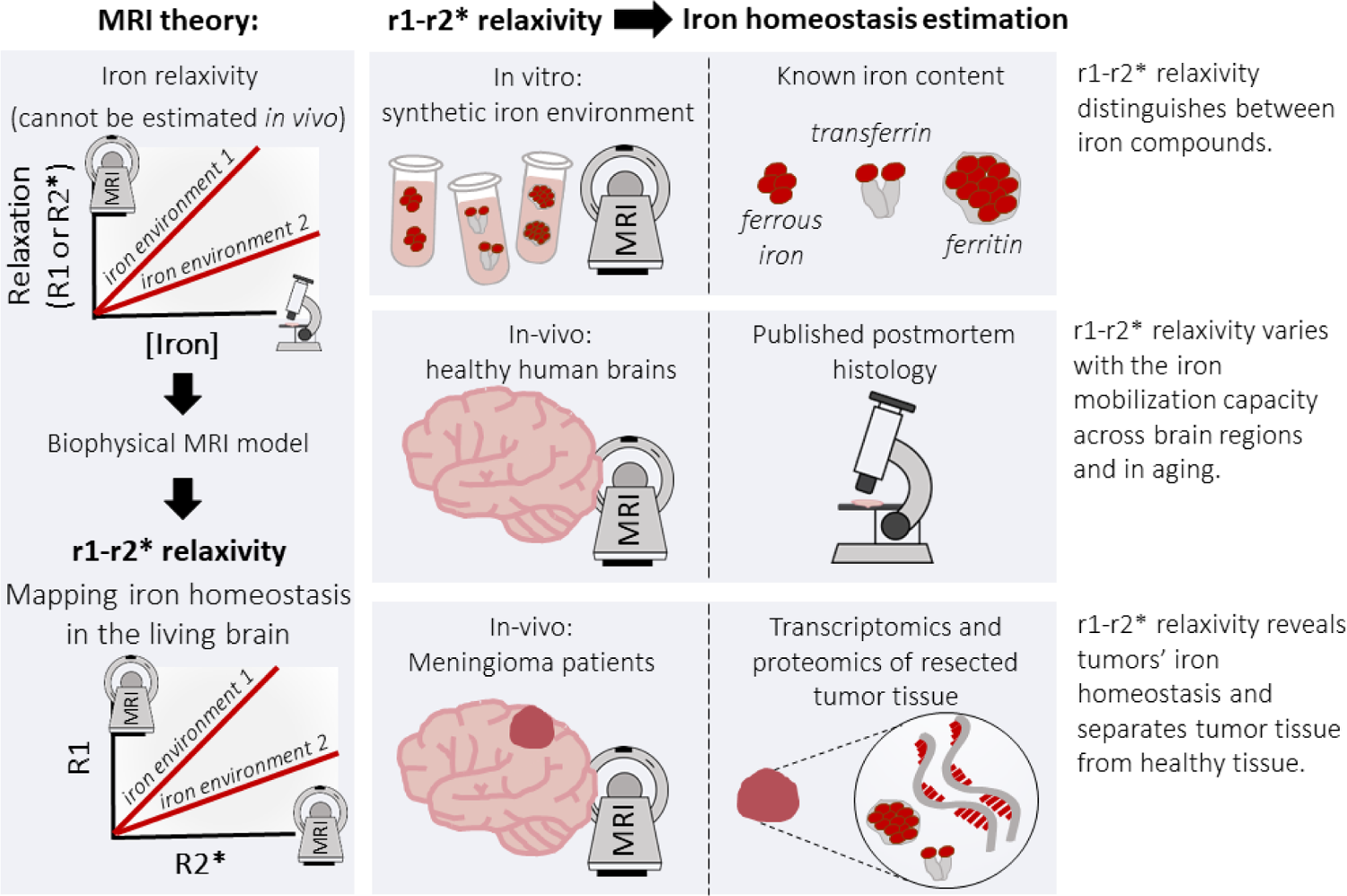

## Introduction

Iron is the most abundant trace element in the human body^1^. It participates in fundamental processes such as oxygen transport, cellular metabolism, myelin formation and the synthesis of neurotransmitters^1–4^. Therefore, strict iron regulation is essential for maintaining normal brain function. Brain tissue’s iron homeostasis could be characterized by the tissue’s molecular iron environment, determined by the specific milieu of available iron compounds, their iron binding capacities and aggregation states. Importantly, the molecular iron environment varies between cell types and across brain regions^3, 5–7^.

Disrupted iron homeostasis plays a major role in normal aging and in neurodegenerative diseases such as Parkinson’s disease (PD), Alzheimer’s disease (AD), multiple sclerosis, Friedreich’s ataxia, aceruloplasminaemia, neuroferritinopathy, Huntington’s disease, and restless legs syndrome^1, 2, 5–10^. The two iron compounds most involved in iron homeostasis are transferrin and ferritin^3^. Transferrin, the main iron transport protein, carries iron from the blood into brain tissue, while ferritin, the main iron storage protein, stores excess iron atoms. When iron concentrations exceed the capacity of iron-binding proteins, this can lead to oxidative stress and cellular damage^10^. For example, the ratio of transferrin to iron, which reflects iron mobilization capacity, differs between elderly controls and patients (AD and PD) in a brain-region–dependent manner^7^. In addition, specifically in the substantia nigra and the locus coeruleus, reduction in neuromelanin-iron complexes is considered a biomarker for PD and AD^11, 12^.

Impaired homeostasis of the molecular iron environment also have been reported in cancer cells^13, 14^. Tumor cell proliferation requires a modulated expression of proteins involved in iron uptake. In addition, iron may affect the immune surveillance of tumors^15^. Therefore, the availability of iron in the tumor cells’ microenvironment may affect their survival and growth rate, and subsequently the course of the disease. For example, meningioma brain tumors^16^, compared to non-pathological tissue, were shown to contain a higher concentration of ferrimagnetic particles and abnormal expression of iron-related genes^17, 18^. These findings suggest there are detectable differences in iron homeostasis between brain tumors and normal brain tissue.

The extensive implications of impaired iron homeostasis in normal aging, neurodegeneration and carcinogenesis suggest that assessment of iron homeostasis in the living brain would be highly valuable for diagnosis, therapeutic monitoring, and understanding pathogenesis of diseases^4^. Iron’s paramagnetic properties make magnetic resonance imaging (MRI) a perfect candidate for non-invasive estimation of iron content in brain tissue. In particular, iron is a major contributor to the longitudinal and effective transverse relaxation rates, R1 and R2* respectively^19–22^. These relaxation rates can be measured using quantitative MRI (qMRI) techniques^23–26^. Indeed, *in vivo* studies often use these qMRI measurements as a proxy for iron presence in brain tissue^21, 27–31^. However, a major limitation of current MRI techniques is that they lack information regarding the state of iron homeostasis, as they do not have the sensitivity to discriminate between different molecular environments of iron in the brain^4^.

Early *in vitro* and postmortem works suggest that different iron environments can be distinguished by their iron relaxivity^31–35^. The iron relaxivity is defined as the dependency of MR relaxation rates on the iron concentration^36^. It was shown that iron relaxivity varies with the specific environment in which the iron resides^31–35^. However, a major limitation of this approach is that it requires direct estimation of the tissue iron concentration, which can only be acquired *in vitro* or postmortem. Therefore, until now the phenomenon of iron relaxivity could not be studied in living humans.

Here we propose an *in vivo* iron relaxivity approach sensitive for the state of iron homeostasis in the brain. Our approach fully relies on MRI parameters, and does not require estimation of the tissue iron concentration, thereby allowing for the first time non-invasive assessment of different molecular iron environments in the living brain. We exploit the distinct iron relaxivities of the MR relaxation rates, R1 and R2*, to construct a biophysical model of their linear interdependency, which we label the r1-r2* relaxivity. Using the r1-r2* relaxivity, we argue that the distinct iron relaxivity of different molecular iron environments can be estimated *in vivo*. We confirm this hypothesis based on a novel validation framework. First, we used a bottom-up strategy in which we evaluated the r1-r2* relaxivity of different iron environments *in vitro*. Next, we used a top-down strategy in which we measured the r1-r2* relaxivity in human brains *in vivo* and compared it to ex vivo quantification of iron compounds and gene expression, both at the group and the single-subject levels. In healthy subjects, we assessed the biological correlates of the r1-r2* relaxivity, and compared it to other MR contrasts across brain regions and in aging. In meningioma patients, we tested the ability of our approach to enhance the distinction between tumor tissue and non-pathological tissue, and to reveal the state of iron homeostasis in tumors. Therefore, we provide a well-validated MRI framework with great implications for the non-invasive research and diagnosis of normal and impaired iron homeostasis in living human brains (see graphical abstract).

## Results

### The theoretical basis for the r1-r2* relaxivity

The iron relaxivity is defined based on the linear relationship between the relaxation rates (R1 and R2*) and the iron concentration ([IC])^36^:

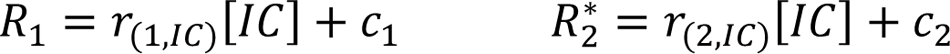

The slopes of these linear dependencies, r_(1,IC)_ and r_(2,IC)_, represent the iron relaxivities of R1 and R2*, which were shown to have different values for different iron environments^31–34^. c_1_ and c_2_ are constants. Notably, the iron relaxivities require estimation of the iron concentration ([IC]), thereby limiting this approach to *in vitro* and ex vivo studies.

Here we propose a theory which advances the relaxivity model and provides *in vivo* iron relaxivity measurements for identifying different iron environments in the brain. We take advantage of the fact that R1 and R2* are governed by different molecular and mesoscopic mechanisms^37–39^, and therefore each of them may have a distinct iron relaxivity in the presence of paramagnetic substances. Based on our theoretical framework (“*In vivo* iron relaxivity model” in Methods), the linear dependency of R1 on R2* can be described by the following equation:

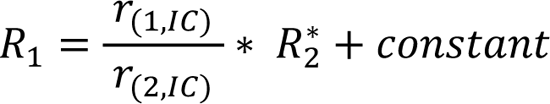

The slope of this linear dependency 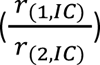 represents the ratio of the iron relaxivities, r_1_ and r_2_^∗^, which are sensitive to the molecular environment of iron. Therefore, we define this MRI-based slope as the r1-r2* relaxivity 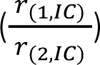 and hypothesize it reveals the distinct properties of different molecular iron environments. Notably, in the brain, the r1-r2* relaxivity could be affected by the entire milieu of available iron compounds, their iron binding capacities and aggregation states (i.e. the molecular iron environment). For an extension of the relaxivity model in the presence of a heterogeneous iron environment and myelin see Supplementary Section 4.1: The theoretical basis for the r1-r2* relaxivity of brain tissue.

### In vitro validation for the sensitivity of the iron relaxivity to molecular iron environments

Before implementing this approach in the living human brain, we validated our theory by manufacturing *in vitro* samples of different iron environments in a synthetic cellular membrane environment. These samples were scanned in the MRI to verify that different iron environments have different iron relaxivities. We then tested whether our r1-r2* relaxivity theory could reveal these different relaxivities.

We prepared samples of transferrin, ferritin and ferrous iron in different cellular-like environments (free in water or adjacent to liposomes and proteins, to achieve physiological iron concentrations the transferrin concentrations are higher than the ones measured *in vivo*^3, 4^). These highly controlled synthetic iron environments were scanned for R1 and R2* mapping. We found that both R1 and R2* increased with the concentration of iron compounds (Figure 1a-b). The rate of this increase, defined as the iron relaxivity, was different for different iron environments (Figure 1a-c, p(ANCOVA)<10^-50^). We show that R1 and R2* change both with the type and concentration of iron, thereby making it impossible to distinguish between iron environments with these measurements. For example, R1 increased with the ferritin concentration, but also was higher for ferritin compared to transferrin (Figure 1a-b, Sup. Figure 1a-b). Consequently, similar R1 values can be obtained for ferritin, transferrin and ferrous iron, depending on their concentrations (Sup. Figure 1b, Figure 1a). This ambiguity can be resolved by the iron relaxivity, which differentiated the iron environments, and was consistent when computed over samples with higher or lower concentrations (Sup. Figure 1c). Therefore, we find that the iron relaxivity reflects changes in the molecular environment of iron and is independent of the iron concentration. Ferritin binds thousands iron ions while transferrin binds only few^3^. Thus, we wanted to exclude the possibility that the relaxivity differences were being driven by the different iron ion concentration. We estimated the iron ion concentrations for the different molecular iron environments (see Methods), and verified that ferritin, transferrin and ferrous iron indeed have different iron relaxivities even when accounting for the discrepancies in iron concentrations (Supplementary Section 1: The dependency of R1 and R2* on the iron concentration.).

**Figure 1:**
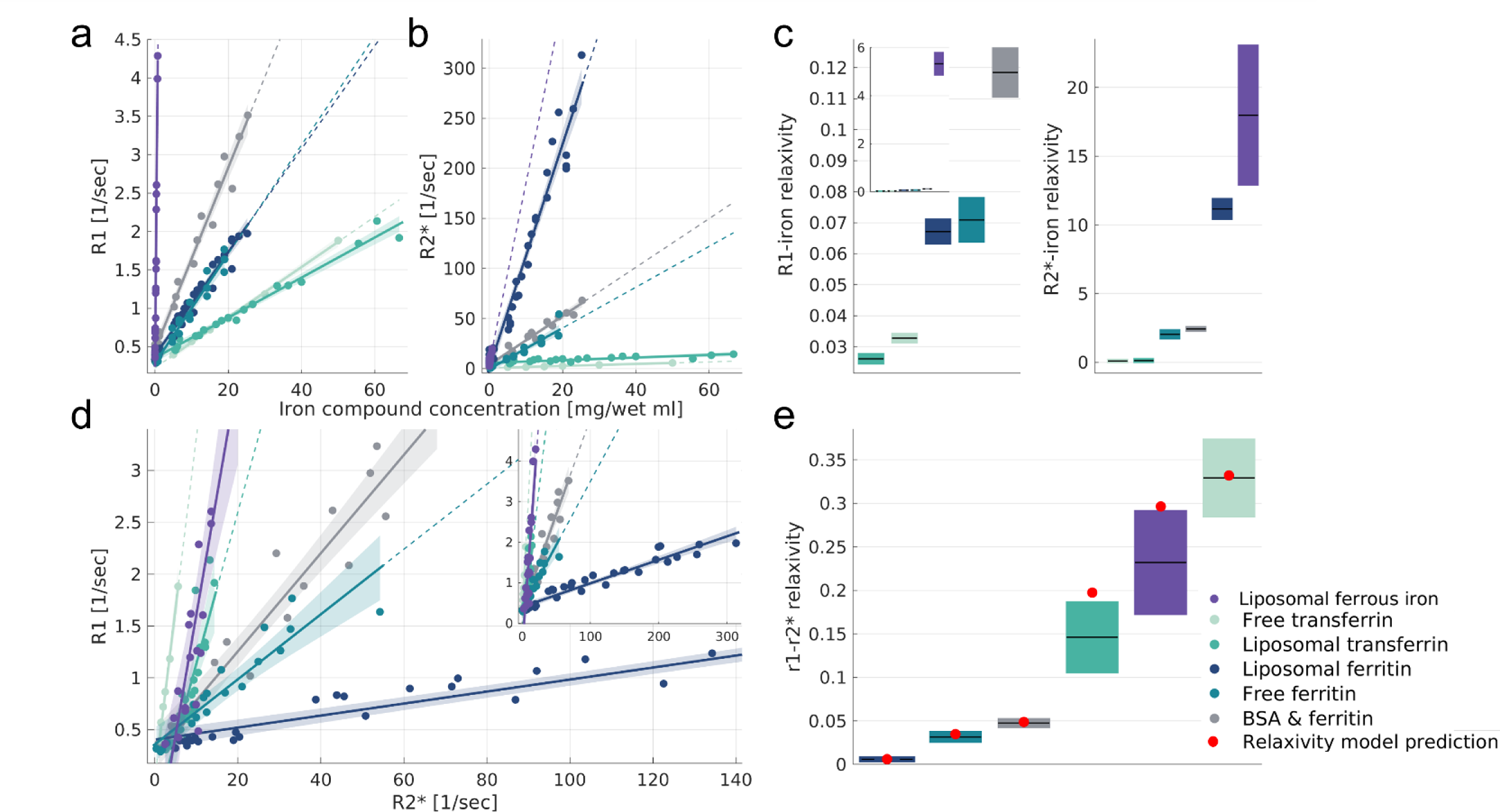
In vitro validation of the non-invasive framework for assessing the iron environments. (a-b) The dependency of R1 and R2* on the iron compound concentrations for different iron environments: free ferritin, liposomal ferritin, bovine serum albumin (BSA)-ferritin mixture, free transferrin, liposomal transferrin and liposomal ferrous iron. Data points represent samples with varying concentrations relative to the water fraction ([mg/wet ml]). The linear relationships between relaxation rates and iron-compounds concentrations are marked by lines. We define the slopes of these lines as the iron relaxivities. Dashed lines represent extrapolation of the linear fit. Shaded areas represent the 95% confidence bounds. (c) The iron relaxivity of R1 and R2* is different for different iron environments (p(ANCOVA)<10^-50^). Iron relaxivities are calculated by taking the slopes of the linear relationships shown in (a,b), and are measured in [sec^-1^/(mg/wet ml)]. For each box, the central mark is the iron relaxivity (slope); the box shows the 95% confidence bounds of the linear fit. For the R1-iron relaxivity, the inset shows a zoom-out of the main figure, presenting the entire range of measured values (d) The dependency of R1 on R2* for different iron environments. Data points represent samples with varying iron compound concentrations relative to the water fraction. The linear relationships of R1 and R2* are marked by lines. The slopes of these lines are the r1-r2* relaxivities, which do not require iron concentration estimation and therefore can be estimated in vivo. Dashed lines represent extrapolation of the linear fit. Shaded areas represent the 95% confidence bounds. The inset shows a zoom-out of the main figure, presenting the entire range of measured R2* values (e) The r1-r2* relaxivities are different for different iron environments (p(ANCOVA)<10^-38^). For each box, the central mark is the r1-r2* relaxivity, and the box shows the 95% confidence bounds of the linear fit. Red dots indicate the successful prediction of the experimental r1-r2* relaxivity from the ratio between the iron relaxivities of R1 and R2* (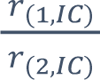, shown in c). This validates our theoretical in vivo relaxivity model.

### The r1-r2* relaxivity reveals the distinct iron relaxivities of different iron environments

In agreement with previous findings^31–34^, our *in vitro* experiments indicate that iron relaxivity can be used to identify different iron environments. While the iron relaxivity cannot be estimated *in vivo*, as it requires measurements of iron concentrations, the r1-r2* relaxivity only relies on MRI measurements that can be estimated *in vivo*. Based on our theory, we argue that two iron environments with different iron relaxivities are also likely to differ in their r1-r2* relaxivities. We validated this hypothesis using synthetic iron-containing samples. As predicted by our theoretical model, iron environments with different iron-relaxivities had different r1-r2* relaxivities (Figure 1d-e, p(ANCOVA)<10^-38^). Notably, as suggested by our theoretical formulation, the r1-r2* relaxivity provides a good MRI approximation for the ratio between the iron relaxivities of R1 and R2* (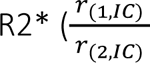, Figure 1e). Similar to the iron relaxivities, the r1-r2* relaxivity is consistent across iron concentrations (Sup. Figure 1d). Hence, the r1-r2* relaxivity is more sensitive to the molecular iron environment than R1 and R2* by themselves. In addition, we validated that the r1-r2* relaxivity is sensitive to the paramagnetic properties of iron-binding proteins. We found that apo-transferrin (transferrin which is not bound to iron) has a much smaller r1-r2* relaxivity compared to iron-bound transferrin (p(ANCOVA)<10^-8^; Sup. Figure 4). This implies that it is paramagnetic properties that induce the r1-r2* relaxivity that we measure. Taken together, these results validate our theory, indicating that the r1-r2* relaxivity can be used to measure iron relaxivity *in vivo* for exposing the distinct paramagnetic properties of different molecular iron environments.

Brain tissue includes a complex milieu of iron compounds and myelin, which is a major contributor to R1 and R2*^4,19,25, 29, 40–45^. Therefore, we tested *in vitro* the r1-r2* relaxivity of a heterogenous molecular iron environment of ferritin-transferrin liposomal mixtures (Supplementary Section 4.2: The r1-r2* relaxivity of ferritin and transferrin mixtures *in vitro*.). We found that changing the transferrin-ferritin ratio leads to considerable changes in the r1-r2* relaxivity, even in mixtures with low ratio of transferrin compared to ferritin as in the brain^3^. Importantly, these changes were above the detection limit of the *in vitro* r1-r2* relaxivity measurement (Sup. Figure 8). Since myelin is composed mainly of lipids, we tested the effect of the myelin fraction on the iron relaxivity by varying the liposomal fractions in our *in vitro* experiments. We found that the r1-r2* relaxivities are stable for different liposomal fractions and lipid types (for more details see Supplementary Section 2: The dependency of the iron relaxivity on the liposomal fraction.). These results highlight the specificity of the r1-r2* relaxivity to differences in the molecular state of iron, unlike the ambiguous measurements of R1 or R2* independently.

### The r1-r2* relaxivity provides a new MRI contrast in the in vivo human brain

Following the *in vitro* validation, we measured the *r1-r2* relaxivity* in the living human brain. For this aim we calculated the linear dependency of R1 on R2* across voxels of different anatomically-defined ROIs (see “r1-r2* relaxivity computation for ROIs in the human brain” in Methods). We found distinct r1-r2* relaxivities for different brain regions (Figure 2a). This indicates a heterogeneous distribution of the *in vivo* iron relaxivity across the brain, which is consistent across healthy subjects (age 27±2 years, N=21, Figure 2b) and is reproducible in scan-rescan experiments (Sup. Figure 9).

**Figure 2:**
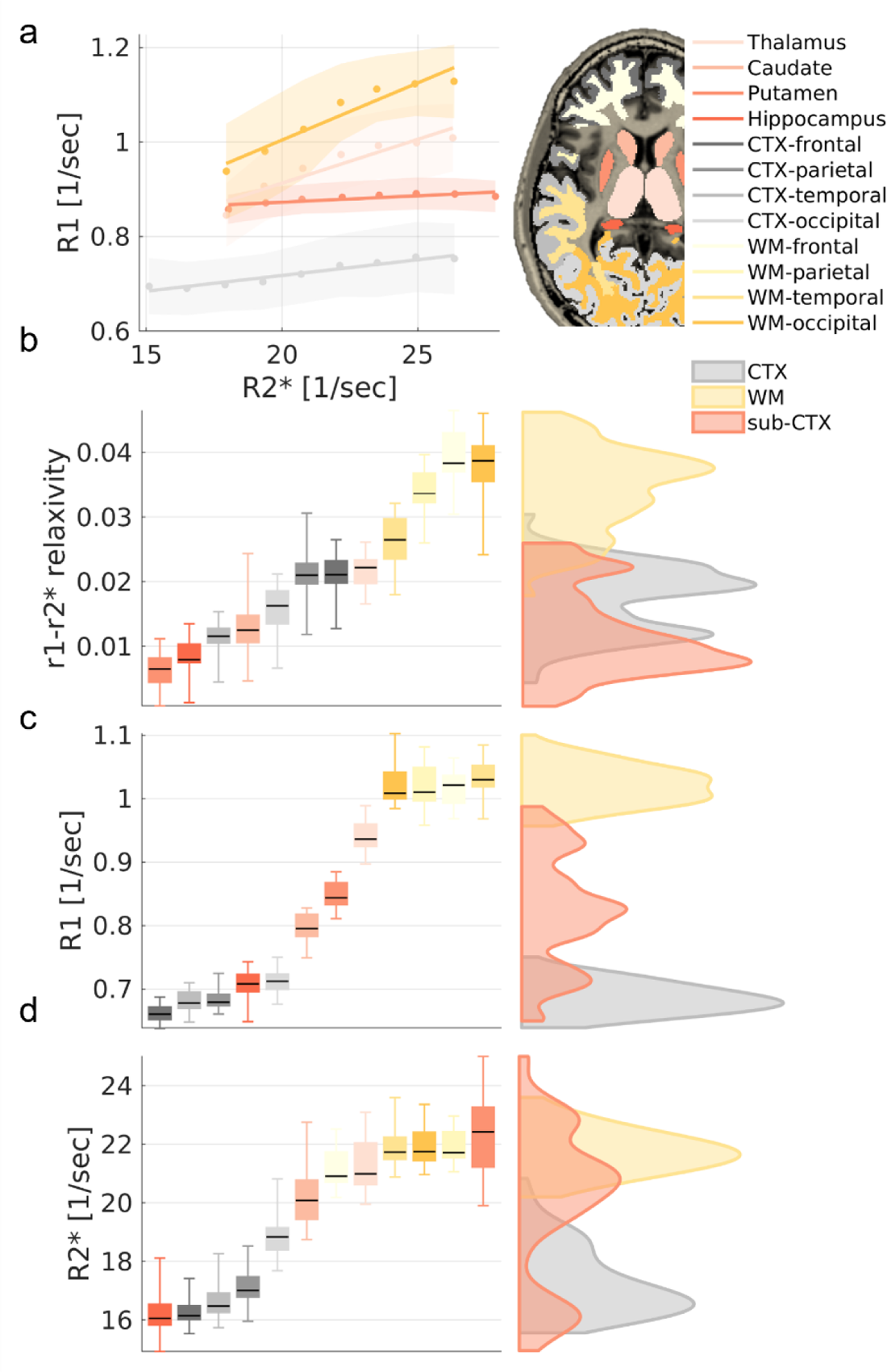
The in vivo r1-r2* relaxivity provides a novel contrast in the brain. (a) The dependency of R1 on R2* in four representative brain regions (WM-occipital, CTX-occipital, Thalamus & Putamen) of a single subject. R2* and R1 were binned (dots represent the median; shaded areas represent the mean absolute deviation), and a linear fit was calculated. The slopes of the linear fit represent the dependency of R1 on R2* (r1-r2* relaxivity) and vary across brain regions. (b) The r1-r2* relaxivity across the brain. Left: the reliability of the method in different brain regions as observed by the variation in the r1-r2* relaxivity across normal subjects (age 27±2, N=21). The 25th, 50th and 75th percentiles and extreme data points are shown for each box. Right: the contrast of the r1-r2* relaxivity across the brain. Red, yellow and gray distributions represent the values of the r1-r2* relaxivities in sub-cortical (sub-CTX), white-matter (WM) and cortical (CTX) brain regions, respectively. (c-d) Similar analyses for R1 and R2* values, in which the gray-matter vs. white-matter contrast is much more dominant compared to the r1-r2* relaxivity. Hence, the r1-r2* relaxivity provides new information compared to R1 and R2*, beyond the WM-GM. Results in this entire figure are for ROIs in the left hemisphere. WM=white-matter, CTX=cortex.

In agreement with our *in vitro* results, which indicated that the r1-r2* relaxivity provides different information compared to R1 and R2*, in the *in vivo* human brain we find that the r1-r2* relaxivity produces a new contrast, statistically different from R1 and R2* (Figure 2b-d; p<0.05 for the two-sample Kolmogorov–Smirnov test comparing the r1-r2* relaxivity distribution to R1, and p<0.001 comparing it to R2*). While the r1-r2* relaxivity is calculated for an anatomically-defined ROI in the brain, a voxel-wise r1-r2* relaxivity visualization based on each voxel’s local neighborhood, as well as comparison to the R1 and R2* contrasts, is demonstrated in Supplementary Section 3: Voxel-wise r1-r2* relaxivity visualization.

### The effect of myelin on the r1-r2* relaxivity

The sensitivity of R1 and R2* to the myelin content is known to produce contrasts that are governed mainly by the differences between white-matter and gray-matter tissues^4, 19, 25, 29, 40–44^. As expected, we find a strong distinction between gray-matter and white-matter regions in R1 and R2* values (Figure 2c-d). However, the contrast of the r1-r2* relaxivity across the brain shows a novel spatial pattern and reveals differences between brain regions beyond the typical white matter—gray matter differentiation. For example, we found the temporal, parietal and occipital white-matter regions to be indistinguishable in terms of their R1 and R2* values (p(ANOVA)>0.4), but these regions were separable based on their different r1-r2* relaxivities (p(ANOVA)<10^-10^, Sup. Figure 13).

To further establish that the r1-r2* relaxivity is less sensitive to the myelin content relative to R1 and R2* we compared it to several *in vivo* myelin markers. The qMRI measurements of the macromolecular tissue volume (MTV)^46^,magnetization transfer saturation (MTsat)^47^, and mean diffusivity (MD) were shown to approximate the myelin content and characteristics^48–53^. These myelin-sensitive markers were highly correlated with R1 and R2* but were not significantly correlated with the r1-r2* relaxivity (Figure 3a, Sup. Figure 14-15). In a biophysical model of the r1-r2* relaxivity, that accounts for the presence of myelin and iron compounds, we find that the variability of myelin within an ROI can affect the r1-r2* relaxivity measurement (Supplementary Section 4.3: Numerical simulations of the r1-r2* relaxivity.). Yet, an *in vivo* estimate of this myelin characteristic revealed it explains only 30% of the variability in the *in vivo* r1-r2* relaxivity across the brain (Sup. Figure 27). We also performed a set of numerical simulations in which we consider the contributions of multiple brain tissue components to the relaxivity measurement (Supplementary Section 4.3: Numerical simulations of the r1-r2* relaxivity.). As in the *in vivo* brain, we found that changes in the myelin concentration substantially affect the simulated measurements of R1 and R2*. However, myelin-related changes were not the main component governing the simulated measurement of the r1-r2* relaxivity, and in simulations of physiological conditions they could not explain the variability in the r1-r2* relaxivity across the brain.

**Figure 3:**
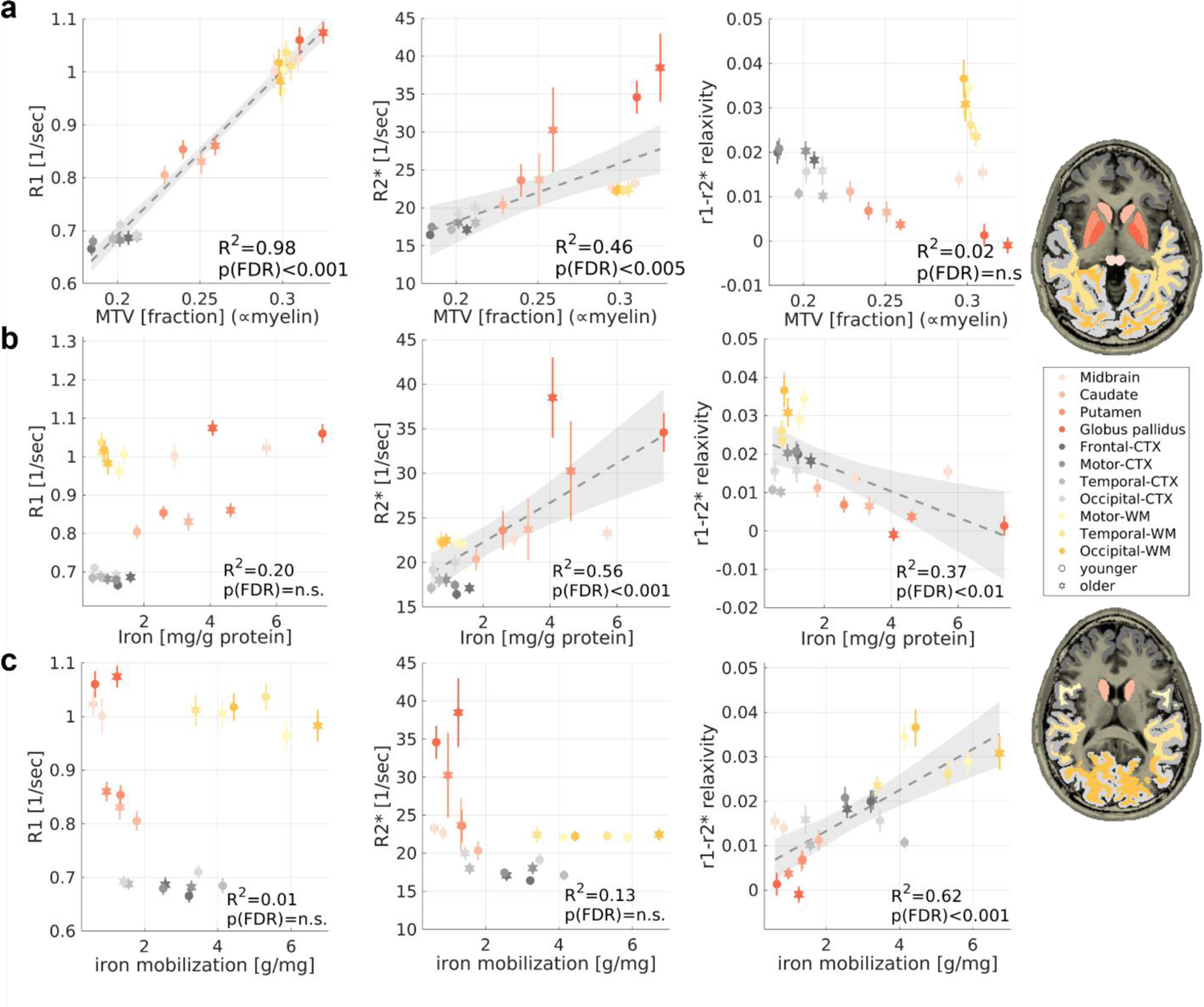
The r1-r2* relaxivity has unique biological correlates compared to R1 and R2* across the brain and in aging. R1, R2* and the r1-r2* relaxivity were measured in vivo across younger (aged 23-63 years, N =26) and older (aged 65-77 years, N=13) subjects (different marker shapes) in 11 brain regions (different colors). Each row presents the correlations of these qMRI measurements with a different in vivo or ex vivo histological feature (fitted model and 95% confidence bounds are presented for significant correlations): (a) qMRI vs. the macromolecular tissue volume (MTV), an in vivo myelin-sensitive marker, measured for younger (aged 23-63 years, N =26) and older (aged 65-77 years, N=13) subjects. Unlike R1 and R2*, the r1-r2* relaxivity is not linearly related to this myelin-sensitive marker (See **Sup. Figure 14, Sup. Figure 15** for additional in vivo myelin-sensitive markers) (b) qMRI vs. the iron concentration (postmortem, from the literature^5,7^) measured for younger (aged 27-64 years, N>=7) and older (aged 65-88 years, N>=8) subjects. Please note that the r1-r2* relaxivity is not significantly correlated with the iron content when excluding the outlier values in the globus pallidum while the R2* correlation survives this exclusion (see **Sup. Figure 31**). (c) qMRI vs. the iron mobilization capacity (transferrin/iron ratio), an iron homeostasis marker (postmortem, from the literature^5,7^, same subjects as in b). Only the r1-r2* relaxivity is significantly correlated with the iron mobilization capacity, implying for its sensitivity to the iron homeostasis across brain regions and in aging. WM=white-matter, CTX=cortex.

### The r1-r2* relaxivity correlates with the iron mobilization capacity across the brain and in aging

Next, we tested the sensitivity of the *r1-r2* relaxivity* to the state of iron homeostasis across the normal brain and in aging. We aggregated previously reported postmortem histological data describing iron, ferritin and transferrin concentrations in different brain regions of young (aged 27-64 years, N>=7) and older (aged 65-88 years, N>=8) adults^5,7,9^ (see “Group-level comparison of qMRI parameters and histological measurements” in Methods). We performed a group-level comparison between these postmortem findings and *in vivo* MRI parameters, which we measured in the same brain regions and age groups (healthy young subjects aged 23-63 years, N=26; older subjects aged 65-77 years, N=13). As expected, R2* was significantly correlated with iron concentration (R^2^=0.41, p-value (FDR)<0.05; Figure 3b). Importantly, this result validates the agreement between the *in vivo* and postmortem datasets, thus allowing to further examine the biological correlates of the r1-r2* relaxivity. For this aim we estimated a feature of the iron homeostasis, the iron mobilization capacity (transferrin/iron^7^), available in the postmortem dataset. This measure was not correlated with R2* or R1 (Figure 3c). However, the iron mobilization capacity^5,7^ was significantly correlated with the r1-r2* relaxivity across brain regions and age groups (R^2^=0.59, p-value(FDR)<0.001, Figure 3c). Hence, the r1-r2* relaxivity, unlike R1 and R2*, is sensitive to the obtained *ex vivo* measurements of the iron mobilization capacity across the brain and can capture the effect of aging on this iron homeostasis marker.

The r1-r2* relaxivity was much less correlated with the absolute ferritin, transferrin or iron concentrations (Figure 3b, Sup. Figure 18). Moreover, the correlation with the iron concentration was driven mostly by outliers (the globus pallidum), unlike the sensitivity of the r1-r2* relaxivity to the iron homeostasis marker (Supplementary Section 5: The r1-r2* relaxivity in the pallidum.). Therefore, the r1-r2* relaxivity is less sensitive to absolute changes in the iron and myelin concentrations. In return, these results imply that the r1-r2* relaxivity enhances the sensitivity of MRI to the iron homeostasis. We corroborated these findings with numerical simulations of the r1-r2* relaxivity of brain tissue. In these simulations we further demonstrated that the measured changes in the r1-r2* relaxivity across the brain can be induced by changes in the iron environment (Sup. Section 4.3). Moreover, we find that the effect of the iron environment on the r1-r2* relaxivity is not confounded by the myelin and iron concentrations and is well above the detection limit of this MRI measurement (Sup. Figure 9). These analyses suggest that the *in vivo* r1-r2* relaxivity is capable of revealing important characteristics of the iron environment previously inaccessible with MRI.

### The r1-r2* relaxivity enhances the distinction between tumor tissues and non-pathological tissue

While the r1-r2* relaxivity forms a unique pattern of changes across the brain, it needs to be established that this contrast contains meaningful clinical information, that can complement the contrasts of R1 and R2*. For this aim, we evaluated the MRI contrast between pathological and normal-appearing tissues of patients with meningioma brain tumors (N=18, Figure 4a-b). The diagnosis of brain tumors and their delineation from the surrounding non-pathological tissue is routinely performed using contrast-enhanced MRI, which requires the injection of an external gadolinium (Gd)-based contrast agent with paramagnetic properties^54^. As expected, when using Gd-based contrast, tumor tissue was distinct from white-matter and gray-matter tissues (Figure 4c, Cohen’s d=1.4, p<10^-4^ for tumor-gray matter; Cohen’s d=1.18, p<10^-3^ for tumor-white matter). Recently renewed concerns about the long-term safety of Gd-based agents^55, 56^, highlight the need for Gd-free MRI techniques that can serve as safe alternatives^57^. However, without Gd-agent injection, both for R1 and R2* values the biggest effect size was observed between white-matter and gray-matter tissues, with no significant difference between gray-matter and tumor tissues (Cohen’s d<0.45, p>0.08 for R1, Cohen’s d<0.11, p>0.65 for R2*, Figure 4d-e). This demonstrates the poor performances of R1 and R2* in Gd-free tumor tissue delineation. Importantly, the r1-r2* relaxivity greatly enhanced the contrast between tumor tissue and non-pathological tissue, without contrast agent injection (Figure 4f, Cohen’s d=1.5, p<10^-5^ for tumor-gray matter; Cohen’s d=4.32, p<10^-11^ for tumor-white matter). This Gd-free enhancement was comparable in size to the effect of Gd-based contrast. Moreover, we provide an example for a voxel-wise visualization of the r1-r2* relaxivity approach in a representative meningioma patient, which demonstrates the Gd-free tumor enhancement (Supplementary Section 3: Voxel-wise r1-r2* relaxivity visualization. These results emphasize the improved sensitivity of the r1-r2* relaxivity to the unique tumor microenvironment, which may have wide clinical implications as a safe alternative for contrast agents’ injections.

**Figure 4:**
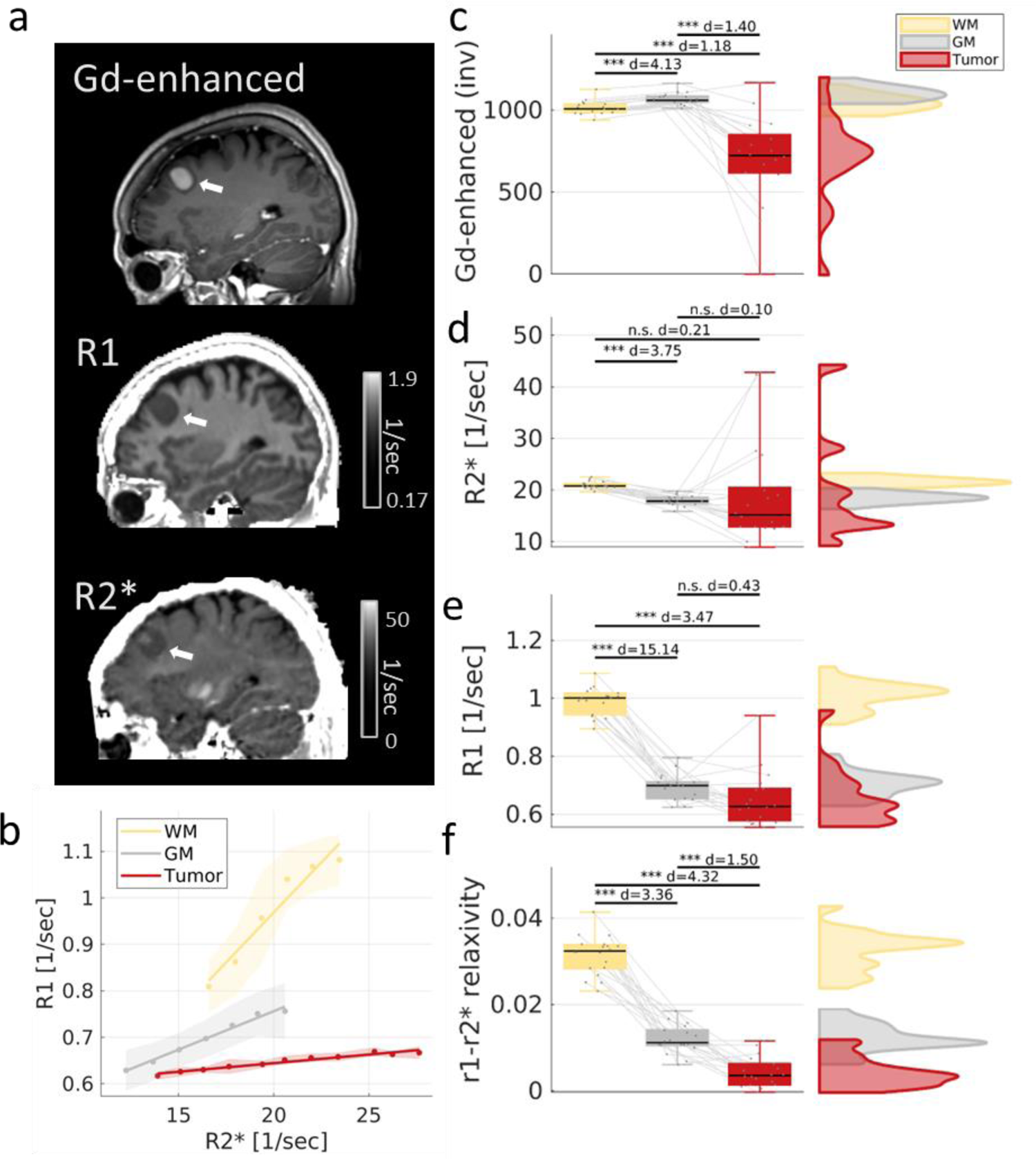
Application of the r1-r2* relaxivity on meningioma brain tumors. (a) From top to bottom: Gd-enhanced T1-weighted image, R1 map and R2* map in a representative subject with a meningioma brain tumor (white arrow). (b) The dependency of R1 on R2* (the r1-r2* relaxivity) for the white matter (WM, frontal), gray matter (GM, frontal) and tumor tissue of the same subject. Tumor tissue exhibits distinct r1-r2* relaxivity relative to non-pathological tissue, as evident by the slopes of the R1-R2* linear dependency. (c-f) The contrast between the white matter (WM), gray matter (GM) and tumor tissues across patients (N=18) for the Gd-enhanced contrast (inverted for visualization, a.u.) (c), R2* (d), R1 (e) and the r1-r2* relaxivity (f). Only the r1-r2* relaxivity produces significant differences between tumor and GM tissues without contrast agents. Left: boxes present the variation in the contrasts. The 25th, 50th and 75th percentiles and extreme data points are shown. The d-values represent the effect size (Cohen’s d) of the differences between tissue types, and the significance level is based on a t-test. Gray lines extend between values of the same patient. Right: the distribution of the values between WM, GM and tumor tissue across patients. Estimates in non-pathological tissues are for the tumor-free hemisphere. p<0.05; **p<0.01; ***p<0.001

### The r1-r2* relaxivity is associated with unique biological pathways and gene expression profiles

To further examine how the tumor characteristics obtained by the r1-r2* relaxivity differ from the information contained in R1 and R2*, we examined the associations of these *in vivo* MRI measurements with underlying gene-expression profiles for the same tissue. To this end, we analyzed cases in which the MRI scans of the meningioma patients were followed by surgical interventions, to obtain matching resected tumor tissue samples that we profiled by bulk RNA-sequencing (Figure 5a). For these tumor samples (N=17), we performed an unbiased analysis to identify genes and molecular pathways that could be linked with the *in vivo* measured MRI parameters. For each gene we calculated the correlation between the expression level and the *in vivo* MRI measurements (r1-r2* relaxivity, R1 and R2*) across patients. We then performed gene set enrichment analysis (GSEA)^58,59^ to identify molecular functions that are significantly associated with each MRI measurement. In total, we found 9, 55 and 59 significantly enriched gene sets for R1, R2* and the r1-r2* relaxivity, respectively (p<0.01 after familywise error rate (FWER) correction; Sup. Table 1). These gene sets define genes linked to a specific biological pathway. Almost half of the significant gene sets were exclusively associated with the r1-r2* relaxivity, and not with R1 or R2* (Sup. Figure 16). The enrichment score represents the degree to which the genes within a set were positively or negatively correlated with MRI measurements. In examining the associations of MRI measures to biological pathways, as reflected in the enrichment score, we found that the r1-r2* relaxivity clustered separately from R1 and R2* (Figure 5b). The clustering results were replicated when performed on the p-value of the enrichment, or on the subset of genes within the top enrichment pathways. This implies that the r1-r2* relaxivity reflects unique cellular and molecular properties, undetectable by the separate analysis of R1 and R2*. Therefore, the *in vivo* r1-r2* relaxivity provides a unique dimension for measuring microstructure and gene expression features across the brain. The gene enrichment analysis that we performed on resected brain tumors (Figure 5b) can provide insights into the biological pathways associated with the r1-r2* relaxivity. The two most enriched pathways for the r1-r2* relaxivity were “immunoglobulin complex” (normalized enrichment score (NES)= −3.62, FWER p-value<0.001; Sup. Figure 17a) and “scavenging of heme from plasma” (NES= −3.27, FWER p-value<0.001; Sup. Figure 17b). While the former may relate to the response of the immune system to the cancerous process^60, 61^, the latter involves the absorption of free heme, a source of redox-active iron^62^. This iron-related pathway was not significantly associated with R1 or R2* (p>0.01). Moreover, we examined the main genes involved in iron regulation: transferrin receptor (TFRC), ferritin heavy-chain polypeptide 1 (FTH1) and ferritin light-chain polypeptide (FTL)^63, 64^. Both TFRC and FTH1 were included in the subset of genes within the top enrichment pathways for the r1-r2* relaxivity, but were not found to be associated with R1 or R2*. These findings provide evidence at the level of gene-expression for the sensitivity of the r1-r2* relaxivity to iron homeostasis.

**Figure 5:**
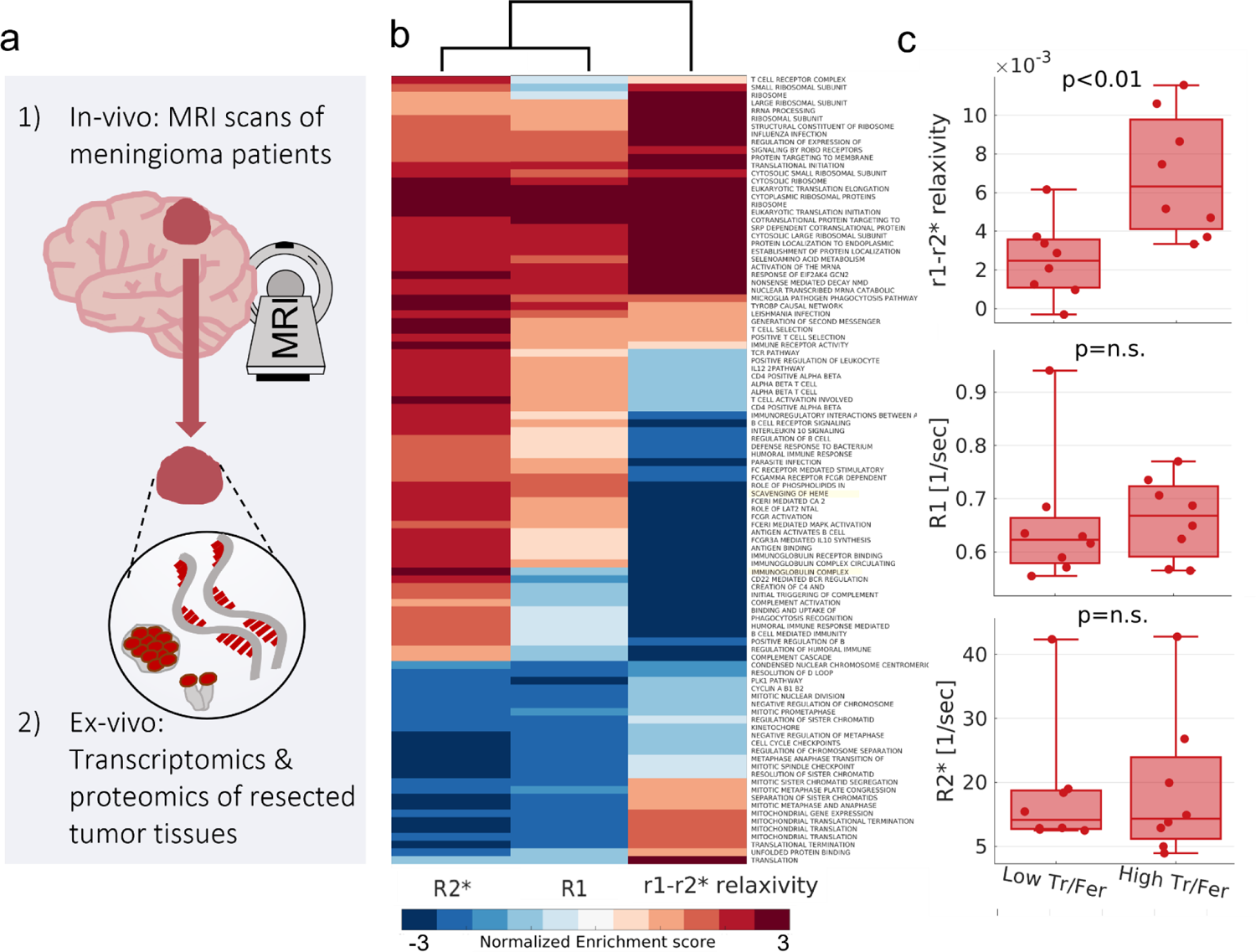
The r1-r2* relaxivity measured in vivo in meningioma patients agrees with iron homeostasis markers estimated ex vivo on surgical specimens of meningiomas from the same patients. (a) Unique comparison between qMRI parameters measured in vivo to ex vivo iron histology of the same human tissue; MRI scans of meningioma patients (N=17) were followed by surgical interventions, to obtain matching resected tumor tissue samples. These tissue samples were profiled by bulk RNA-sequencing to obtain gene expression profiles, and by western-blot analysis to estimate the transferrin/ferritin ratio, a marker for iron homeostasis. (b) Gene set enrichment analysis for the correlation of MRI with gene expression. Rows show significant biological pathways, columns represent R1, R2* and the r1-r2* relaxivity. The dendrogram shows hierarchical clustering of the normalized enrichment scores. The r1-r2* relaxivity clustered separately from R1 and R2* and is therefore enriched for unique biological pathways. The two most enriched pathways for the r1-r2* relaxivity are highlighted in yellow. (c) The r1-r2* relaxivity, R1 and R2* measured in vivo for tumor tissues (N=16) classified as having either low or high transferrin-to-ferritin ratios (Tf/Fer). Tf/Fer ratio, an iron homeostasis marker, was estimated using western-blot analysis following surgical resection of the tissue. The threshold between groups was set to 1 based on the median across subjects. While R1 and R2* cannot distinguish between the groups with different Tf/Fer ratios, the r1-r2* relaxivity is higher in tissues with a high Tf/Fer ratio, indicating its sensitivity to iron homeostasis. p-values presented are for two-sample t-tests.

### The r1-r2* relaxivity reveals differences in iron homeostasis between tumor tissues

We further validated the sensitivity of the r1-r2* relaxivity to iron homeostasis at the proteomics level, by comparing *in vivo* MRI measurements to ex vivo iron homeostasis estimation on the same tissue (Figure 5a). MRI scans of the meningioma patients were followed by surgical interventions, to obtain matching resected tumor tissue samples that we analyzed by western-blot. We compared *in vivo* MRI values of tumor tissue to its transferrin/ferritin ratio which serves as a proxy for iron homeostasis. Neither R1 nor R2* showed significant differences between tumors with low and high transferrin/ferritin ratios (Figure 5c). Notably, the r1-r2* relaxivity was significantly higher for tumors with high transferrin/ferritin ratio compared to tumors with low transferrin/ferritin ratio (p<0.01, Figure 5c). Therefore, as established by both gene expression and proteomics analyses, the r1-r2* relaxivity measured *in vivo* detects pathological disruptions in iron homeostasis which were previously only observable ex vivo.

## Discussion

We present a new MRI relaxivity approach, with increased sensitivity to different molecular environments of iron. First, we confirm *in vitro* that different iron environments induce different relaxivities, which can be estimated with MRI using the r1-r2* relaxivity. When examining R1 and R2* of different iron environments, we find the molecular state of iron is confounded by the strong effects of iron and myelin concentrations. In this *in vitro* setting, we show that the r1-r2* relaxivity reveals the sensitivity of MRI to the intrinsic paramagnetic properties of different iron environments. In the healthy human brain, we show that the r1-r2* relaxivity provides a unique MRI contrast. This contrast is less sensitive to myelin compared to other quantitative MRI parameters. Interestingly, it does vary with an iron homeostasis marker^7^ across brain regions and age groups. We further demonstrate that the r1-r2* relaxivity contains meaningful clinical information associated with iron homeostasis, which was previously inaccessible to conventional MRI approaches. In meningioma patients, we find that the r1-r2* relaxivity contrast is useful for enhancing the distinction between tumor tissue and non-pathological tissue. We substantiate this finding by associating *in vivo* MRI measurements with RNA sequencing and protein expression levels of the same tumor tissues. We find that the r1-r2* relaxivity is associated with unique iron-related biological pathways and reveals the state of iron homeostasis in tumors.

Relaxivity commonly is employed to characterize MR contrast agents^65^. While most contrast agents induce relaxation based on their paramagnetic or superparamagnetic properties, some agents elevate the R1 relaxation rate more efficiently while others elevate R2*. R1 relaxation mechanisms are affected by local molecular interactions, while R2* is sensitive to more global effects of extended paramagnetic interactions at the mesoscopic scale^37^. We show that by contrasting these two different mechanisms, we gain sensitivity to the endogenous iron environment, without the injection of an external contrast agent.

The concept of iron relaxivity, and its sensitivity to the molecular environment of iron, was previously suggested by several postmortem and *in vitro* studies^31–35^. We reproduce these results in our *in vitro* experiments and further demonstrate that different iron environments have different iron relaxivities. Moreover, Ogg et al.^31^ calculated the iron relaxivity by comparing postmortem measurements of iron concentration for different age groups to the typical R1 values in those age groups. They found that this approximation of iron relaxivity was higher in the gray matter and white matter than in sub-cortical structures. Remarkably, we replicate this result in living subjects, based on our novel approach for estimating the iron relaxivity *in vivo*. The theoretical derivation we propose for the r1-r2* relaxivity shows that it represents the ratio of the iron relaxivities of R1 and R2*. This theory was supported by our *in vitro* experiments. To account for the complex molecular environment of the brain, we expanded the biophysical model of the r1-r2* relaxivity for the case of a heterogenous iron environment and in the presence of myelin. Numerical simulations of this model under physiological conditions demonstrate that variations in the iron environment change the r1-r2* relaxivity considerably. Therefore, we exploit the different relaxation rates for a biophysical model of their linear interdependency, thus allowing to approximate the iron relaxivity in the living brain for the first time.

First, we evaluated the sensitivity of the r1-r2* relaxivity to the molecular environment of iron based on *in vitro* experiments with ferritin, transferrin and ferrous iron. We show that these iron environments induce different relaxivities, even when accounting for the discrepancies in iron-binding. Moreover, we show that the macromolecular environment in which the iron reside can alter the relaxivity as well. For example, ferritin induces different relaxivities in the presence of liposomes, proteins or when it is free. As implied previously^32^, this effect can be explained by the encapsulation and the spatial distribution of paramagnetic molecules. Therefore, similarly to different contrast agents, the physical mechanisms by which iron compounds interact with the surrounding water environment are inherently different. To further confirm that our results are related to paramagnetic properties and not the presence of the proteins themself, we tested MRI measurements of apo-transferrin (transferrin unbound to iron). In this case, the R1 relaxivity of transferrin vanished. Therefore, the presence of iron in different *in vitro* environments can be detected by the r1-r2* relaxivity. Moreover, evaluating ferritin-transferrin mixtures, we find that in heterogenous iron environments the specific composition of the iron milieu affects the r1-r2* relaxivity.

The biological interpretation of the r1-r2* relaxivity measurement is more ambiguous when applied to the *in vivo* human brain. This is due to the fact that most qMRI parameters, including R1 and R2*, are known to suffer from low biological specificity in the brain^4,19, 25, 29, 40–42^. The primary MRI contrast between gray matter and white matter usually is associated with myelin, while an additional and often correlated effect is attributed to the iron concentration^4^.

A major strength of our relaxivity approach is that it captures new biological and pathological information, not acquired by traditional qMRI parameters such as R1 and R2*. We show that the great contrast between white matter and gray matter usually observed in R1 and R2* is no longer as substantial in the r1-r2* relaxivity. *In vitro*, we show that the r1-r2* relaxivity is stable across iron and liposomal concentrations. To farther demonstrate the minimal effect of myelin on the r1-r2* relaxivity in the brain, we employed the qMRI measurements of MTV^46^, MTsat^47^ and MD, which were shown to approximate the myelin content^48–53^. These *in vivo* myelin markers are all highly correlated with R1 and R2* but not with the r1-r2* relaxivity. Adapting the biophysical model for the r1-r2* relaxivity to account for myelin (Supplementary Section 4.3: Numerical simulations of the r1-r2* relaxivity.), we find that the variability in myelin within an ROI can still affect the r1-r2* relaxivity measurement. Yet, *in vivo* estimate of the myelin variability only explains ∼30% of the variation in the *in vivo* r1-r2* relaxivity across the brain. In brain tissue numerical simulations, we show that the myelin content substantially affects the measurements of R1 and R2*, but it cannot by itself explain the measured variability in the r1-r2* relaxivity across the brain (Supplementary Section 4.3: Numerical simulations of the r1-r2* relaxivity.). Nonetheless, it could be that the r1-r2* relaxivity contains some residual contribution of the myelin content. Yet, we show that this contribution is much more limited compared to traditional qMRI parameters such as R1 and R2*. In return, the novel MRI measurement of the r1-r2* relaxivity enhances the contrast between pathological tissue and normal tissue and is associated with distinct gene expression pathways. Hence, the relaxivity framework reveals distinct biological features otherwise undetectable in standard qMRI measurements.

Still, an open question remains whether the biological interpretation of the r1-r2* relaxivity demonstrated *in vitro* is also measurable *in vivo*. We bring several evidence that the r1-r2* relaxivity reveals the sensitivity of MRI to properties of the molecular iron environment, otherwise confounded by myelin and iron concentrations. First, in the healthy and aging brain we show that unlike R1 and R2*, the r1-r2* relaxivity is correlated with an iron homeostasis marker, the iron mobilization capacity (the transferrin/iron ratio). We also show that the r1-r2* relaxivity is less correlated with the absolute ferritin, transferrin or iron concentrations, indicating that it enhances the sensitivity to the homeostasis between iron compounds, rather than their absolute concentrations. Importantly, we corroborate these findings based on *in vivo* and *ex vivo* analyses of meningioma patients. We find that the variability in the *in vivo* r1-r2* relaxivity of brain tumors is explained by their transferrin-ferritin ratios and is associated with iron-related genes. These results could not be obtained for the individual measurements of R1 and R2*. To further validate that the r1-r2* relaxivity could be sensitive to the iron homeostasis, even when accounting for the massive effect of the myelin and iron concentrations on the MRI signal, we generated a simulation of a brain-like environment which contains multiple tissue components (Supplementary Section 4.3: Numerical simulations of the r1-r2* relaxivity.). We provide an example for physiological changes in the iron environment, based on the ferritin-transferrin fraction, which could lead to considerable changes in the r1-r2* relaxivity, well above the detection limit of this MRI measurement. Therefore, we show *in vitro*, *in vivo*, ex vivo and in numerical simulations, that the r1-r2* relaxivity measurement could allow to detect changes in the iron homeostasis under physiological conditions.

Our *in vitro* experiments are based on ferritin, transferrin and ferrous iron as examples for variable iron environments. Yet, in the human brain our approach is probably more broadly sensitive to the entire milieu of iron compounds, their iron binding characteristics and aggregation states. Other iron compounds that exist in the brain, such as hemoglobin, hemosiderin, neuromelanin, magnetite, ferric ion, lactoferrin and melanotransferrin ^4^, might have distinct iron relaxivities as well. Moreover, other characteristics of the iron environment such as iron compounds’ aggregate sizes, intra-aggregate spacing, spatial distributions and iron loadings, could all have an additional effect on the iron relaxivity^35^. For example, catecholamine neurons of the substantia nigra and locus coeruleus are rich in neuromelanin-iron complexes^4,66^ which could contribute to the r1-r2* relaxivity measurement in these regions (see also Supplementary Section 5: The r1-r2* relaxivity in the pallidum.). In addition, the paramagnetic properties of deoxyhemoglobin in capillaries and veins and their orientations are known to affect the R2* measurement^67–69^. Thus, the r1-r2* relaxivity might also be sensitive to the iron environment in the vascular system, which is crucial for brain iron metabolism and homeostasis^70^. Moreover, on the cellular scale there is substantial heterogeneity in the iron environment^71, 72^. R1 and R2* each represent some average of different spatial and cellular compartments, but it is unclear how the r1-r2* relaxivity is weighting such cellular contributions. Last, similar to the measurements of R1 and R2*, the r1-r2* relaxivity might be contaminated by biases related to magnetic field inhomogeneities^23^. Yet, we show that the r1-r2* relaxivity is reproducible in scan-rescan experiments (both *in vitro* and *in vivo*) and is stable across subject, thus indicating the limited effect of such biases. To conclude, while we demonstrate our approach *in vitro* based on ferritin, transferrin and ferrous iron samples, we believe that in the human brain the r1-r2* relaxivity provides more comprehensive information on the molecular iron milieu previously inaccessible by MRI.

In order to further model the separate contributions to the MRI signal of different iron compounds, it would be necessary to increase the dimensionality of the *in vivo* iron relaxivity measurement. In addition to R1 and R2*, other qMRI parameters known to be sensitive to iron include quantitative susceptibility mapping (QSM) and R2^3,4^. In addition, it was suggested that magnetization transfer (MT) measurements are affected by neuromelanin-iron complexes^12, 73^. The linear interdependencies of these other iron-related MRI measurements may uncover additional features of the iron environment^74^. Therefore, we speculate that the concept we introduce here, of exposing the iron relaxivities *in vivo* based on the linear dependency of R1 on R2* (the r1-r2* relaxivity), can be generalized to further increase MRI’s specificity for iron using additional complementary measurements. For example, in a previous work we implemented a different aspect of relaxivity for the detection of lipid composition, based on the linear dependency of qMRI parameters on the macromolecular tissue volume (MTV)^46^. Here, we demonstrate that the r1-r2* relaxivity and the dependency of R1 on MTV provide two orthogonal microstructural axes (Supplementary Section 2: The dependency of the iron relaxivity on the liposomal fraction.). The dependency of R1 on MTV changes according to the lipid types, even in the presence of iron, while the r1-r2* relaxivity is insensitive to the lipid types and provides better distinctions between iron environments. Therefore, the presented framework can be generalized further to boost MRI’s specificity and support a more comprehensive *in vivo* histology with qMRI.

While the field of *in vivo* histology with MRI is rapidly growing, ground-truth validation remains a great challenge. Here we propose a cutting-edge validation strategy combining both bottom-up and top-down approaches in which we incorporate *in vitro*, *in vivo* and ex vivo analyses. For the bottom-up analysis, we developed a unique, synthetic biological system that allows us to examine the biophysical interpretation of the r1-r2* relaxivity in highly controlled *in vitro* settings. For the top-down analysis, we tested whether our interpretation remains valid in the context of the extremely complex biological tissue. We compared the r1-r2* relaxivity measured *in vivo* to histological measurements of iron homeostasis and gene expression. This comparison was done both at the group level, based on previously reported findings, and at the single-subject level, by analyzing resected tumor tissues. To our knowledge, this is the first time that qMRI parameters measured *in vivo* have been compared to ex vivo iron histology and gene expression of the same human tissue. Taken together, the different validation strategies all indicate that the r1-r2* relaxivity increases the specificity of MRI to different molecular environments of iron, highlighting the robustness of our findings.

Our proposed approach for measuring iron homeostasis *in vivo* using the r1-r2* relaxivity may have wide clinical and scientific implications. First, the r1-r2* relaxivity provides a new contrast for imaging the brain, which is associated with 45 distinct gene sets, not associated with R1 or R2* by themselves. Moreover, we show that the r1-r2* relaxivity, which captures paramagnetic properties, enhances the contrast between tumor tissue and normal-appearing white-matter and gray-matter tissues. In agreement with these findings, meningioma tumors have been shown to contain a higher concentration of ferrimagnetic particles and an abnormal expression of iron-related genes compared to non-pathological brain tissue^17, 18^. Indeed, the contrast enhancement we saw with the r1-r2* relaxivity is similar to the one observed for Gd-enhanced imaging, which is based on the altered relaxivity in the presence of paramagnetic agents^75^. Concerns regarding the safety of Gd-based contrast agents raise the need for Gd-free diagnosis of brain tumors^55, 56^. Adjusting our approach for clinical imaging might offer safer alternatives for brain tumor diagnosis.

Finally, the sensitivity of the r1-r2* relaxivity to iron homeostasis in the brain may have clinical implications for neurodegenerative diseases. Alterations in the distribution of molecular iron compounds can lead to cellular damage which is disease-specific^10^. In particular, the iron mobilization capacity was shown to differ between elderly controls and patients diagnosed with either Parkinson’s disease (PD) or Alzheimer’s disease (AD)^7^. We found that this iron homeostasis marker is correlated with the r1-r2* relaxivity across brain regions and age groups. This result, in addition to our other validation strategies, demonstrate the sensitivity of the r1-r2* relaxivity to the iron homeostasis in the brain and in the aging process. Therefore, our approach can add a new important layer of information to existing *in vivo* PD and AD biomarkers such as neuromelanin MRI^12^ and may further advance the research, diagnosis and treatment of neurodegenerative diseases^1,2,5–9^.

## Conclusion

We present a novel MRI contrast, based on the r1-r2* relaxivity, which is sensitive to the iron homeostasis in the human brain. This new technology can differentiate between tumor tissue and non-pathological tissue without injecting contrast agents, and can detect biological properties inaccessible to conventional MRI approaches. We validated the sensitivity of the r1-r2* relaxivity to the molecular state of iron using *in vitro*, *in vivo* and ex vivo analyses. We show that our MRI technology reveals the intrinsic paramagnetic properties of different *in vitro* iron environments. Furthermore, the r1-r2* relaxivity varies with the iron mobilization capacity across brain regions and age groups, and reveals differences in iron homeostasis and iron-related gene expression in pathological tissues. Therefore, this approach may further advance our understanding of the impaired iron homeostasis in cancer, normal aging and neurodegenerative diseases, and may open new avenues for the non-invasive research and diagnosis of the living human brain.

## Methods

### In vivo iron relaxivity model

The iron relaxivity model assumes a linear relationship between relaxation rates and iron concentration^31–34^.

This linear relationship for two different iron environments *a* and *b* with iron concentrations [a] and [b] can be expressed using the following equations:

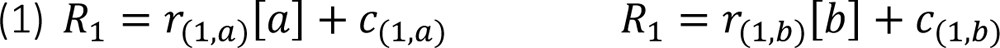

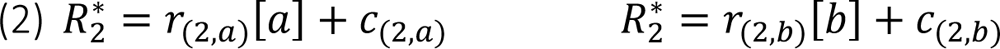

where r_(1/2,a/b)_ represents the R1-iron relaxivity or the R2*-iron relaxivity of the two iron environments, and c_(1/2,a/b)_ are constants.

The two iron environments are distinguished by their iron relaxivities under the assumption:

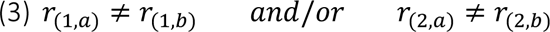

Rearranging Eq. 2:

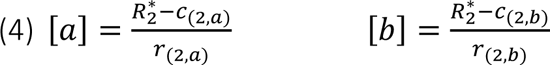

Substituting Eq. 4 in Eq. 1:

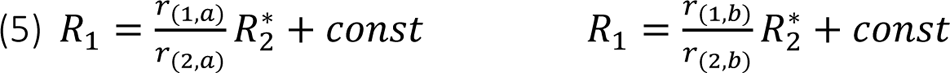

Where 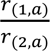 and 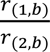 represent the linear dependencies of R1 on R2* (r1-r2* relaxivities) of the two iron environments *a* and *b*. Importantly, the MRI-measured r1-r2* relaxivity serves as an *in vivo* estimator of iron relaxivity and reveals intrinsic properties of the iron environment. Assuming that the iron relaxivity of R1 provides a different separation between the two iron environments *a* and *b* compared to the iron relaxivity of R2*:

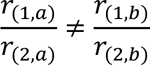

These two iron environments *a* and *b* then can be distinguished by their r1-r2* relaxivities (i.e., the *in vivo* iron relaxivity). In the brain, the r1-r2* relaxivity could be affected by the entire milieu of available iron compounds, their iron binding capacities and aggregation states (i.e. the molecular iron environment). For an elaboration of this model under the assumption of a complex iron environment and in the presence of myelin see Supplementary Section 4: Exploring the biophysical sources of the r1-r2* relaxivity.

### Phantom samples experiments

#### Phantom system

We prepared samples of four different iron compounds: transferrin (holo-transferrin human, Sigma), apo-transferrin (apo-transferrin human, Sigma), ferritin (equine spleen, Sigma), and ferrous (iron (II) sulfate heptahydrate, Sigma). These samples were prepared in three different molecular environments: liposomes, 18.2 MΩ-cm water, and bovine serum albumin (BSA, Sigma)^76, 77^. For each combination of iron compound and molecular environment, we made different samples by varying both the iron compound concentration and the lipid/BSA-water fractions. For liposomal/BSA environments, the iron compounds concentrations were divided by the water-lipid or water-BSA fractions to get units of [mg/wet ml]. The liposomes were made from a mixture of soy phosphatidylcholine (PC) and egg sphingomyelin (SM) purchased from Lipoid and used without further purification. Additional results with PC and PC-cholesterol (Sigma) liposomes are presented in Sup. Section 2. The lipid samples were mixed in chloroform at desired mole ratios and evaporated under reduced pressure (8 mbar) in a Buchi rotary evaporator vacuum system (Flawil, Switzerland). The resulting lipid film was resuspended in a 10 mM ammonium bicarbonate solution, lyophilized, and subsequently hydrated in the reassembly buffer. To achieve the desired lipid-protein concentration, the protein solution (∼50 mg/ml in water) was diluted to the right concentration and subsequently was added to the lyophilized lipid powder. For the BSA phantoms, samples were prepared by dissolving lyophilized BSA in 18.2 MΩ-cm water at the desired concentrations.

Samples were placed in a 2-ml glass vials glued to a glass box, which was then filled with ∼1% SeaKem® LE Agarose (Ornat) and 0.1% gadolinium (Gadoteric acid (Dotarem, Guerbet)) dissolved in distilled water (DW). The purpose of the agarose with gadolinium (Agar-Gd) was to stabilize the vials and to create a smooth area in the space surrounding the samples that minimalized air-sample interfaces.

### MRI acquisition for phantoms

Data were collected on a 3T Siemens MAGNETOM Skyra scanner equipped with a 32-channel head receive-only coil at the ELSC Neuroimaging Unit at the Hebrew University.

*Quantitative R1 & MTV:* 3D Spoiled gradient echo (SPGR) images were acquired with different flip angles (FA = 4°, 8°, 16°, and 30°). The TE/TR were 4.45/18 ms. The scan resolution was 0.5 mm x 0.5 mm x 0.6 mm. For calibration, we acquired an additional spin-echo inversion recovery (SEIR) scan. This scan was done on a single slice, with an adiabatic inversion pulse and inversion times of TI = 2,000, 1,200, 800, 400, and 50 ms. The TE/TR were 73/2,540 ms. The scan resolution was 1.2 mm x 1.2 mm x 2.0 mm.

*Quantitative R2*:* SPGR images were acquired with different flip angles (α = 4°, 8°, 16°, and 30°). The TR was 27 ms and 5 echoes were equally spaced between 4.45 and 20.85 ms. The scan resolution was 0.5 mm x 0.5 mm x 0.6 mm.

### Estimation of qMRI parameters for phantoms

*Quantitative R1 & MTV mapping:* R1 and MTV estimations for the lipid samples were computed with the mrQ ^46^ (https://github.com/mezera/mrQ) and Vista Lab (https://github.com/vistalab/vistasoft/wiki) software packages. The mrQ software was modified to suit the phantom system^77^. The modification utilizes the fact that the Agar-Gd mixture which fills the box around the vials is homogeneous, and therefore can be assumed to have a constant R1 value. We used this gold-standard R1 value generated from the SEIR scan to correct for the excite bias in the SPGR scans.

A mask labeling the different phantom samples was generated based on MATLAB’s “imfindcircles” function, and was filtered to remove voxels with extremely high and low signals. Voxels were filtered based on a fixed threshold on the SPGR signal at FA=16. In addition, we also filtered out those voxels in which the SPGR signal at FA=16 was two median absolute deviations away from the median value. We further edited this mask manually, removing voxels with susceptibility artifacts resulting from the vials and air pockets. To fit the R1 and proton density of each phantom sample, we calculated the median values of the SPGR signal as well as the excite and receive biases across all the voxels of each sample. These median values were used in the Vista Lab function “relaxFitT1” to find the median R1 and proton density of each sample. proton density values then were calibrated using the proton density of a water-filled vial in order to calculate the MTV values.

*Quantitative R2* mapping:* We used the SPGR scans with multiple echoes to estimate R2*. Fitting was done by taking the median values of the SPGR signal across all the voxels of the phantom sample for each TE. To label the different samples, we used the same mask that was used to calculate R1 and MTV. We then used an exponential fitting process to find R2*. As we had four SPGR scans with variable flip angles, we averaged the R2* values acquired from each of these scans for increased signal to noise ratio.

### r1-r2* relaxivity computation for phantoms

For each iron compound in each molecular environment, we computed the linear dependency of R1 on R2* across samples with varying iron-binding proteins concentrations relative to the water fractions. We fitted the following linear model across samples:

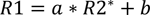

The slope of this linear model (a) represents the r1-r2* relaxivity. b is constant. This process was implemented in MATLAB.

### Estimation of total iron content in phantoms

We estimated the iron content of our transferrin and ferritin samples using the following equation:

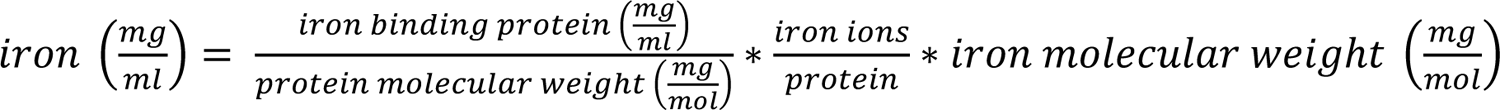

Transferrin contains 2 iron ions per protein molecule^3^, and its molecular weight was estimated as 76*10^6^ mg/mol (based on manufacturer information). The iron loading of ferritin was estimated as 2,250 iron ions per protein molecule (based on manufacturer information) and its molecular weight was estimated as 440*10^6^ mg/mol^3^. The molecular weight of iron was set to 55.847*10^3^ mg/mol.

This resulted in the following equation for converting iron-binding protein concentrations into iron concentrations:

1 mg/ml transferrin = 1.4 µg/ml iron

1 mg/ml ferritin = 0.29 mg/ml iron

### MRI human dataset

#### Healthy Human subjects

We scanned 26 young adults (aged 27 ± 10 years, 10 females), and 13 older adults (aged 70 ± 3 years, 4 females). Healthy volunteers were recruited from the community surrounding the Hebrew University of Jerusalem. The experimental procedure was approved by the Helsinki Ethics Committee of Hadassah Hospital, Jerusalem, Israel. Written informed consent was obtained from each participant prior to the procedure. This data was first used in our previous work^76^.

#### Meningioma patients

During the study period May 2019 to August 2020, we recruited 19 patients who had undergone surgery for the resection of brain meningiomas. All patients had preoperative qMRI scans in addition to their clinical brain MRI assessment. One subject, with a titanium cranial fixation plate adjacent to the tumor, was excluded from the study due to local disruption of the magnetic field. The final cohort included 18 patients (11 females). Imaging studies were anonymized before they were transferred for further analysis. Brain meningioma surgical specimens, available for 16 patients, were obtained from the fresh frozen tissue biobank of the Department of Neurosurgery, Shaare Zedek Medical Center, Jerusalem, Israel, and were transferred on dry ice for western-blot and gene expression analyses. Study participants provided informed consent according to an institutional review board.

### MRI acquisition for healthy human subjects

Data were collected on a 3T Siemens MAGNETOM Skyra scanner equipped with a 32-channel head receive-only coil at the ELSC Neuroimaging Unit at the Hebrew University.

*Quantitative R1, R2* & MTV mapping:* SPGR echo images were acquired with different flip angles (α = 4°, 10°, 20° and 30°). Each image included 5 equally spaced echoes (TE=3.34-14.02 ms) and the TR was 19 ms. The scan resolution was 1 mm isotropic. Additional SPGR echo image was acquired with an MT pulse (TE=3.34, TR=27, α = 10, 1 mm isotropic). For calibration, we acquired an additional spin-echo inversion recovery scan with an echo-planar imaging read-out (SEIR-epi). This scan was done with a slab-inversion pulse and spatial-spectral fat suppression. For SEIR-epi, the TE/TR were 49/2,920 ms. The TIs were 200, 400, 1,200, and 2,400 ms. We used 2-mm in-plane resolution with a slice thickness of 3 mm. The EPI readout was performed using 2× acceleration.

*Anatomical images:* 3D magnetization-prepared rapid gradient echo (MP-RAGE) scans were acquired for 30 of the 39 healthy subjects. The scan resolution was 1 mm isotropic, the TE/TR were 2.98/2,300 ms. Magnetization-prepared 2 rapid acquisition gradient echo (MP2RAGE) scans were acquired for the remaining 9 subjects. The scan resolution was 1 mm isotropic, the TE/TR were 2.98/5,000 ms.

*Whole-brain DTI measurements:* performed using a diffusion-weighted spin-echo EPI sequence with isotropic 1.5-mm resolution. Diffusion weighting gradients were applied at 64 directions and the strength of the diffusion weighting was set to b = 2000 s/mm2 (TE/TR=95.80/6,000 ms, G=45 mT/m, δ=32.25 ms, Δ=52.02 ms). The data includes eight non-diffusion-weighted images (*b = *0). In addition, we collected non-diffusion-weighted images with reversed phase-encode blips. For two subjects (1 young, 1 old) we failed to acquire this correction data and they were excluded from the diffusion analysis.

### MRI acquisition for meningioma patients

Data were collected on a 3T Siemens MAGNETOM Skyra scanner equipped with a 32-channel head receive-only coil at the Shaare Zedek Medical Center.

*Quantitative R1, R2* & MTV mapping:* SPGR echo images were acquired with different flip angles (α = 4°, 10°, 20° and 30°). Each image included 5 equally spaced echoes (TE=2.85-14.02 ms) and the TR was 18 ms. The scan resolution was 1.5 mm isotropic. For calibration, we acquired an additional SEIR-epi scan. This scan was done with a slab-inversion pulse and spatial-spectral fat suppression. For SEIR-epi, the TE/TR were 49/2,920 ms. The TIs were 200, 400, 1,200, and 2,400 ms. We used 2-mm in-plane resolution with a slice thickness of 3 mm. The EPI readout was performed using 2× acceleration.

*Gd-enhanced anatomical images:* Gd-enhanced MPRAGE scans were acquired. The scan resolution was 1 mm isotropic, the TE/TR were 2.4/1,800 ms. The contrast agent was either Multihance or Dotarem at a dose of 0.1 mmol/kg. Contrast agent injection and MPRAGE acquisition were done after the acquisition of the quantitative MRI protocol, or on a different day.

### Estimation of qMRI parameters for human subjects

*Quantitative R1 & MTV mapping:* Whole-brain MTV and R1 maps, together with bias correction maps of B1+ and B1-, were computed using the mrQ software^46, 78^.

*Quantitative R2* mapping:* We used the SPGR scans with multiple echoes to estimate R2*. Fitting was performed with the MPM toolbox^79^. As we had four SPGR scans with variable flip angles, we averaged the R2* maps acquired from each of these scans for increased SNR.

*Quantitative MD mapping:* Diffusion analysis was done using the FDT toolbox in FSL ^80, 81^. Susceptibility and eddy current induced distortions were corrected using the reverse phase-encode data, with the eddy and topup commands^82, 83^. MD maps were calculated using vistasoft (https://github.com/vistalab/vistasoft/wiki).

*Quantitative MTsat mapping:* MTsat maps were computed as described in Helms et al. (2008)^47^. The MTsat measurement was extracted from equation:

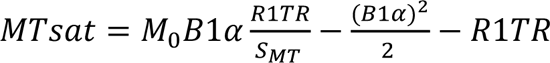

Where S_MT_ is the signal of the SPGR scan with additional MT pulse, α is the flip angle and TR is the repetition time. M_0_ (the equilibrium magnetization parameter), B1 (the transmit inhomogeneity) and R1 estimations were computed from the non-MT weighted SPGR scans, during the pipeline described under “*Quantitative R1 & MTV mapping*”. Registration of the S_MT_ image to the imaging space of the R1 map was done using a rigid-body alignment (R1, B1 and M_0_ are all in the same space).

### Brain segmentation in healthy subjects

Whole-brain segmentation was computed automatically using the FreeSurfer segmentation algorithm^84^. For subjects with MPRAGE scan, we used that as a reference; for the other subjects the MP2RAGE scan was used. These anatomical images were registered to the R1 space prior to the segmentation process, using a rigid-body alignment. FreeSurfer’s estimates of subcortical gray-matter structures were replaced with estimates from FSL’s FIRST tool^85^.

### Brain segmentation in meningioma patients

Tumor contouring was performed by the neurosurgeon (T.S.) using BrainLab’s Elements software (BrainLab AG, Munich, Germany) over the Gd-enhanced MPRAGE images, and exported as a DICOM file for further analysis. The contours of the tumors were registered to the R1 space using rigid-body segmentation. Cases that required manual adjustment were examined and approved for accuracy by the neurosurgeon (T.S.).

Whole-brain segmentation was computed automatically using the FreeSurfer segmentation algorithm^84^. We used the synthetic T1w image generated with mrQ as the reference image, from which we removed the skull and the tumor. We then ran FreeSurfer with the “-noskullstrip” flag. For each patient, we used FreeSurfer’s segmentation in the tumor-free hemisphere. Estimates for the entire white matter and gray matter tissues were averaged across the different FreeSurfer parcellations in these regions.

### r1-r2* relaxivity computation for ROIs in the human brain

We used MATLAB to compute the r1-r2* relaxivity in different brain areas. For each ROI, we extracted the R2* and R1 values from all voxels. R2* values were pooled into 36 bins spaced equally between 0 and 50 s^-1^. This was done so that the linear fit would not be heavily affected by the density of the voxels in different R2* regimes. We removed any bins in which the number of voxels was smaller than 4% of the total voxel count in the ROI. The median R2* of each bin was computed, along with the R1 median. We used these data points to fit the following linear model across bins using simple linear regression:

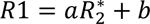

The slope of this linear model (a) represents the r1-r2* relaxivity. b is constant.

### Generating voxel-wise r1-r2* relaxivity visualizations

In order to generate a voxel-wise visualization of the r1-r2* relaxivity in the brain, we calculated the local linear dependency of R1 on R2* using a moving-window approach. For each voxel within the brain mask, we extracted R1 and R2* values of that voxel and all its neighboring voxels (a box of 125 voxels total). If at least 10 of these voxels were inside the brain mask, we fit the following linear model across these voxels:

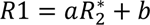

The slope of this linear model (a) represents the local r1-r2* relaxivity of the voxel. b is constant.

### Group-level comparison of qMRI parameters and histological measurements

We aggregated data published in different papers^5,7,9^ that describe ferritin, transferrin and iron concentrations in 11 brain regions. One of the papers^9^ described the concentration of L-rich ferritin and H-rich ferritin independently and we combined these estimates for each ROI to get the total ferritin concentration. One of the papers^5^ reported the iron level in units of [µmol Fe/ g protein] and we converted these measurements to units of [µg Fe/ g protein]. In order to use this data for our analysis with we matched the brain regions reported in the literature with their corresponding FreeSurfer labels (see table). The table summarizes the aggregated data:

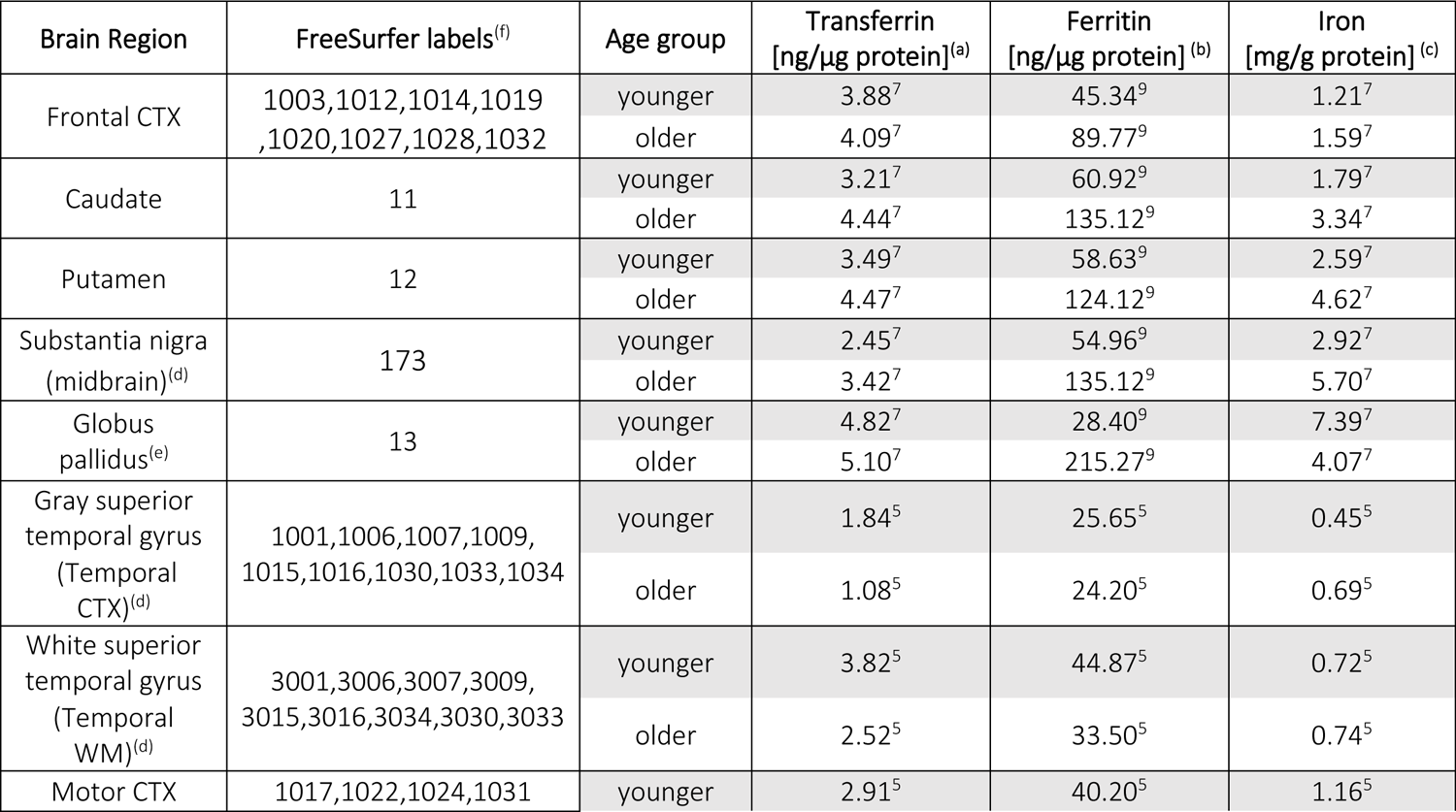
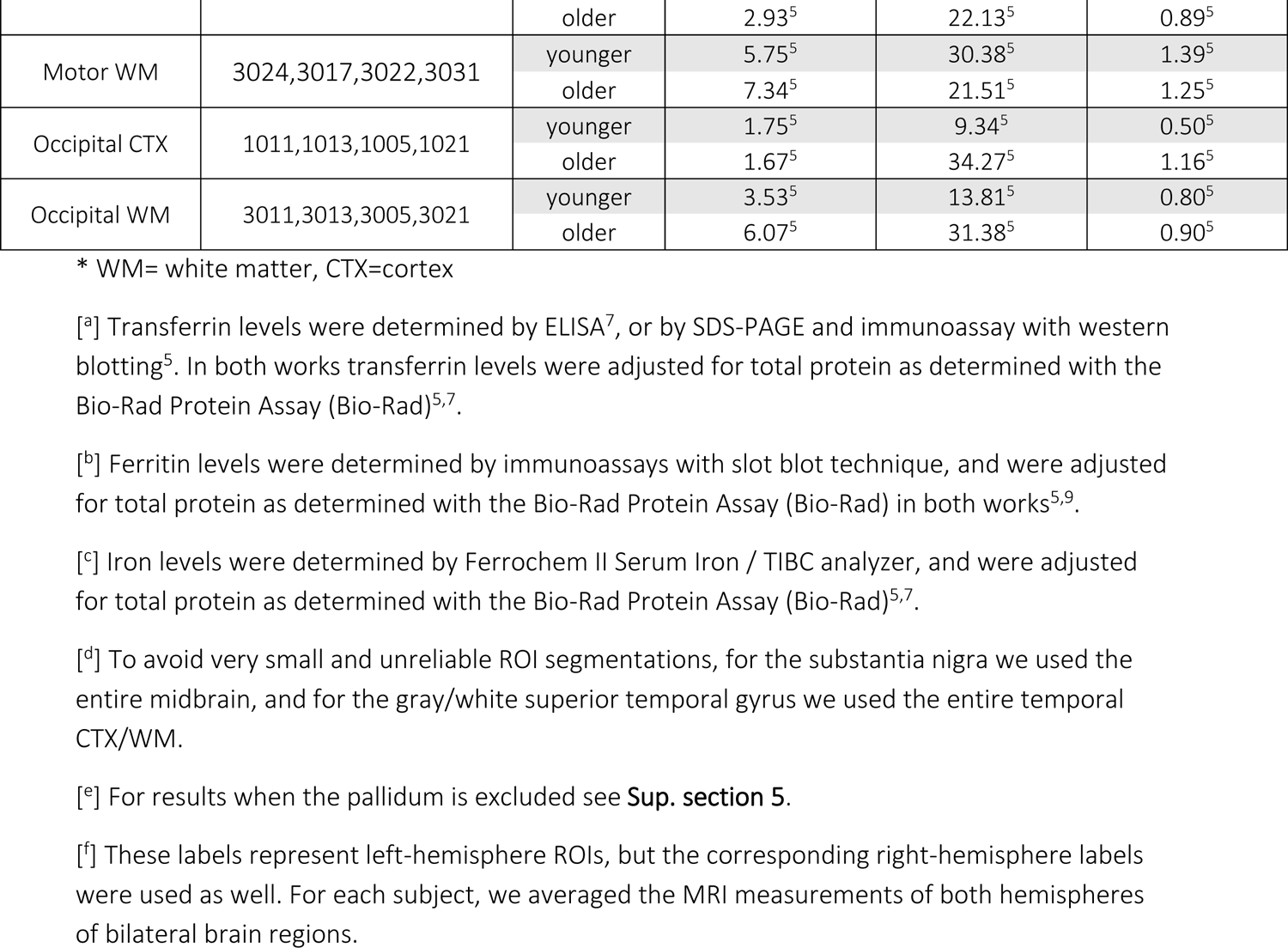

| Brain Region | FreeSurfer labels<sup>(f)</sup> | Age group | Transferrin<br>[ng/$\mu\text{g}$ protein]<sup>(a)</sup> | Ferritin<br>[ng/$\mu\text{g}$ protein]<sup>(b)</sup> | Iron<br>[mg/g protein]<sup>(c)</sup> |
| --- | --- | --- | --- | --- | --- |
| Frontal CTX | 1003,1012,1014,1019<br>,1020,1027,1028,1032 | younger | 3.88<sup>7</sup> | 45.34<sup>9</sup> | 1.21<sup>7</sup> |
|  |  | older | 4.09<sup>7</sup> | 89.77<sup>9</sup> | 1.59<sup>7</sup> |
| Caudate | 11 | younger | 3.21<sup>7</sup> | 60.92<sup>9</sup> | 1.79<sup>7</sup> |
|  |  | older | 4.44<sup>7</sup> | 135.12<sup>9</sup> | 3.34<sup>7</sup> |
| Putamen | 12 | younger | 3.49<sup>7</sup> | 58.63<sup>9</sup> | 2.59<sup>7</sup> |
|  |  | older | 4.47<sup>7</sup> | 124.12<sup>9</sup> | 4.62<sup>7</sup> |
| Substantia nigra<br>(midbrain)<sup>(d)</sup> | 173 | younger | 2.45<sup>7</sup> | 54.96<sup>9</sup> | 2.92<sup>7</sup> |
|  |  | older | 3.42<sup>7</sup> | 135.12<sup>9</sup> | 5.70<sup>7</sup> |
| Globus<br>pallidus<sup>(e)</sup> | 13 | younger | 4.82<sup>7</sup> | 28.40<sup>9</sup> | 7.39<sup>7</sup> |
|  |  | older | 5.10<sup>7</sup> | 215.27<sup>9</sup> | 4.07<sup>7</sup> |
| Gray superior<br>temporal gyrus<br>(Temporal<br>CTX)<sup>(d)</sup> | 1001,1006,1007,1009,<br>1015,1016,1030,1033,1034 | younger | 1.84<sup>5</sup> | 25.65<sup>5</sup> | 0.45<sup>5</sup> |
|  |  | older | 1.08<sup>5</sup> | 24.20<sup>5</sup> | 0.69<sup>5</sup> |
| White superior<br>temporal gyrus<br>(Temporal<br>WM)<sup>(d)</sup> | 3001,3006,3007,3009,<br>3015,3016,3034,3030,3033 | younger | 3.82<sup>5</sup> | 44.87<sup>5</sup> | 0.72<sup>5</sup> |
|  |  | older | 2.52<sup>5</sup> | 33.50<sup>5</sup> | 0.74<sup>5</sup> |
| Motor CTX | 1017,1022,1024,1031 | younger | 2.91<sup>5</sup> | 40.20<sup>5</sup> | 1.16<sup>5</sup> |
|  |  | older | 2.93<sup>5</sup> | 22.13<sup>5</sup> | 0.89<sup>5</sup> |
| Motor WM | 3024,3017,3022,3031 | younger | 5.75<sup>5</sup> | 30.38<sup>5</sup> | 1.39<sup>5</sup> |
|  |  | older | 7.34<sup>5</sup> | 21.51<sup>5</sup> | 1.25<sup>5</sup> |
| Occipital CTX | 1011,1013,1005,1021 | younger | 1.75<sup>5</sup> | 9.34<sup>5</sup> | 0.50<sup>5</sup> |
|  |  | older | 1.67<sup>5</sup> | 34.27<sup>5</sup> | 1.16<sup>5</sup> |
| Occipital WM | 3011,3013,3005,3021 | younger | 3.53<sup>5</sup> | 13.81<sup>5</sup> | 0.80<sup>5</sup> |
|  |  | older | 6.07<sup>5</sup> | 31.38<sup>5</sup> | 0.90<sup>5</sup> |

\* WM= white matter, CTX=cortex
[<sup>a</sup>] Transferrin levels were determined by ELISA<sup>7</sup>, or by SDS-PAGE and immunoassay with western blotting<sup>5</sup>. In both works transferrin levels were adjusted for total protein as determined with the Bio-Rad Protein Assay (Bio-Rad)<sup>5,7</sup>.
[<sup>b</sup>] Ferritin levels were determined by immunoassays with slot blot technique, and were adjusted for total protein as determined with the Bio-Rad Protein Assay (Bio-Rad) in both works<sup>5,9</sup>.
[<sup>c</sup>] Iron levels were determined by Ferrochem II Serum Iron / TIBC analyzer, and were adjusted for total protein as determined with the Bio-Rad Protein Assay (Bio-Rad)<sup>5,7</sup>.
[<sup>d</sup>] To avoid very small and unreliable ROI segmentations, for the substantia nigra we used the entire midbrain, and for the gray/white superior temporal gyrus we used the entire temporal CTX/WM.
[<sup>e</sup>] For results when the pallidum is excluded see **Sup. section 5**.
[<sup>f</sup>] These labels represent left-hemisphere ROIs, but the corresponding right-hemisphere labels were used as well. For each subject, we averaged the MRI measurements of both hemispheres of bilateral brain regions.

### Western blot analysis of meningioma tissue

Fresh frozen meningioma samples (40-50 mg) from 16 patients were homogenized in 200 µL of RIPA buffer (Thermo Fisher Scientific) supplemented with protease inhibitor (Sigma-Aldrich,) using a Bioruptor Pico sonication device (Diagenode) and protein extraction beads (Diagenode, NJ, USA) according to the manufacturer instructions. Protein concentration was determined using the Pierce assay (Thermo Fisher Scientific, MA, USA). Samples containing 20 µg of protein were separated on 4-20% Tris-Glycine SDS-PAGE gel (Bio-Rad, CA, USA) and transferred to PVDF membrane using Trans-Blot Turbo transfer system and transfer packs (Bio-Rad, Hercules, CA, USA). Membranes were probed using Anti-Ferritin Light chain (#AB69090, Abcam, 1:1,000 dilution) and Anti-Transferrin (#AB82411, Abcam, 1:10,000 dilution) primary antibodies and appropriate horseradish peroxidase-conjugated secondary antibody (Abcam, Cambridge, UK). Membranes were treated with EZ-ECL (Biological industries, Beit-Ha’emek, Israel) and visualized using ImageQuant LAS 4000 (GE Healthcare, IL, USA). Blot intensities were quantified using the FIJI ImageJ software^86^. The ratio of transferrin/ferritin was based on the ratio in the blot intensities of transferrin and ferritin. Due to the noisy nature of the western-blot analysis, we averaged the estimates over six repetitions. We then used the median transferrin/ferritin ratio across subjects (which was equal to 1) as the threshold between the two groups (low and high transferrin/ferritin ratio).

### RNA-sequencing of meningioma tissue

RNA-seq libraries: Tumor samples from 17 patients (samples from 16 patients and a replicate for one) were flash frozen and kept in −80c until processing. RNA isolation was done with the following steps: First, frozen tissue was chopped and transferred with a 2ml lysis buffer (Macherey-Nagel, 740955) five times through a needle attached to a 0.9 mm syringe to achieve homogenization. Next, total RNA was extracted with NucleoSpin RNA kit (Macherey-Nagel, 740955), following the standard protocol. Finally, mRNA was isolated using the NEBNext Poly(A) mRNA Magnetic Isolation Module (NEB E7490S), using 5ug of total RNA as an input and following the standard protocol. The purified mRNA was used as input for cDNA library preparation, using NEBNext® Ultra™ II Directional RNA Library Prep Kit for Illumina (NEB E7760), and following the standard protocol. Quantification of the libraries was done by Qubit and TapeStation. Paired-end sequencing of the libraries was performed on Nextseq 550.

Data processing: The demultiplexing of the samples was done with Illumina’s bcl2fastq software. The fastq files were next aligned to the human genome (hg38) using STAR and the transcriptome alignment and gene counts were obtained with HTseq. For quality control RNAseQC software was used. Quality control and data normalization was done in R using the DEseq2 package from Bioconductor (version 3.13). The counts matrix per gene and sample were normalized using the Variance stabilizing transformation. Genes with less than 5 counts were filtered out of the analysis. The filtered and normalized matrix was used in all downstream analysis.

### Gene set enrichment analysis (GSEA)

The final sequencing dataset included the expression of approximately 27,000 genes in 17 tumor samples. We then excluded unannotated genes based on the gene ontology resource (http://geneontology.org/) as well as genes with low (<6) expression levels, yielding 19,500 genes.

We used GSEA to further validate that the subset of highly correlated genes is not random, but rather represents known biological pathways. For this aim, we calculated the correlations across patients between the expression of each of the genes and one of the qMRI parameters (R1, R2* or r1-r2* relaxivity). For each of the qMRI parameters, genes were ranked based on the *r* values of the correlations, and the ranked list was used in the GSEAPranked toolbox^58, 59^. The gene sets databases used for this analysis included go, biocarta, kegg, pid, reactome and wikipathways. The primary result of the GSEA is the enrichment score (ES), which reflects the degree to which a gene set is overrepresented at the top or bottom of a ranked list of genes: A positive ES indicates gene set enrichment at the top of the list, while a negative ES indicates gene set enrichment at the bottom. The normalized ES (NES) accounts for differences in gene set size and in correlations between gene sets and the expression dataset.

One tumor sample was excluded from the analysis, as the R1 and R2* values in the tumor were relatively high, which led to the fact that no significantly enriched pathway were found for R1 and R2* (though we did detect significantly enriched pathways for the r1-r2* relaxivity). Removing this outlier improved GSEA results for R1 and R2* and we therefore excluded this subject.

Following the GSEA analysis, we found a total of 101 significantly enriched pathways for at least one of the R1, R2* and r1-r2* relaxivity. We then clustered those significantly enriched pathways using the “clustergram” function in MATLAB. In order to evaluate which genes are included within the top enrichment pathways for each MRI parameter, we used Leading Edge Analysis (as implemented in the GSEA toolbox).

## Data availability

The data that support the findings of this study are available on request from the corresponding author (S.F.). The data are not publicly available due to them containing information that could compromise research participant privacy/consent.

## Code availability

A toolbox for computing the r1-r2* relaxivity, including example data, is available at: [https://github.com/shirfilo/r1_r2s_rel_toolbox].

## Supplementary Materials

**Sup. Figure 1:**
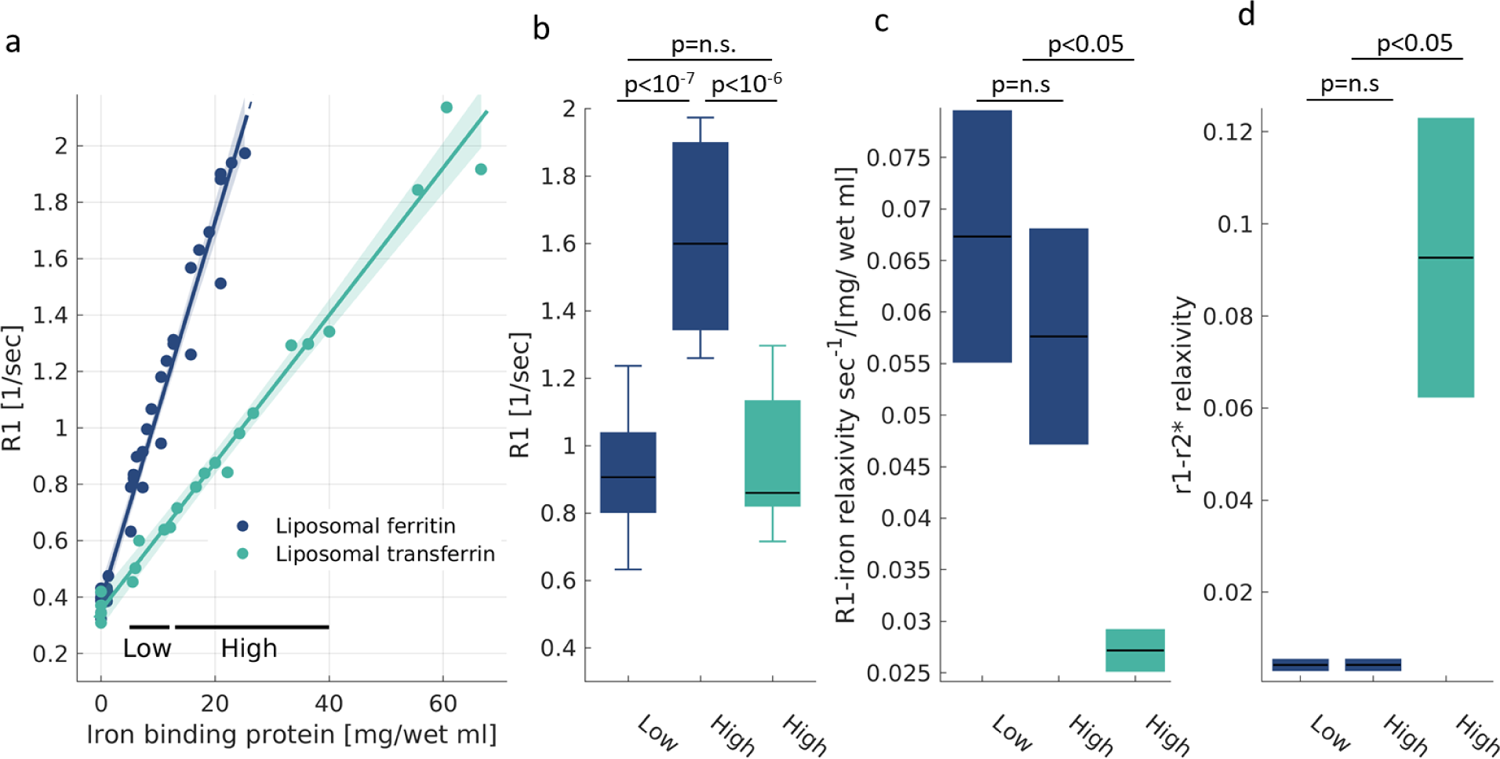
The effect of iron concentration on different MR estimations. **(a)** The dependency of R1 on the iron-binding protein concentration for liposomal ferritin and liposomal transferrin. Data points represent liposomal samples with varying iron-binding protein concentrations relative to the water fraction ([mg/wet ml]). The linear relationships between relaxation rates and iron-binding protein concentrations are marked by lines. The slopes of these lines are the iron relaxivities. Shaded areas represent the 95% confidence bounds. **(b)** The ambiguity in R1; R1 changes as a function of both iron environment and iron concentration. This is shown by calculating the median R1 value over samples with high and low iron-binding protein concentrations (marked in (a); concentration ranges were chosen so that the number of data points in each range is similar). We find that R1 is greater for a higher ferritin concentration than for a lower ferritin concentration, but also find that R1 is greater for ferritin than for transferrin. For each box, the central line marks the median, the box extends vertically between the 25th and 75th percentiles, and the whiskers extend to the most extreme data points. **(c-d)** The ambiguity in R1 is resolved by the R1-iron relaxivity (c) and the r1-r2* (d), which are consistent when computed over higher or lower ferritin concentrations, and are consistently different from the iron relaxivity of transferrin regardless the concentration. For each box, the central lines marks the iron relaxivity, and the box shows the 95% confidence bounds of the linear fit. p-values are for the ANCOVA test corrected for multiple comparisons.

## Supplementary Section 1: The dependency of R1 and R2* on the iron concentration

In *Figure 1* we computed the iron relaxivity as the dependency of R1 and R2* on the concentration of iron-binding proteins. We showed that different iron environments have different relaxivities. However, different proteins bind different amounts of iron. For example, ferritin binds three orders of magnitude more iron ions than does transferrin^3^. Therefore, we wanted to exclude the possibility that the different iron ion concentrations drive the different relaxivities of ferritin and transferrin.

**Sup. Figure 2:**
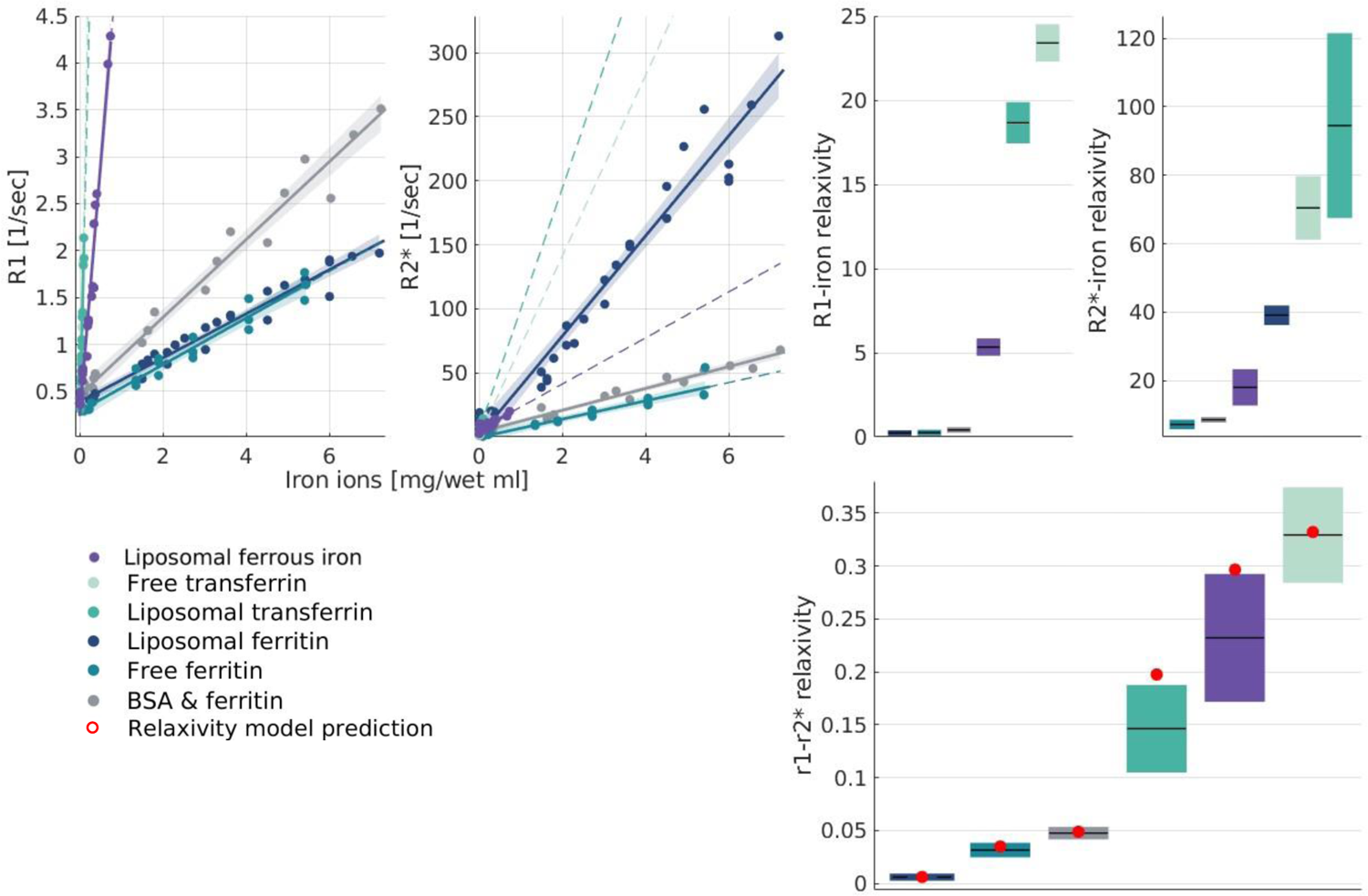
The iron relaxivity and the r1-r2* relaxivity are sensitive to the molecular type of iron regardless of the differences in iron-binding. (a-b) The dependency of R1 and R2* on the estimated iron concentration (see method section “Estimation of total iron content in phantoms”) for six different iron compounds: free ferritin, liposomal-ferritin, Bovine Serum Albumin (BSA)-ferritin mixture, free transferrin, liposomal transferrin and liposomal ferrous iron. Data points represent samples with varied estimated iron ion concentrations relative to the water fraction ([mg/wet ml]). The linear relationships between relaxation rates and iron concentration are marked by lines. The slopes of these lines are the iron relaxivities. Dashed lines represent extrapolations the linear fits, and shaded areas represent the 95% confidence bounds. (c) The iron relaxivities of R1 and R2* are different for different iron environments (p(ANCOVA)<10^39^). Iron relaxivity is calculated here based on the estimated iron concentration (and not iron-binding proteins concentrations, as in Figure 1). To do so, we use the slope of the linear relationships shown in (a,b), expressed in [sec-1/(mg/wet ml)]. For each box the central lines marks the iron relaxivity, and the box shows the 95% confidence bounds of the linear fit. (d) The theoretical model successfully predicts the r1-r2* relaxivity even when it is based on the estimated iron ions concentration (and not iron compound concentrations, as in Figure 1). The model’s prediction is based on the ratio between the iron relaxivities of R1 and R2* as shown in (c). For each box the central line marks the r1-r2* relaxivity, and the box shows the 95% confidence bounds of the linear fit. Red dots represent the prediction of the theoretical model.

**Sup. Figure 3:**
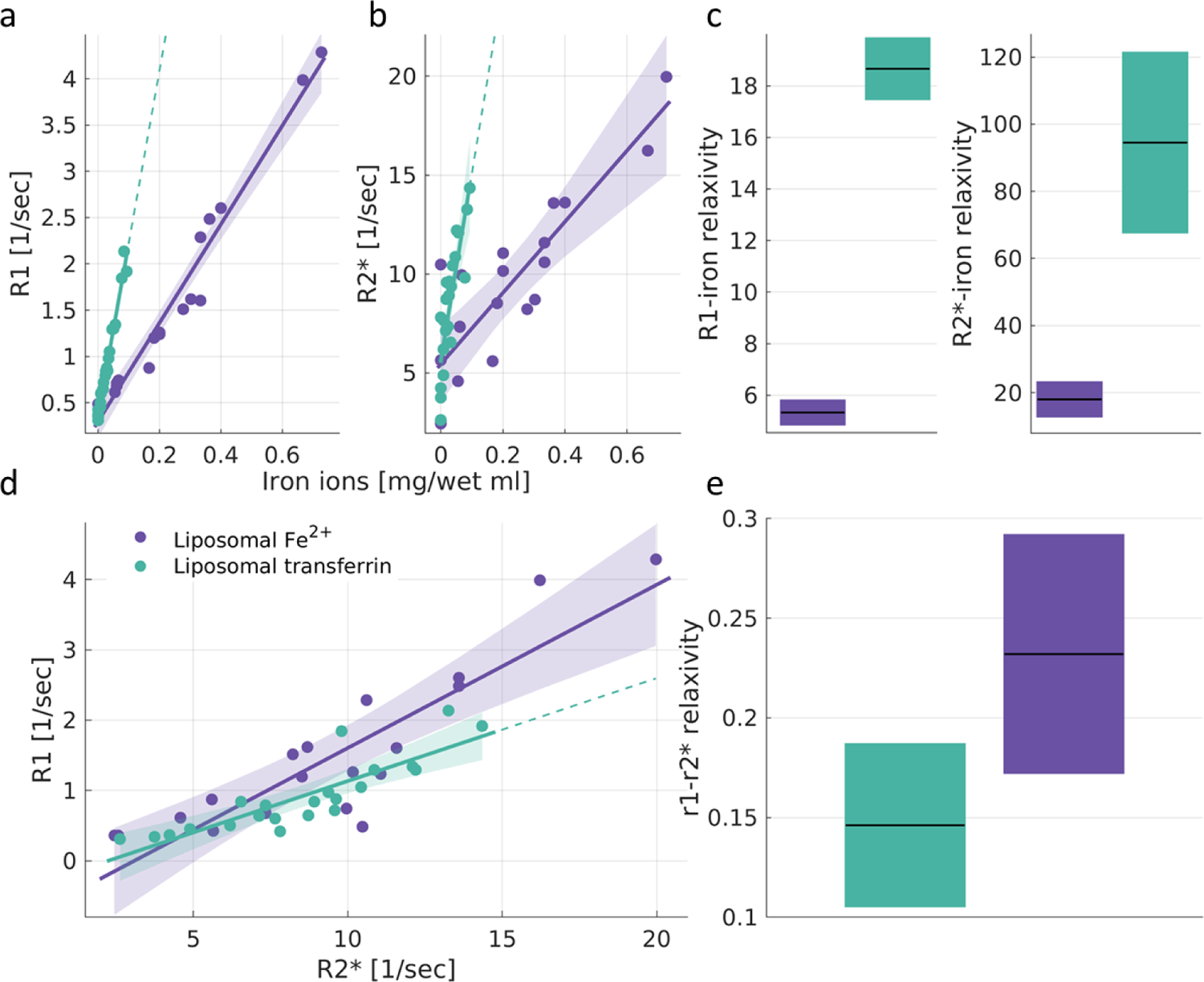
The iron relaxivity and the r1-r2* relaxivity are sensitive to the molecular iron environment even when the iron concentration is similar. (a-b) The dependency of R1 and R2* on the estimated iron concentration for two different iron environments: liposomal transferrin (purple) and liposomal Fe^2+^ (green). Data points represent liposomal samples with varying iron ion concentrations relative to the water fraction ([mg/wet ml]). The linear relationships between relaxation rates and iron ion concentration are marked by lines. The slopes of these lines are the iron relaxivities. Dashed lines represent extrapolations of the linear fits. Shaded areas represent the 95% confidence bounds. (c) The iron relaxivity of R1 and R2* is different for different iron environments (p(ANCOVA)<10^-4^). Iron relaxivity is calculated by taking the slope of the linear relationships shown in (a,b), and is measured in [sec-1/(mg/wet ml)]. For each box, the central line marks the iron relaxivity, and the box shows the 95% confidence bounds of the linear fit. (d) The dependency of R1 on R2* for different iron environments. Data points represent samples with varying concentrations. The linear relationships between R1 and R2* are marked by lines. The slopes of these lines are the r1-r2* relaxivities. Dashed lines represent extrapolations of the linear fits. Shaded areas represent the 95% confidence bounds. (e) The r1-r2* relaxivity is different for different iron environments (p(ANCOVA)<0.05). For each box, the central line marks the r1-r2* relaxivity, and the box shows the 95% confidence bounds of the linear fit.

**Sup. Figure 4:**
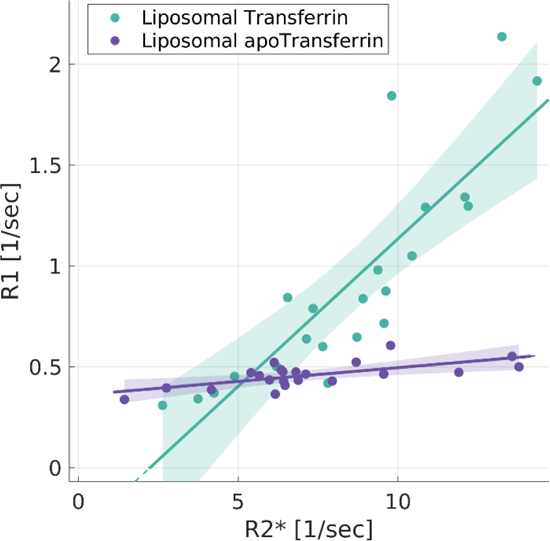
Validating the sensitivity of the the r1-r2* relaxivity to the paramagnetic properties of transferrin. Data points represent liposomal samples with varying concentrations of transferrin (green) and apo-transferrin (transferrin which is not bound to iron, in purple). The linear relationships between R1 and R2* are marked by lines. The slopes of these lines are the r1-r2* relaxivities. Shaded areas represent the 95% confidence bounds. Apo-transferrin with no iron has lower r1-r2* relaxivity compared to iron-bound transferrin (p(ANCOVA)<10^-8^). Therefore, the r1-r2* relaxivity is sensitive to the paramagnetic properties of iron-binding proteins and not to the proteins themselves.

We verified that the relaxivity changes according to the molecular type of iron, even when accounting for discrepancies in iron loading. We estimated the iron ion concentrations for ferritin and transferrin (see Methods section “Estimation of total iron content in phantoms”) and tested whether those values can explain their different iron relaxivities. Importantly, after computing the iron relaxivity as the dependency of relaxation rates on the iron ion concentrations (rather than the concentration of iron-binding proteins), we still find that different iron environments have distinct relaxivities (**Sup. Figure 2**, p(ANCOVA)<10^-39^).

To further stress the sensitivity of the r1-r2* relaxivity to the iron environment, we compared the relaxivity of liposomal ferrous iron (Fe^2+^) and iron bound to liposomal transferrin (**Sup. Figure 3**). Unlike ferritin and transferrin, these two iron compounds have relatively similar iron ion concentrations. Yet we find that they produce different iron relaxivities (p(ANCOVA)<10^-4^). The r1-r2* relaxivities of these two iron environments are different as well (p(ANCOVA)<0.05). Therefore, the iron relaxivity and the r1-r2* relaxivity are changing as a function of the molecular iron environment, even when accounting for the differences in iron binding.

## Supplementary Section 2: The dependency of the iron relaxivity on the liposomal fraction

R1 and R2* measured in the brain are known to be sensitive to myelin content^4,19, 25, 29, 40–42^. Myelin is composed mainly of lipids, though it also includes proteins. We tested the effect of the myelin fraction on iron relaxivity by varying the liposomal and protein (BSA) fractions in our phantoms.

In histological studies of brain iron, the iron concentrations often are reported relative to the wet weight, as this is considered more accurate^4^. To match our *in vitro* analysis to brain histology as much as possible, we calculated the iron-binding proteins’ concentrations relative to the water fraction ([mg/wet ml]). This was done by computing the ratio between the iron concentration and the water fraction (which is complementary to the liposomal or protein fractions). The iron relaxivities shown in **Figure 1** were therefore calculated as the linear dependencies of relaxation rates on the iron-binding protein concentration relative to the water fraction. **Sup. Figure 5** presents the effect of the variable liposomal (or BSA) fractions on the iron relaxivity and on the r1-r2* relaxivity. When iron-binding protein concentrations were not calibrated to the water fraction (units of [mg/ml]), some variability in R1 and R2* values for different liposomal (or BSA) fractions was observed (**Sup. Figure 5a,c**). However, the iron relaxivities of different iron environments were still distinct, despite the liposomal (or BSA) fractions’ variability. Calibrating the iron-binding protein concentrations to the water fraction (units of [mg/wet ml]) further eliminated the effect of the variable liposomal (or BSA) fractions on the iron relaxivities (**Sup. Figure 5b,d**). This is evident by the alignment of the data points with different liposomal (or BSA) fractions along the iron relaxivity’s linear fits. Therefore, while the non-water fraction has an effect on the relaxation rates, it does not disrupt the sensitivity of the iron relaxivities to the molecular type of iron.

We further estimated the effect of the liposomal (or BSA) fractions on the r1-r2* relaxivity. In **Sup. Figure 5e** we show the same r1-r2* relaxivities presented in **Figure 1**, but now the liposomal (or BSA) fractions are indicated by different symbols. Similarly to the iron relaxivities, the r1-r2* relaxivities of different iron environments were distinct, even though they were calculated across varying liposomal (or BSA) fractions. Moreover, we estimated the r1-r2* relaxivity separately for each liposomal (or BSA) fraction (**Sup. Figure 5f**). We find that the r1-r2* relaxivity differences between iron compounds are greater than the differences within each iron compound for the variable liposomal (or BSA) fractions.

The dependency of R1 on the macromolecular tissue volume (MTV) was associated with lipid composition in our previous work^76^. We tested this finding in the presence of iron by calculating the R1-MTV dependencies for different types of lipids mixed with iron (**Sup. Figure 6a-b**). Notably, in the current study we sampled only three liposomal fractions, and therefore the variation in the iron concentration between the samples was much richer than the variation in lipid concentration. Still, we were able to replicate our finding regarding the sensitivity of the R1-MTV dependency to lipid type. We find that the R1-MTV dependencies are different for two types of lipid mixtures (phosphatidylcholine (PC) and phosphatidylcholine-sphingomyelin (PC-SM)) mixed with ferrous (Fe^2+^) iron (**Sup. Figure 6a**). In the presence of ferritin, the difference between the R1-MTV dependencies of the two lipids is smaller (**Sup. Figure 6b**).

Unlike the R1-MTV dependency, the r1-r2* relaxivity is insensitive to the lipid composition (**Sup. Figure 6c**): different lipids mixed with ferritin have a similar r1-r2* relaxivity (p(ANCOVA)=0.11). The variability in the r1-r2* relaxivity was much bigger when comparing these different liposomal ferritin samples to liposomal transferrin (p(ANOCVA)<10^-7^). Compared to the R1-MTV dependencies, we find that the r1-r2* relaxivity provides a better distinction between iron compounds. **Sup. Figure 7** presents the r1-r2* relaxivities and the R1-MTV dependencies for different iron compounds. ANCOVA tests for the R1-MTV dependencies reveal that the only significant distinction is between the BSA-ferritin mixture and all the liposomal iron compounds (p(ANCOVA)<10^-5^). The rest of the iron environments are indistinguishable in terms of their R1-MTV dependencies. On the contrary, all iron environments were distinguishable in terms of their r1-r2* relaxivity (p(ANCOA)<10^-32^).

**Sup. Figure 5:**
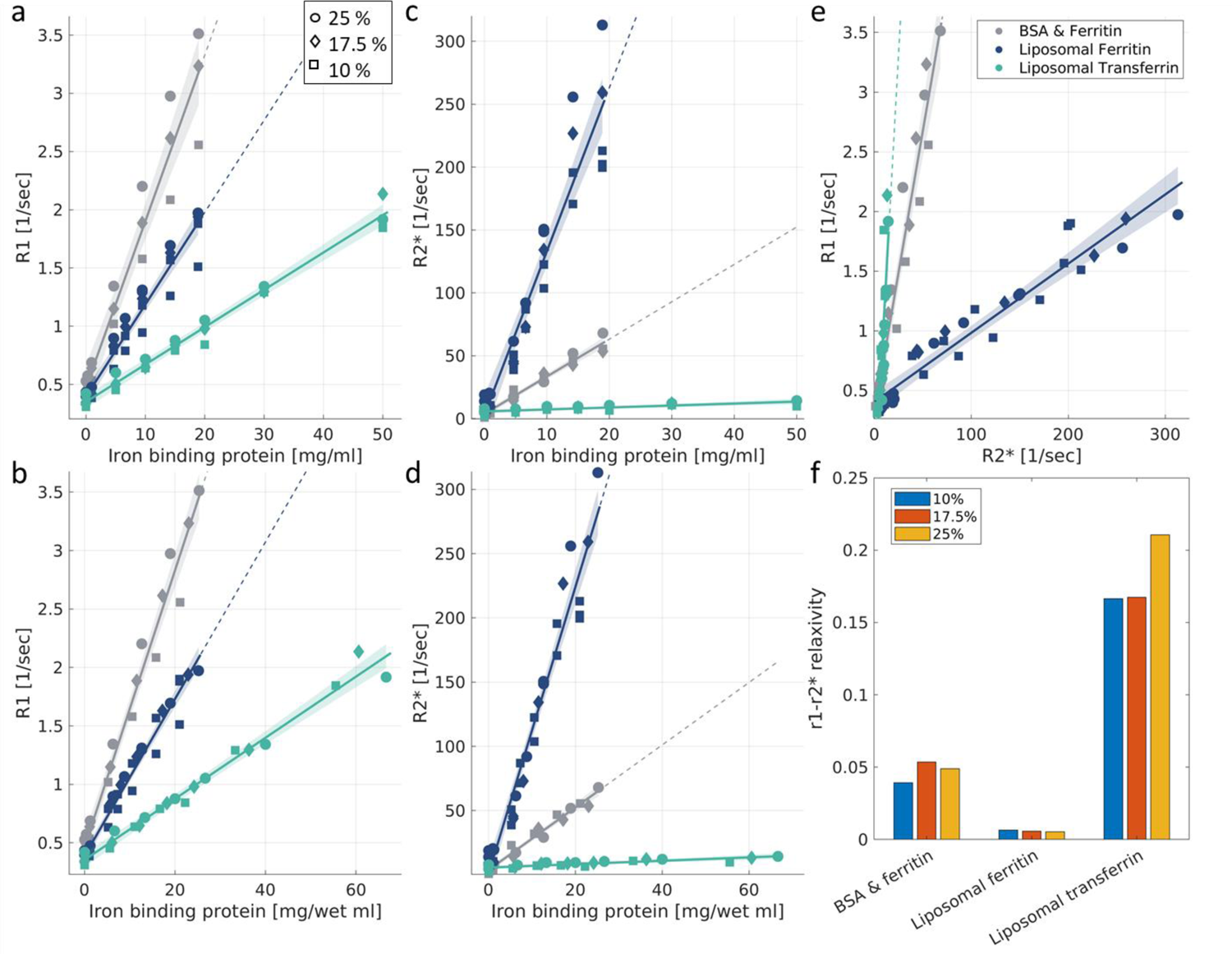
Relaxivities are stable across liposomal or BSA fractions. **(a)** The dependency of R1 on the iron-binding protein concentration for different liposomal (or BSA) fractions (different symbols) and different iron environments (different colors). The x-axis represents the absolute concentration of iron-binding proteins (not relative to the water concentration, as in b). The linear relationships between relaxation rates and iron-binding protein concentration are marked by lines. The slopes of these lines are defined as the iron relaxivities. R1 values are affected by the variable liposomal (or BSA) fractions, but the iron relaxivities of different iron environments are still distinct, regardless of this manipulation. Dashed lines represent extrapolations of the linear fits. Shaded areas represent the 95% confidence bounds. **(b)** The dependency of R1 on the iron-binding protein concentration for different liposomal (or BSA) fractions (different symbols) and different iron environments (different colors). Here the x-axis represents the concentration of iron-binding proteins relative to the water fraction (which varies with the liposomal or BSA fraction). This estimation, in units of [mg/wet ml], further eliminates the effect of the liposomal (or BSA) fraction on the iron relaxivities. This is evident by the alignment of the data points with different liposomal (or BSA) fractions (different symbols) along the iron relaxivity linear fit. **(c-d)** A similar analysis for the R2*-iron relaxivity. The effect of the different liposomal (or BSA) fractions on the R2*-iron relaxivity is eliminated by the calculation of the iron-binding proteins concentration relative to the water fraction ([mg/wet ml]). **(e)** The dependency of R1 on R2* for different liposomal (or BSA) fractions (different symbols) and different iron environments (different colors). The r1-r2* relaxivities of different iron environments are distinct even when calculated across liposomal (or BSA) fractions**. (f)** The r1-r2* relaxivity (y-axis) for different compounds of iron (liposomal ferritin, liposomal transferrin and BSA-ferritin mixture) in three different liposomal (or BSA) fractions (colors). The differences in the r1-r2* relaxivity between iron environments are greater than the differences within each iron environment for the variable liposomal (or BSA) fractions.

**Sup. Figure 6:**
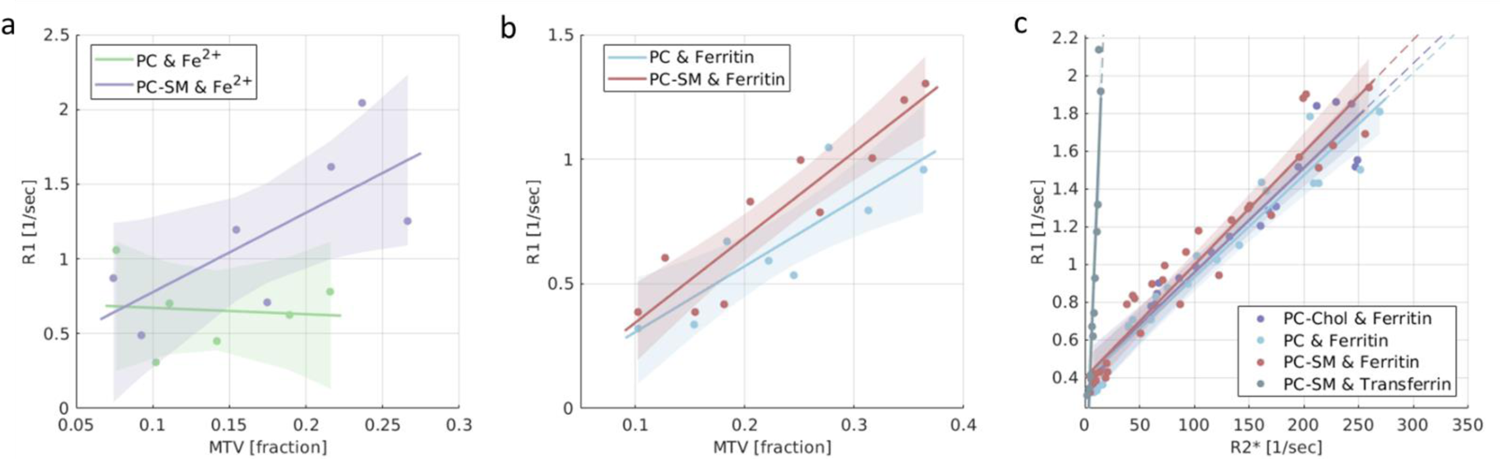
The r1-r2* relaxivity is stable for different types of lipids, while the R1-MTV dependency is sensitive to the lipid type. (a) The dependency of R1 on MTV for an iron ion compound (Fe^2+^) mixed with two different lipids: phosphatidylcholine (PC, green) and a mixture of PC-sphingomyelin (PC-SM, blue). This result replicates the sensitivity of the MTV dependencies to lipid types^76^ in Fe^2+^-containing phantoms. (b) The dependency of R1 on MTV for a second iron compound (ferritin) mixed with the same two lipids (PC and PC-SM). (c) The dependency of R1 on R2* (r1-r2* relaxivity) for four different iron-lipid mixtures: ferritin-PC ferritin-PC-SM, transferrin-PC-SM and ferritin-PC-cholesterol (PC-Chol, blue). The r1-r2* relaxivity is similar for the different lipid types mixed with ferritin, and the main difference is between the iron binding proteins; i.e., transferrin sample and the ferritin samples.

**Sup. Figure 7:**
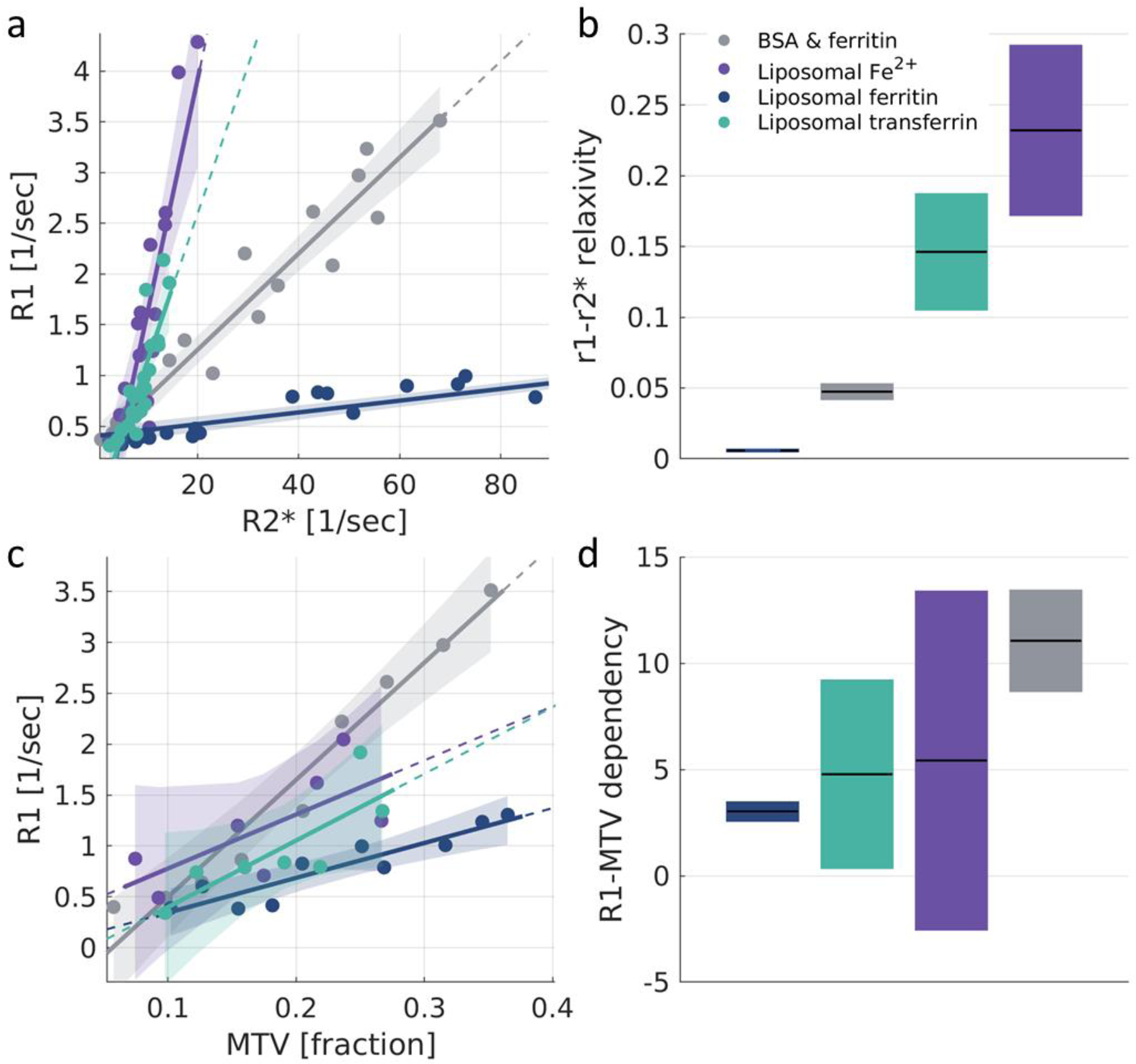
Iron environments are less distinguishable with MTV dependencies than with the r1-r2* relaxivity. (a) The dependency of R1 on R2* (r1-r2* relaxivity) for different iron environments: liposomal-ferritin, BSA-ferritin mixture, liposomal transferrin and liposomal Fe^2+^. Liposomal samples are based on PC-sphingomyelin. Data points represent samples with varying iron compounds concentrations relative to the water fraction. The linear relationships between relaxation rates are marked by lines, whose slopes represent the r1-r2* relaxivities. Dashed lines represent extrapolations of the linear fits. Shaded areas represent the 95% confidence bounds. The x-axis presents only partial range of R2* values, similar to *Figure 1d* (for the entire R2* range, see the inset of *Figure 1d*). (b) The r1-r2* relaxivities are different for these four iron environments. For each box, the central line marks the r1-r2* relaxivity, and the box shows the 95% confidence bounds of the linear fit. (c) The dependency of R1 on MTV for these four iron environments. Data points represent samples with varying iron compounds concentrations relative to the water fraction. The linear relationships between R1 and MTV are marked by lines, whose slopes represent the R1-MTV dependencies. Dashed lines represent extrapolations of the linear fits. Shaded areas represent the 95% confidence bounds. (d) The R1-MTV dependencies for the four iron environments. For each box, the central line marks the R1-MTV dependency, and the box showes the 95% confidence bounds of the linear fit.

**Sup. Figure 8:**
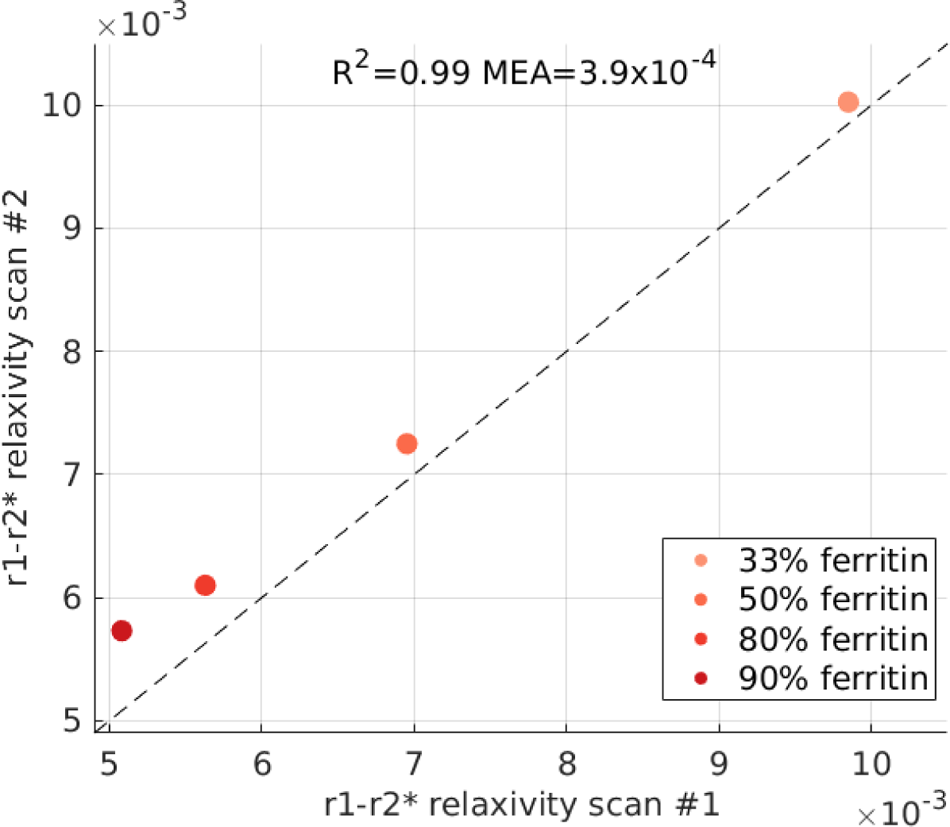
The reproducibility of the r1-r2* relaxivity measurement in vitro. The reproducibility of the r1-r2* relaxivity measurement for in vitro ferritin-transferrin mixtures was estimated based on scan-rescan experiments. Four different transferrin-ferritin mixtures were scanned twice (on different days). Each mixture experiment had a different transferrin-ferritin fraction (different colors, legend shows the percentage of ferritin in the mixture). The r1-r2* relaxivity of each mixture experiment was calculated over samples with the same transferrin-ferritin fraction but varying total iron-binding protein concentrations. Figure shows the r1-r2* relaxivity values measured in the first scan (x-axis) vs. the r1-r2* relaxivity values measured in the second scan (y-axis) for each in vitro experiment. Dashed line is the identity line. The measured scan-rescan mean absolute error (MAE) represents an experimental estimate of the detection limit of the r1-r2* relaxivity.

**Sup. Figure 9:**
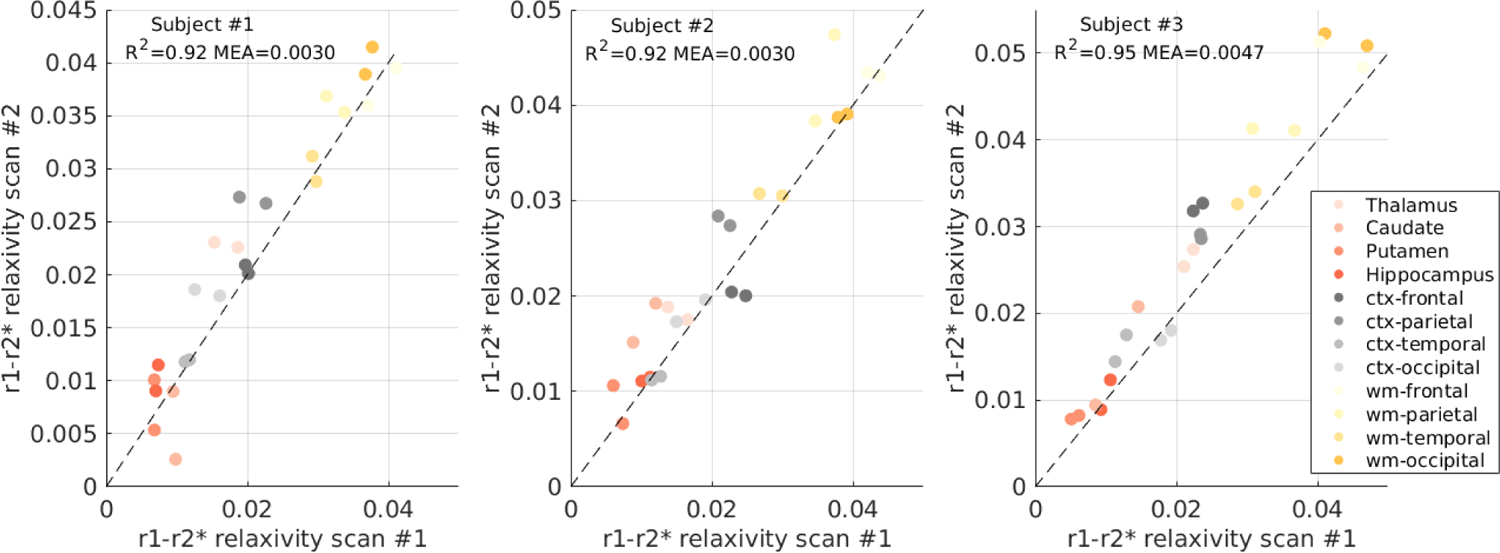
The reproducibility of the r1-r2* relaxivity measurement in the in vivo brain. The reproducibility of the r1-r2* relaxivity measurement in the in vivo brain was estimated based on scan-rescan experiments in three human subjects. Each subject was scanned twice in the MRI (on different days). The r1-r2* relaxivity was calculated for each scan in 12 different brain regions (different colors) in both hemispheres. Panels show the r1-r2* relaxivity values measured in the first scan (x-axis) vs. the r1-r2* relaxivity values measured in the scanned scan for each subject. Dashed line is the identity line. The measured scan-rescan mean absolute error (MAE) represents an experimental estimate of the detection limit of the r1-r2* relaxivity.

## Supplementary Section 3: Voxel-wise r1-r2* relaxivity visualization

Figure 2 and Figure 4 compare the contrast of R1 and R2* in the brain to the new contrast generated by the r1-r2* relaxivity. The measurement of the r1-r2* relaxivity is calculated across all the voxels of a specific ROI in the brain (see “r1-r2* relaxivity computation for ROIs in the human brain” in Methods). Therefore, the contrasts are presented across different entire brain regions. In order to demonstrate a visualization of a voxel-wise r1-r2* relaxivity contrast, we generated representative maps of the local r1-r2* relaxivity in a healthy young subject and in a Meningioma patient. For this purpose, we used a moving-window approach, in which the r1-r2* relaxivity of each voxel is based on the local linear dependency of R1 on R2* in that voxel and all its neighboring voxels (125 voxels total, for more details see “Generating voxel-wise r1-r2* relaxivity visualizations” in Methods).

A comparison of the voxel-wise r1-r2* relaxivity to the R1 and R2* maps in the healthy brain can be seen in Sup. Figure 10. Similarly to the ROI-based approach (Figure 2), this voxel-wise comparison also shows that the r1-r2* relaxivity generates a new contrast in the brain compared to R1 and R2*. Interestingly, this local relaxivity contrast highlights the differences between superficial and deep white matter. Such contrast was previously suggested to be driven by the microscopic iron distribution^45^.

In meningioma patients, we show that the ROI-based approach for the r1-r2* relaxivity allows to enhance the contrast between tumor tissue and non-pathological tissue without contrast agent injection (Figure 4). A comparison of the voxel-wise r1-r2* relaxivity to the R1 and R2* maps and to the Gd-enhanced contrast in a representative meningioma patient can be seen in Sup. Figure 11. In this example, the boundaries of the tumor can be separated from the surrounding non-pathological tissue based on the voxel-wise contrast of the r1-r2* relaxivity. Importantly, for this patient we were able to replicate our ROI-based results on the voxel-wise level (Sup. Figure 12). Across voxels, we find that the r1-r2* relaxivity allows to distinguish between tumor tissue and non-pathological tissue better than R1 and R2* (for example, effect size for the difference between tumor and gray matter is more than 10 times larger in the r1-r2* relaxivity compared to R1 and R2*).

Nonetheless, these representative visualizations merely provide preliminary evidence for the adaptiveness of the r1-r2* relaxivity approach for voxel-wise analyses. Notably, the presented implementation still has some limitations. First, the moving-window approach used for calculating the local r1-r2* relaxivity leads to inherent smoothing. As a result, this approach is sensitive to partial volume effects for voxels on the border between tissue types, which could be driving the observed contrast between superficial and deep white matter and between tumor tissue and non-pathological tissue. In addition, the local computation of the r1-r2* relaxivity use fewer voxels compared to the ROI-based approach. It also does not include the binning procedure prior to the fitting which we used in the ROI-based approach. Therefore, this computation is less stable and is more sensitive to the inherent SNR of R1 and R2*, which is affected by magnetic field inhomogeneities, imperfections of the shim, and the heterogeneous magnetic susceptibility of the head. As a result, some of the calculated values in the brain are negative (4% of the voxels in the healthy subject and 11% of the voxels in the meningioma patient). These limitations should be accounted for prior to any future implementation of this voxel-wise approach for purposes other than visualization.

**Sup. Figure 10:**
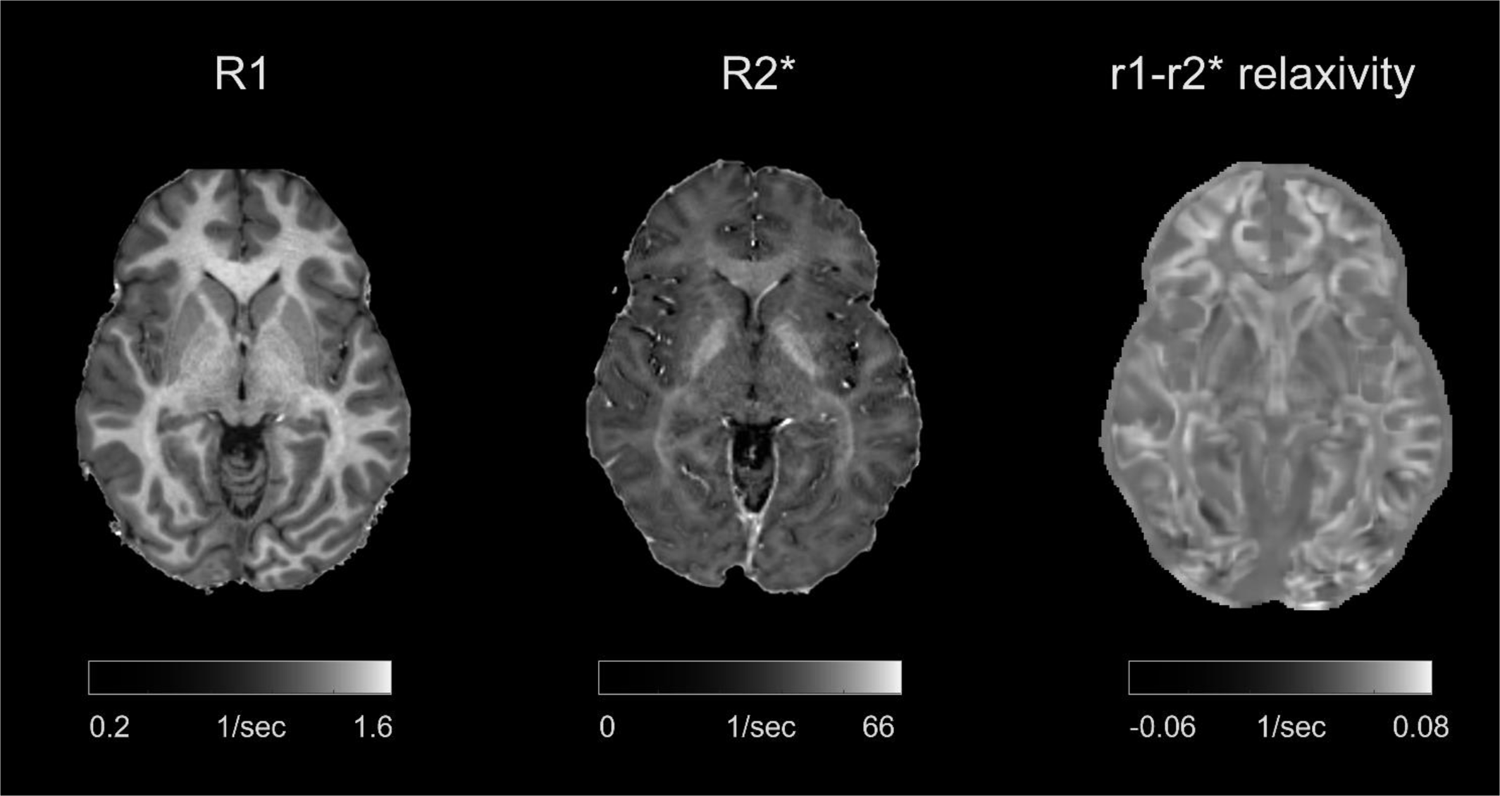
Voxel-wise comparison of the r1-r2* relaxivity map to R1 and R2* maps in the in vivo healthy brain. The maps of R1 (left) and R2* (middle) are compared to the local r1-r2* relaxivity visualization (right) on a representative young healthy subject. The voxel-wise visualization of the r1-r2* relaxivity in the brain was generated based on the local linear dependency of R1 on R2* using a moving-window approach (for more details see “Generating voxel-wise r1-r2* relaxivity visualizations” in Methods).

**Sup. Figure 11:**
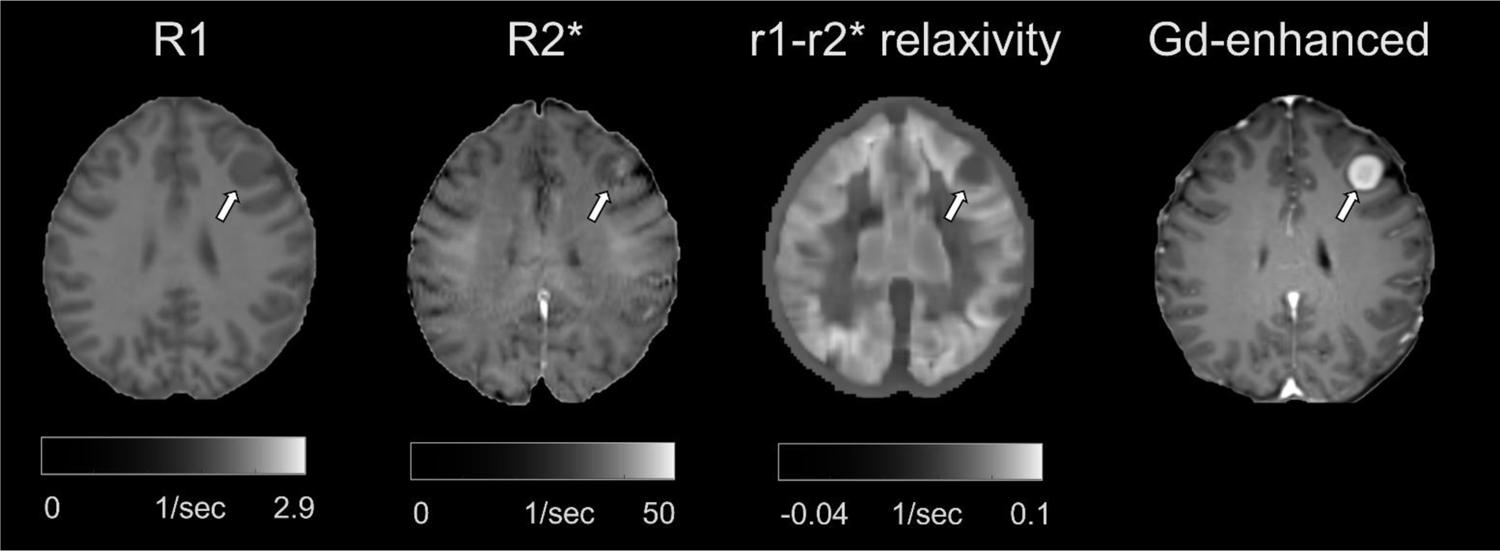
Voxel-wise comparison of the r1-r2* relaxivity map to R1 and R2* maps and to the Gd-enhanced contrast in the in vivo brain of a meningioma patient. Representative visualization of R1, R2*, the voxel-wise r1-r2* relaxivity map and the Gd-enhanced contrast for a meningioma patient. Tumors are marked with arrows. The voxel-wise r1-r2* relaxivity map was generated based on the local linear dependency of R1 on R2* using a moving-window approach (for more details see “Generating voxel-wise r1-r2* relaxivity visualizations” in Methods).

**Sup. Figure 12:**
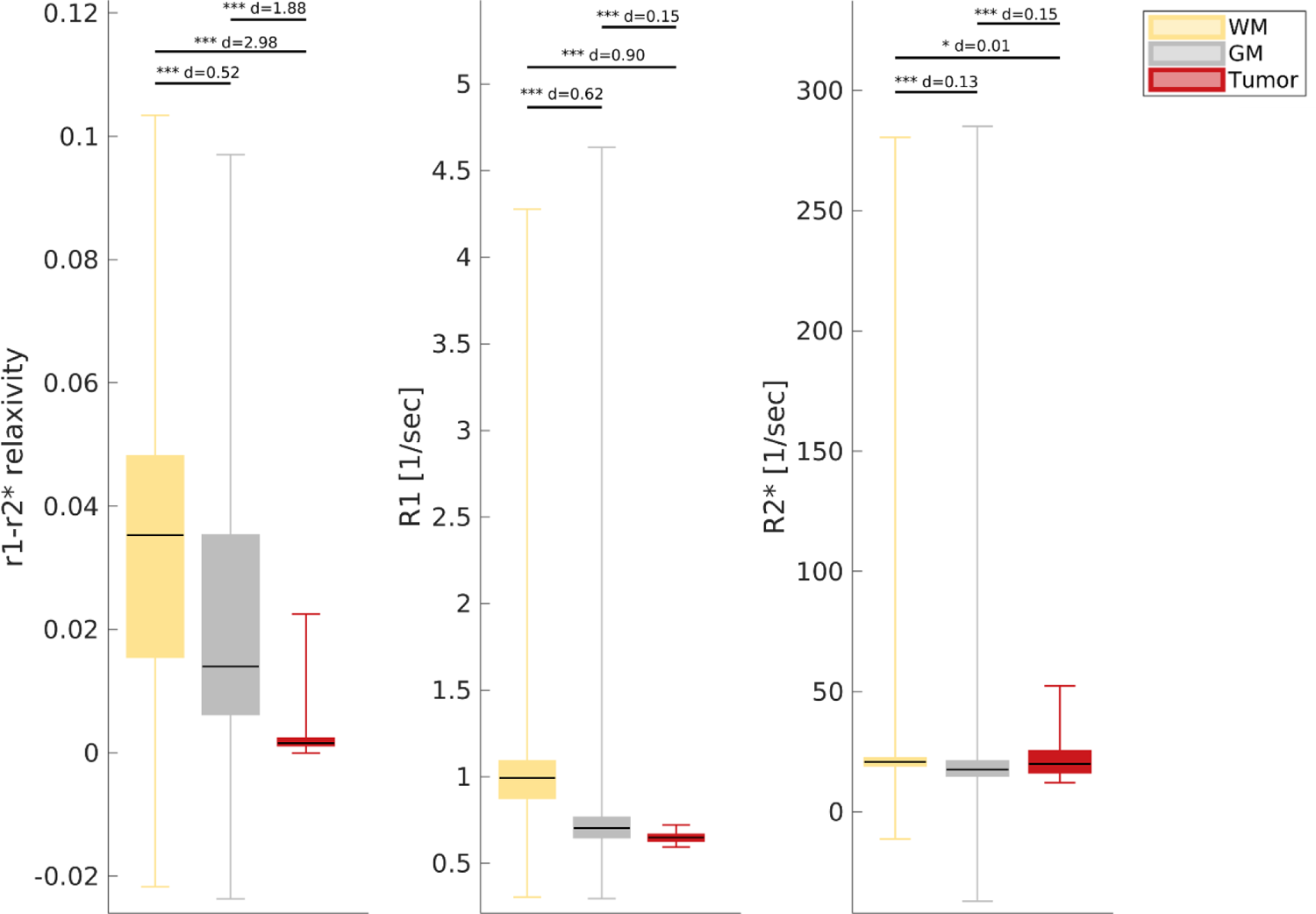
The voxel-wise r1-r2* relaxivity enhances the contrast between tumor tissue and non-pathological tissue on a representative meningioma patient. Replication of the ROI-based results presented in *figure 4d-f* on the voxel-wise level. The contrast between the white matter (WM), gray matter (GM) and tumor tissues is presented for R1, R2* and the voxel-wise r1-r2* relaxivity. The variation in each box is calculated across voxels in a representative meningioma patient (Sup. Figure 11). The 25th, 50th and 75th percentiles and extreme data points are shown. The d-values represent the effect size (Cohen’s d) of the differences between tissue types, and the significance level is based on a t-test. Across voxels, the r1-r2* relaxivity allows to distinguish between tumor tissue and non-pathological tissue better than R1 and R2*. Estimates in non-pathological tissues are for the tumor-free hemisphere. p < 0.05; **p < 0.01; ***p < 0.001

**Sup. Figure 13:**
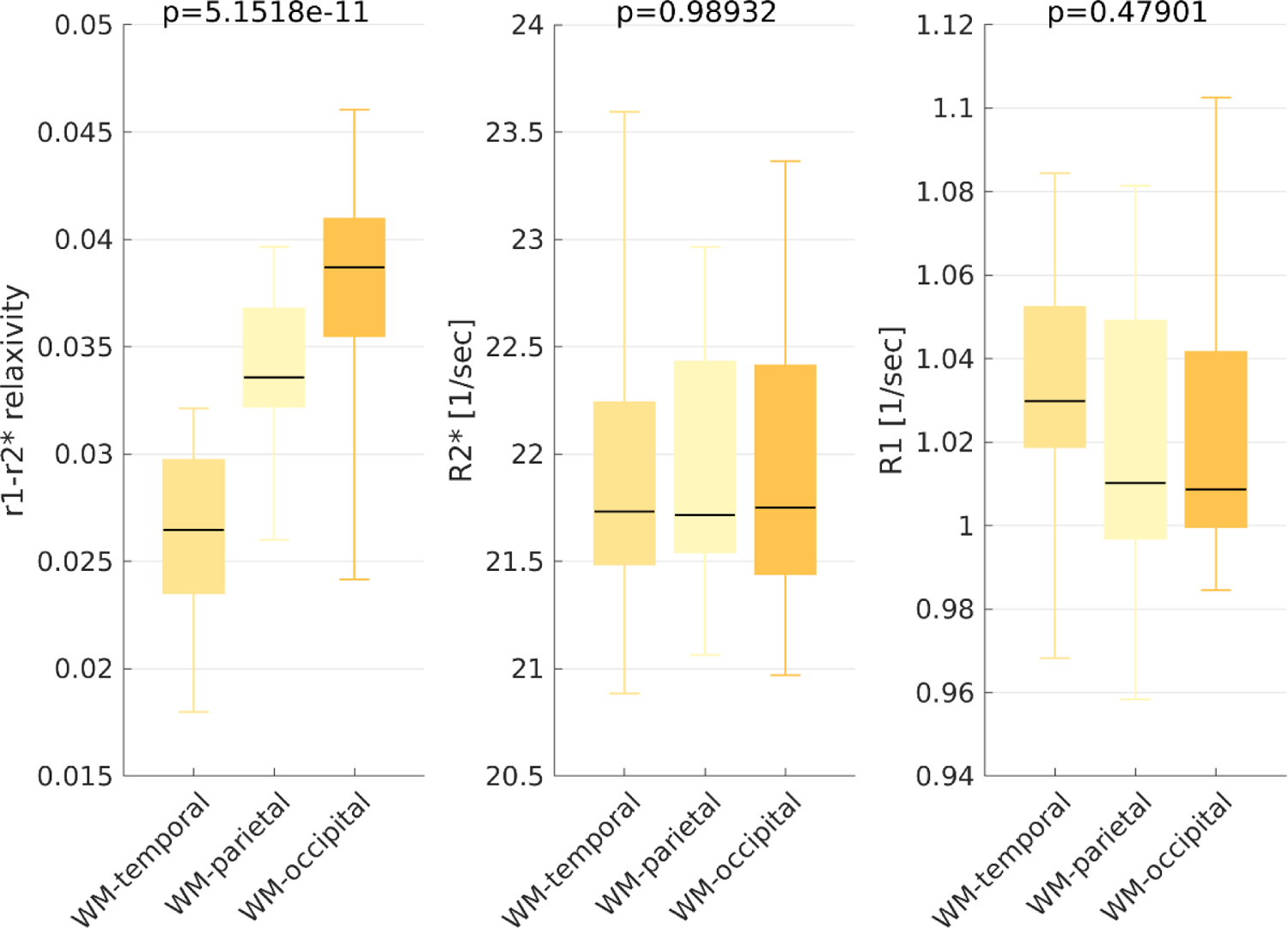
The new contrast of the r1-r2* relaxivity in white matter. From left to right, r1-r2* relaxivity, R2* and R1 for three different white-matter (WM) ROIs (temporal, parietal and occipital). We can separate these three WM ROIs with the r1-r2* relaxivity but not with either R2* or R1. Boxes represent the variation in the MRI parameters across normal subjects (age 27±2, N = 21). The 50^th^ percentile (horizontal black lines) 25th and 75th percentiles (box edges) and extreme data points (whiskers) are shown for each box. p-values are for the ANOVA test.

**Sup. Figure 14:**
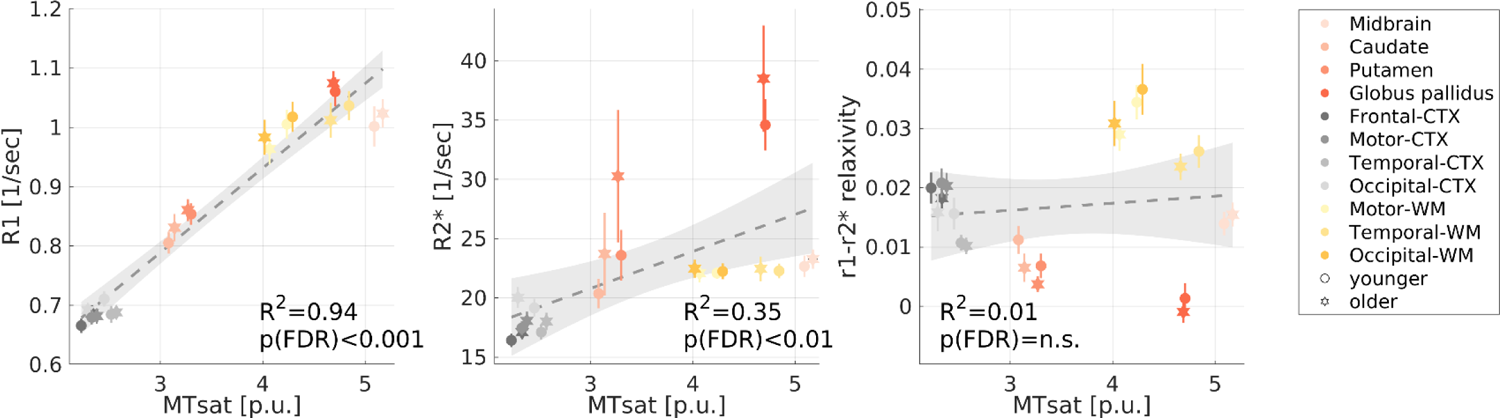
The correlations of MRI parameters with MTsat. The qMRI measurement of the magnetization transfer saturation (MTsat) vs. R1, R2* and the r1-r2* relaxivity measured in vivo across younger (aged 23-63 years, N =26) and older (aged 65-77 years, N=13) subjects (different marker shapes) in 10 brain regions (different colors). Unlike R1 and R2*, the r1-r2* relaxivity is not significantly correlated with MTsat.

**Sup. Figure 15:**
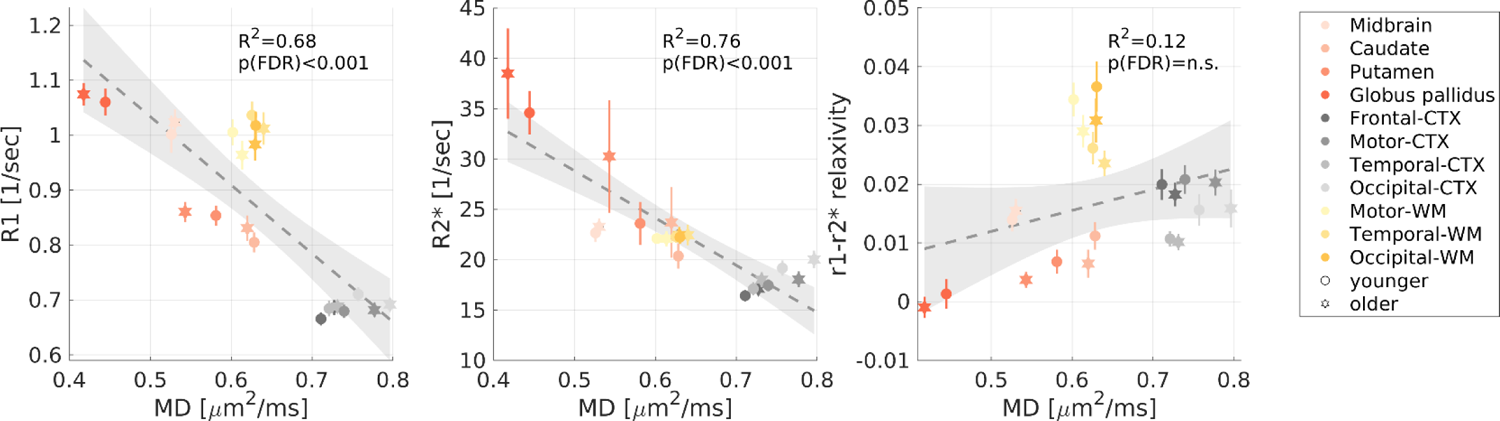
The correlations of MRI parameters with MD. The qMRI measurement of the mean diffusivity (MD) measured in vivo across younger (aged 23-63 years, N =25) and older (aged 65-77 years, N=12) subjects vs. R1, R2* and the r1-r2* relaxivity measured in vivo across younger (aged 23-63 years, N =26) and older (aged 65-77 years, N=13) subjects (different marker shapes) in 10 brain regions (different colors). Unlike R1 and R2*, the r1-r2* relaxivity is not significantly correlated with MD.

**Sup. Figure 16:**
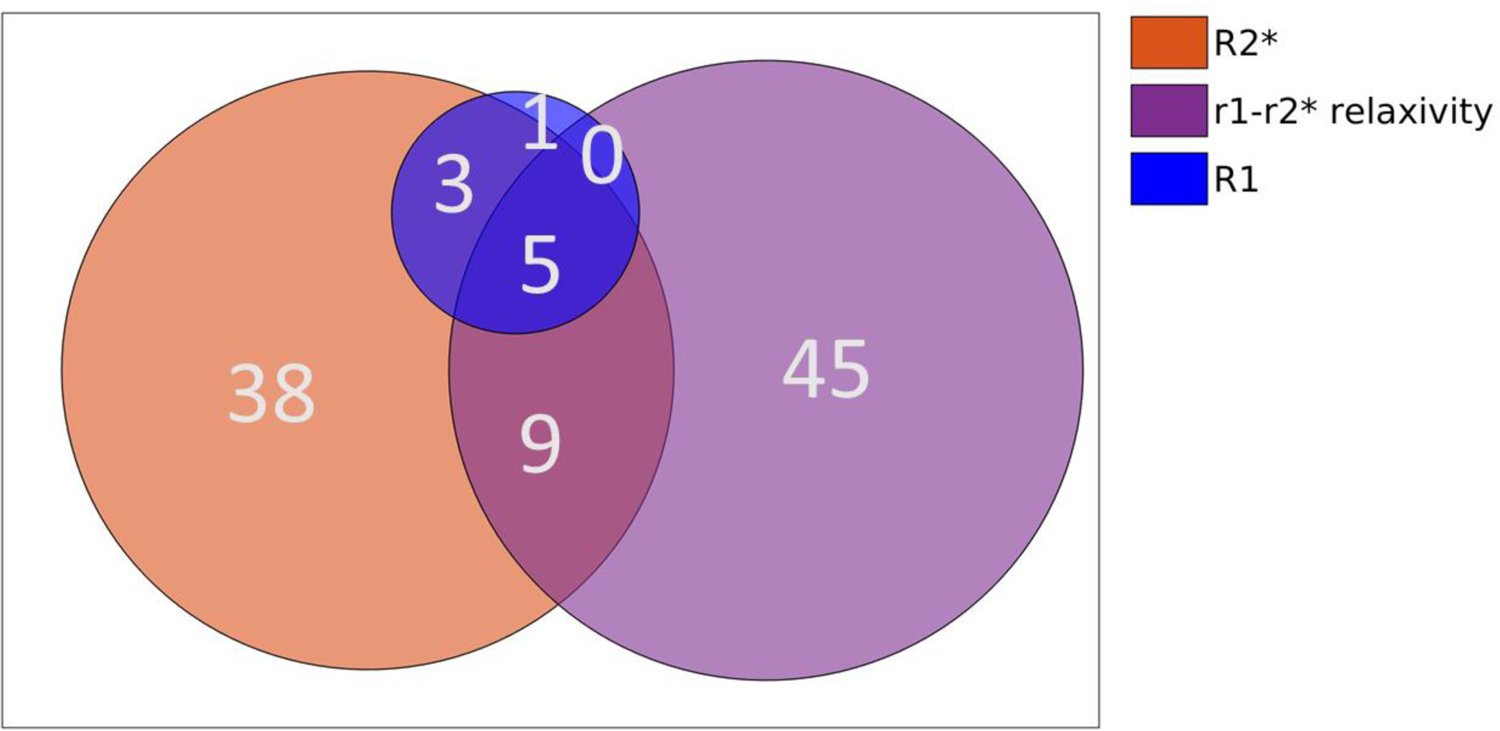
The number of significantly enriched pathways associated with each qMRI parameter. The Venn diagram shows the number of significantly enriched pathways (p(FWER)<0.01) for each qMRI parameter (R2*, R1 and the r1-r2* relaxivity). Almost half of the significantly enriched pathways are exclusive for the r1-r2* relaxivity. See also supplementary *table 1*.

**Sup. Table 1:**
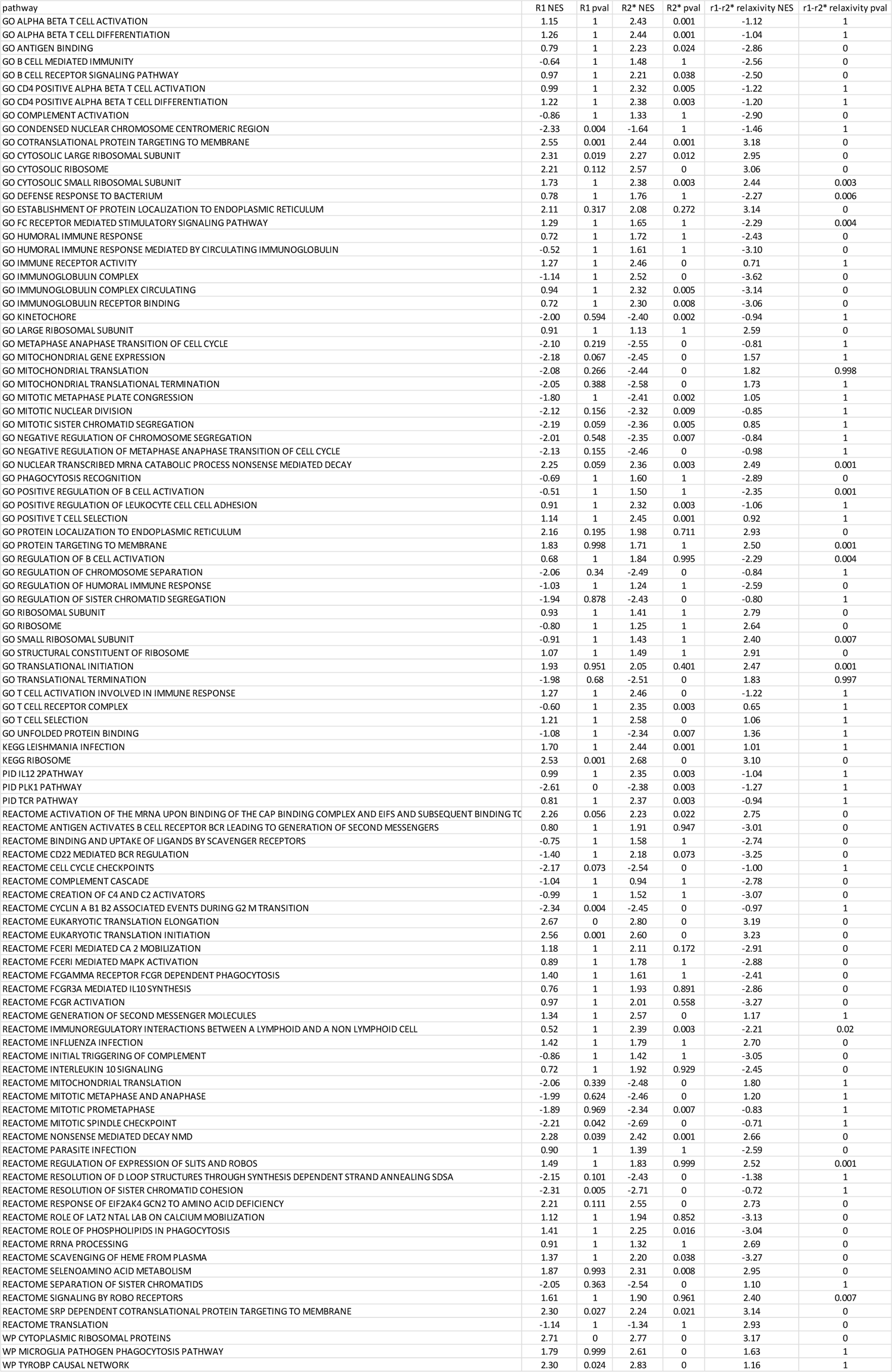
Significantly enriched pathways for R1, R2* and the r1-r2* relaxivity. For each of the 101 gene sets, we show the normalized enrichments score (NES) for each of the three qMRI parameters, along with the FWER-corrected p-value. This table was used for the clustering shown in *Figure 5b*.

**Sup. Figure 17:**
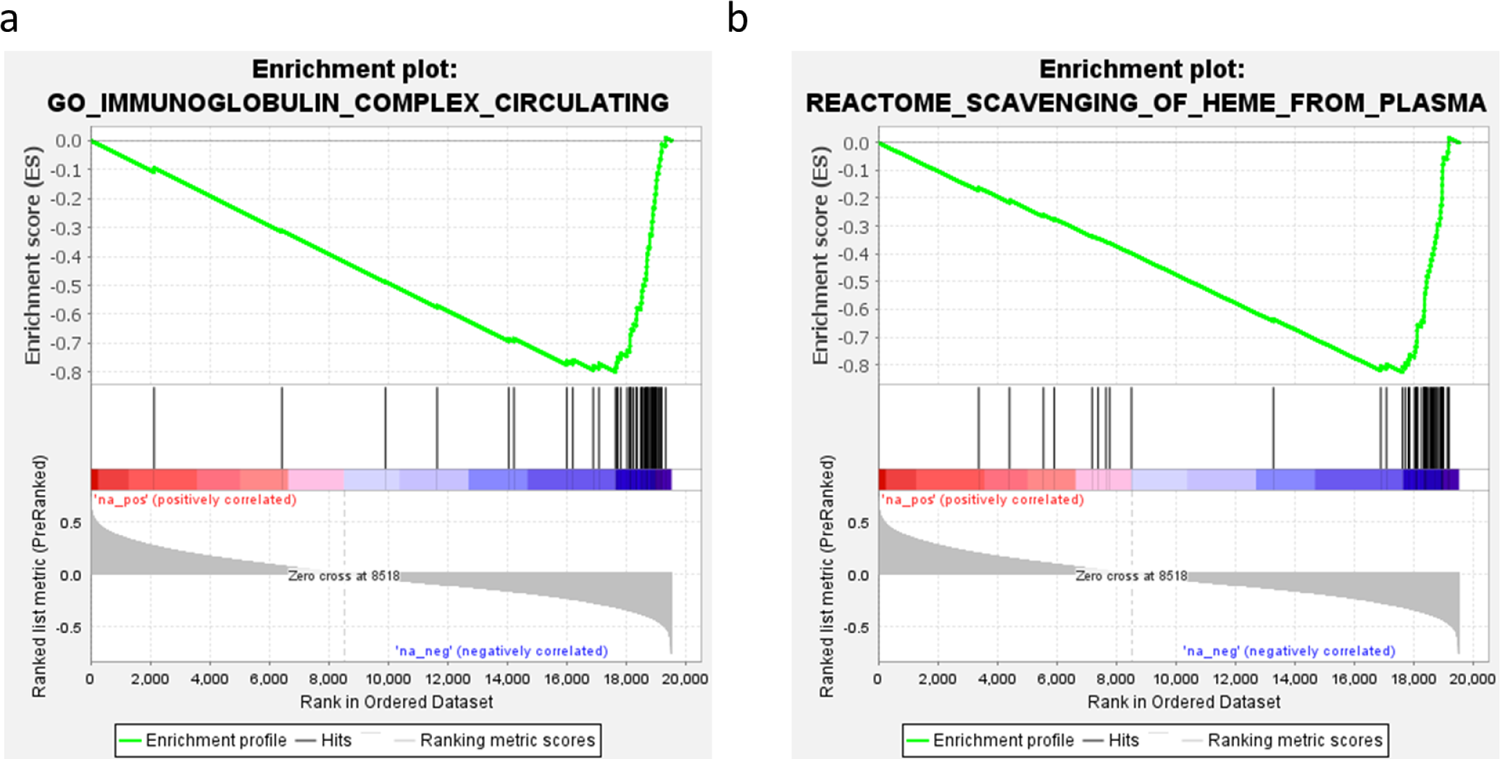
Enrichment plots of the two most enriched pathways for the r1-r2* relaxivity. Gene enrichment plots of the two most enriched pathways for the r1-r2* relaxivity “Immunoglobulin complex” **(a)** and “scavenging of heme from plasma” **(b)**. The top portion of each panel shows the running enrichment score for the gene set as the analysis goes over the ranked list of genes. The list is based on the genes’ correlation with the r1-r2* relaxivity. The middle portion of each panel shows where the members of the gene set appear in the ranked list of genes. The bottom portion of each panel shows the r value of the correlation between genes and the r1-r2* relaxivity. The two gene sets preferentially fall toward the negative end of the correlation spectrum, indicating their significant association with the r1-r2* relaxivity.

**Sup. Figure 18:**
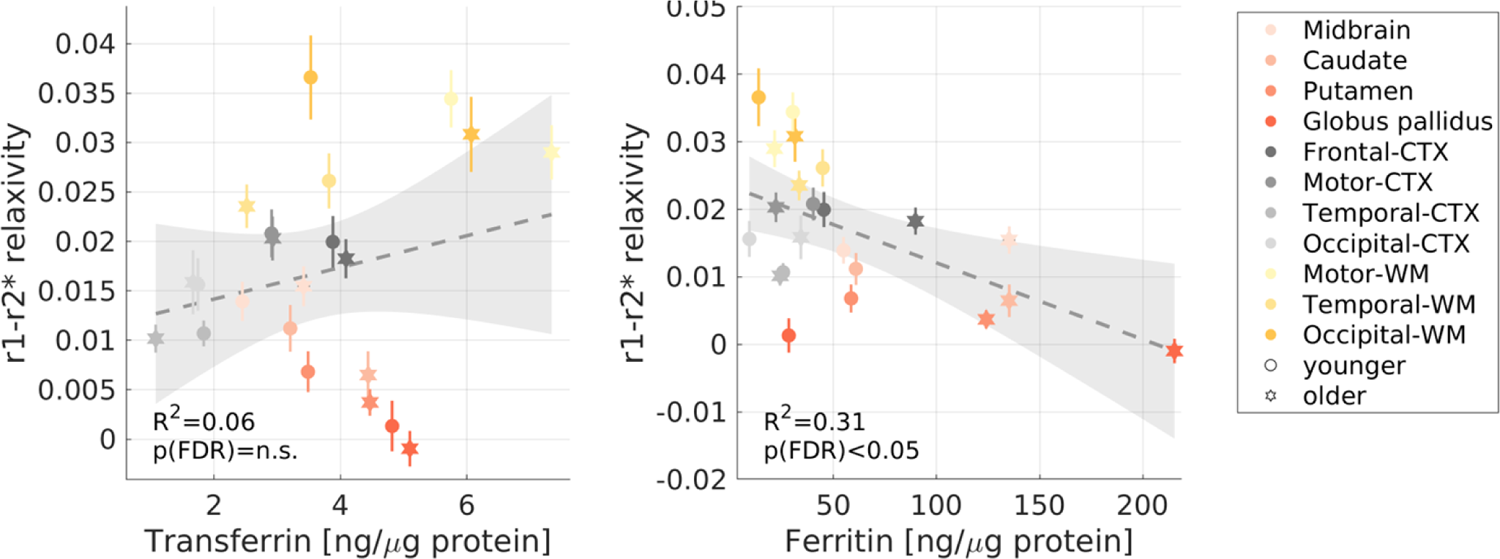
The correlations of the r1-r2* relaxivity with the transferrin and ferritin concentrations. The transferrin and ferritin concentrations (postmortem, from the literature^5,7,9^) in different brain regions of younger (aged 27-64 years, N>=7) and older (aged 65-88 years, N>=8) subjects vs. the r1-r2* relaxivity measured in vivo across younger (aged 23-63 years, N =26) and older (aged 65-77 years, N=13) subjects (different marker shapes) in 10 brain regions (different colors).

## Supplementary Section 4: Exploring the biophysical sources of the r1-r2* relaxivity

### Supplementary Section 4.1: The theoretical basis for the r1-r2* relaxivity of brain tissue

Brain tissue contains a complex milieu of iron compounds with variable iron binding capacities and aggregation states and includes myelin. Here we will expand the biophysical model of the r1-r2* relaxivity (“*In vivo* iron relaxivity model” in Methods) for the case of a heterogenous iron environment and in the presence of myelin.

Assuming the iron environment of a myelinated tissue contains N different iron compounds, and under the assumption that water can freely diffuse, the MR relaxation rates can be expressed as^87, 88^:

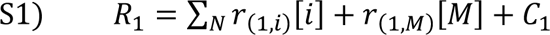

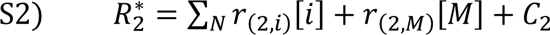

Where [i] is the concentration of the i’th iron compound, and [M] is the myelin concentration. r_(1,i)_ and r_(2,i)_ are the R1- and R2*-relaxivities of the i’th iron compound. r_(1,M)_ and r_(2,M)_ are the R1- and R2*-relaxivities of myelin. C_1_ and C_2_ are constants which capture other non-iron and non-myelin contributions.

The r1-r2* relaxivity measurement is defined as the linear dependency of R1 on R2* within an ROI in the brain (or across *in vitro* samples). This is equivalent to the total change in R1 relative to the total change in 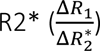

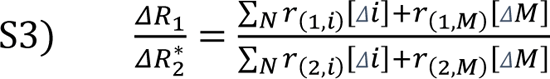

Where [Δi] and [ΔM] are the changes in the iron compounds and myelin concentrations within an ROI respectively.

### Supplementary Section 4.2: The r1-r2* relaxivity of ferritin and transferrin mixtures in vitro

First, we tested the theoretical formulation presented in eq. 1-S3 in an artificial environment of multiple iron compounds. For this aim, we constructed phantom experiments containing both ferritin and transferrin in a liposomal environment. In this synthetic toy model, the transferrin-ferritin fraction 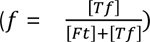 represents an example for a feature of the molecular iron environment which should affect the r1-r2* relaxivity. To test this assumption, the biophysical model shown in Supp. Section 4.1 can be further developed under the conditions of these phantom experiments:

When ferritin, transferrin and myelin are present, eq. 1-S2 can be expressed as:

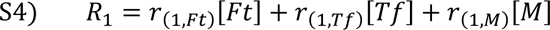

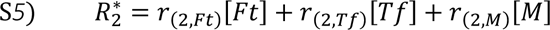

Where [Ft], [Tf] and [M] are the ferritin, transferrin and myelin concentrations respectively. r_(1,Ft)_, r_(1,Tf)_ and r_(1,M)_ are the R1-relaxivities of ferritin, transferrin and myelin respectively. r_(2,Ft)_, r_(2,Tf)_ and r_(2,M)_ are the R2*-relaxivities of ferritin, transferrin and myelin respectively.

According to eq. 3, the r1-r2* relaxivity measurement is equivalent to the total change in R1 relative to the total change in 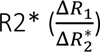

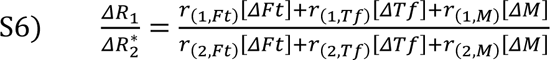

Where [ΔFt], [ΔTf], and [ΔM] are the changes in the ferritin, transferrin and myelin concentrations across *in vitro* samples, respectively.

In the ferritin-transferrin mixtures experiments we tested four different transferrin-ferritin fractions 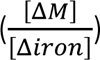. For each fraction, we prepared samples with varying ferritin and transferrin concentrations, while keeping the fixed fraction between them (**Sup. Figure 19**). This allowed us to fit the linear relationship between R1 and R2* (the r1-r2* relaxivity) for each transferrin-ferritin fraction (**Sup. Figure 20**).

Assuming the transferrin-ferritin fraction (f) remains fixed across the *in vitro* samples over which the r1-r2* relaxivity is calculated:

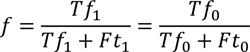

(defining [ΔTf] = Tf_1_ − Tf_0_ and [ΔFt] = Ft_1_ − Ft_0_).

Under this condition:

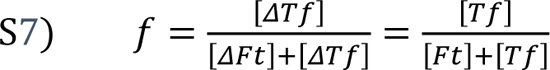

And therefore eq. 6 can be expressed as:

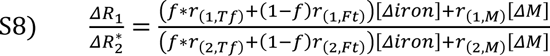

Where [ΔFt] + [ΔTf] = [Δiron].

In the ferritin-transferrin mixtures experiments, the liposomal fraction, which mimics the effect of myelin, was fixed at 17.5%. Therefore, in this case [ΔM] = 0 and eq. 8 reduces to:

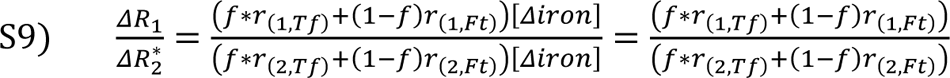

Using the ferritin and transferrin relaxivities (r_(1/2,Tf/Ft)_) measured for each liposomal iron compound individually (Figure 1a-c), we could test the prediction of this model. Notably, the theoretical r1-r2* relaxivities calculated with eq. 9 were in agreement with the experimental r1-r2* relaxivities (**Sup. Figure 21**), validating the presented biophysical framework *in vitro*. Moreover, the differences in the r1-r2* relaxivities measured for different transferrin-ferritin fractions were above the detection limit of this MRI measurement as estimated in a scan-rescan experiment (MAE=3.9*10^-4^, **Sup. Figure 8**). Therefore, this toy example demonstrates that the heterogeneity in the iron environment can produce measurable changes in the r1-r2* relaxivity, which are in agreement with the biophysical modeling of this measurement.

**Sup. Figure 19:**
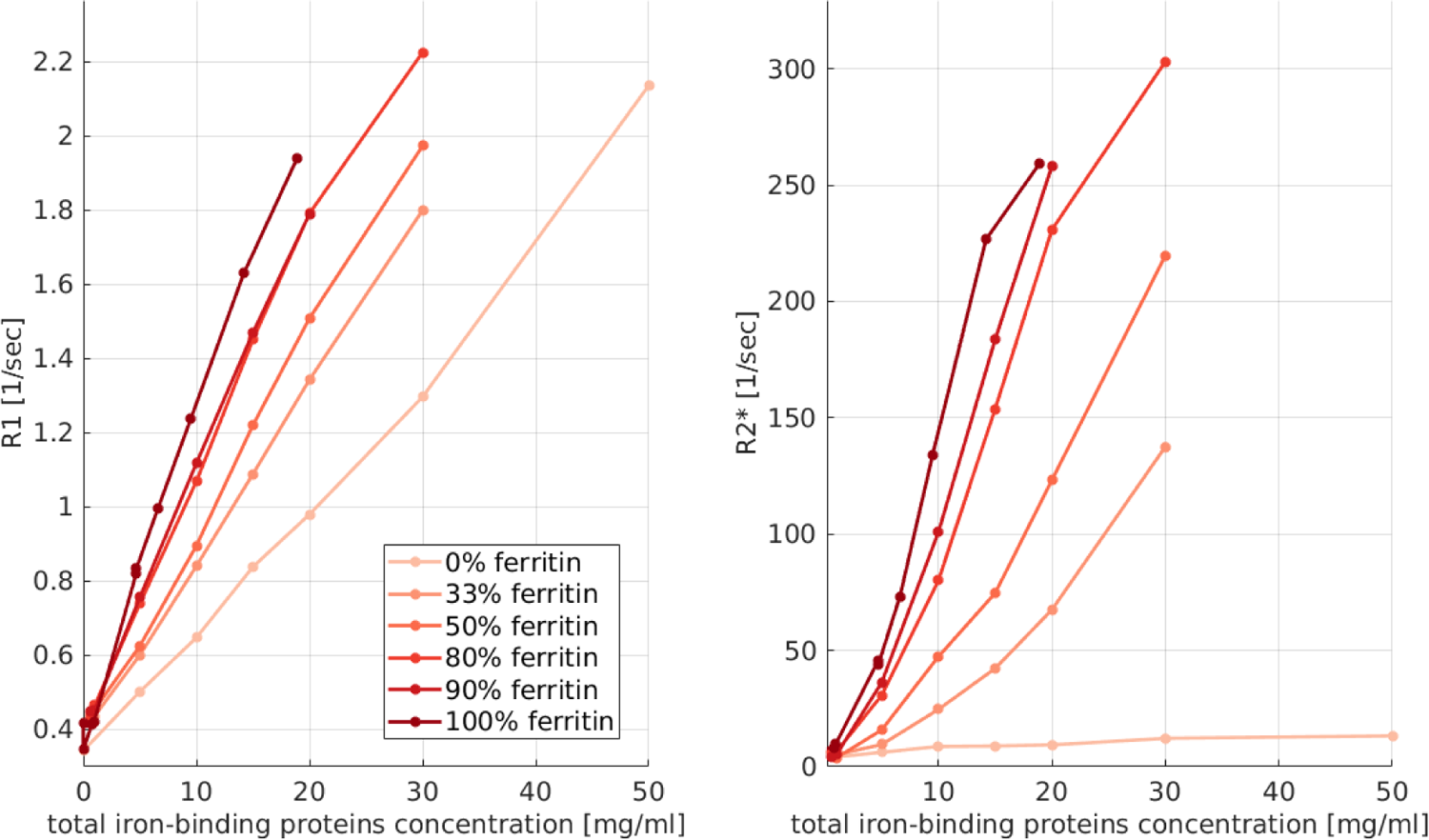
The dependency of R1 (left) and R2* (right) on the total iron-binding proteins concentration for four transferrin-ferritin mixtures. Each mixture has a different transferrin-ferritin fraction (different colors, legend shows the percentage of ferritin in the mixture). Data points are different transferrin-ferritin samples, line connect between samples with the same transferrin-ferritin fraction and varying total iron-binding protein concentrations.

**Sup. Figure 20:**
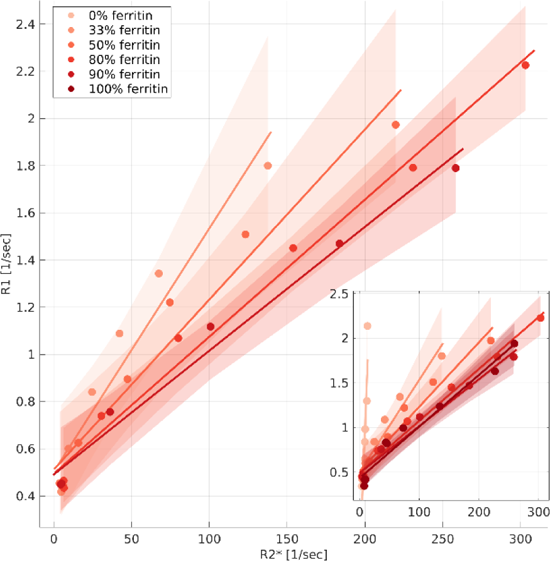
The dependency of R1 on R2* for four transferrin-ferritin mixtures. Each mixture has a different transferrin-ferritin fraction (different colors, legend shows the percentage of ferritin in the mixture). Data points represent samples with varying total iron-binding proteins concentrations. The linear relationships of R1 and R2* are marked by lines. The slopes of these lines are the r1-r2* relaxivities. Shaded areas represent the 95% confidence bounds. Inset shows the pure ferritin (100% ferritin) and pure transferrin (0% ferritin) samples as well.

**Sup. Figure 21:**
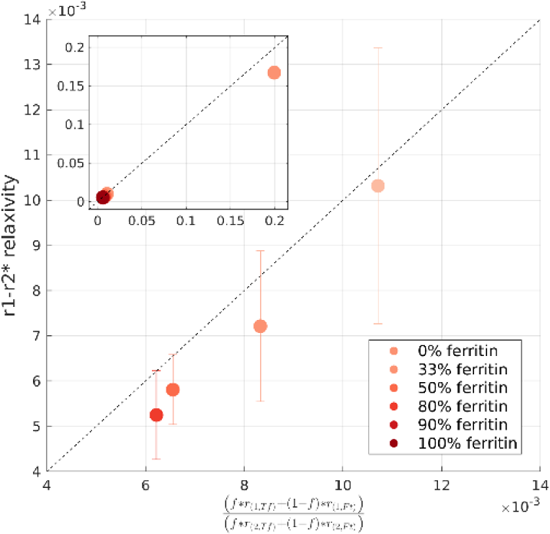
The theoretical r1-r2* relaxivities calculated with eq. 9 are in agreement with the experimental r1-r2* relaxivities. The y-axis shows the r1-r2* relaxivity calculated for four transferrin-ferritin mixtures with different transferrin-ferritin fractions (different colors, Sup. Figure 20). Errorbars show the 95% confidence bounds. The x-axis shows the prediction for the r1-r2* relaxivity based on eq. 9. The ferritin and transferrin relaxivities in the equation were plugged in based on our experimental results for liposomal ferritin and liposomal transferrin samples (Figure 1a-c). f represents the transferrin-ferritin fractions and varies between data points. Dashed line is the identity line. Inset shows the data points for pure ferritin (100% ferritin) and pure transferrin (0% ferritin) samples as well. Note that for these pure samples the prediction is already presented in *figure 1e*.

### Supplementary Section 4.3: Numerical simulations of the r1-r2* relaxivity

The phantom experiments of ferritin and transferrin mixtures allowed us to establish a theoretical framework for the r1-r2* relaxivity in an *in vitro* environment where only the iron concentration changes, and the liposomal fraction mimicking the myelin is fixed ([ΔM] = 0). In the brain, we estimate the r1-r2* relaxivity across all voxels of an anatomically-defined ROI. Within brain tissue ROIs, both the iron and the myelin concentration may vary^45^.

Importantly, rearranging eq. 8 we find that the strength of the myelin effect on the r1-r2* relaxivity depends on how variable is the myelin content within an ROI relative to how variable is the iron content 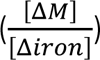:

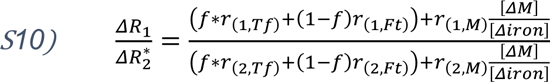

In order to evaluate how the r1-r2* relaxivity is modulated by the molecular iron environment and by the myelin and iron variabilities within an ROI in the brain, we performed a set of numerical simulations. In these simulations, similar to the *in vitro* mixtures experiments, the transferrin-ferritin fraction 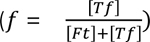 represents an example for a feature of the molecular iron environment which should affect the r1-r2* relaxivity.

In these analyses we aim to simulate realistic concentrations of ferritin, transferrin and myelin, in order to achieve brain-like R1 and R2* values. Next, we follow our analysis pipeline; binning of the R2* and R1 measurements, excluding bins with small number of voxels, and assessing the r1-r2* relaxivity across the binned values. By varying the simulated content of the myelin and iron compounds we test to what extent each biological source contributes to the measurement of the r1-r2* relaxivity. First, we will examine our hypothesis that changes in the molecular iron environment, reflected by the transferrin-ferritin fraction, but not in the iron concentration, affect the r1-r2* relaxivity. We will then evaluate how the r1-r2* relaxivity is modulated by myelin. We will show that non-physiological conditions are required in order for the myelin by itself to fully explain the r1-r2* relaxivity changes measured in the brain.

Each numerical simulation was designed to mimic an ROI in the brain containing 1M voxels, with a fixed transferrin-ferritin fraction (f) across all voxels and varying myelin, transferrin and ferritin concentrations (**Sup. Table 2**). We synthetically generated R1 and R2* values for each voxel based on eq.*S4*-S*5*. The relaxivities of ferritin and transferrin (r_(1/2,Ft)_, r_(1/2,Tf)_) were taken from the results of our phantom experiments (Figure 1). In order to generate simulations that are as realistic as possible, the rest of the parameters were adapted from the human brain. The mean ferritin and transferrin concentrations (across all voxels of the ROI) were estimated based on post-mortem findings^5,7,9^. The myelin characteristics were simulated based on the qMRI measurement of the macromolecular tissue volume (MTV)^46^, defined as 1-water fraction, which was shown to approximate the myelin content ^48–52^. The myelin relaxivity (r_(1/2,M)_) is defined as the dependency of relaxation rates on the myelin concentration^76^. We estimated the myelin relaxivity as the linear dependency of R1 and R2* on MTV, averaged across 16 ROIs in the brains of 21 young subjects. In order to assess the changes in myelin content within brain ROIs ([ΔM]), we calculated the range of MTV values within white-matter (WM) or gray-matter (GM) regions averaged across 8 ROIs in the brains of 21 young subjects (**Sup. Figure 22**). Finally, the changes in ferritin and transferrin concentrations within brain ROIs ([Δiron]) were determined based on the range of R2* values within 16 WM and GM regions in the brains of 21 young subjects. We assumed that the changes in R2* not explained by MTV are related to changes in iron concentration (**Sup. Figure 23**):

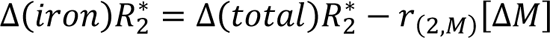

We found that the total change in R2* within ROIs in the human brain is on average Δ(total)R_2_^∗^=9.0 1/sec, from which about 61% (Δ(iron)R_2_^∗^=5.6 1/sec) could be related to changes in iron concentration. Therefore, the simulated variably in the ferritin and transferrin concentrations were set to satisfy this requirement.

An example result of the numerical simulation is presented in **Sup. Figure 24A**. It is evident that both the changes in myelin concentration across voxels of the simulated ROI (represented by different colors) and the changes in ferritin and transferrin concentrations across voxels (represented by the symbols size) affect the measured r1-r2* relaxivity. In our analysis pipeline we first bin the R1 and R2* values within the ROI and next calculate r1-r2* relaxivity over the binned values. Therefore, the variability in R1 for a given R2* bin is collapsed to an average R1 value (black data points). We assume that this approach eliminates some of the variability related to myelin. We hypothesized that the r1-r2* relaxivity is sensitive to the iron environment. Indeed, in our simulations we find that by setting different physiological transferrin-ferritin fractions and leaving the myelin parameters constant, the r1-r2* relaxivity changes considerably (**Sup. Figure 24**).

In addition, we hypothesized that the r1-r2* relaxivity less sensitive to the iron concentration and is more sensitive to the iron environment. To test this, we run two numerical simulations with the same transferrin-ferritin fraction but with different transferrin and ferritin concentrations. As expected, R1 and R2* values changed with increased ferritin and transferrin concentrations but the r1-r2* relaxivity did not change (**Sup. Figure 25**). This indicates that the r1-r2* relaxivity measurement is less sensitive to absolute changes in the iron concentration and is sensitive to the interplay between iron compounds, reflected in the simulations by the transferrin-ferritin fraction. In the brain, changes in the myelin content between GM and WM are known to substantially affect the measurements of R1 and R2*^4,19, 25, 29, 40–44^. To test the potential contribution of the myelin to the r1-r2* relaxivity, we changed the myelin concentration in our simulation while keeping the rest of the parameters fixed. Setting the myelin concentration to that typical for GM or WM (as estimated *in vivo* by MTV) led to considerable changes in R1 and R2*, but did not produce any change in the r1-r2* relaxivity (**Sup. Figure 26**).

The theoretical formulation presented here indicates that it is not the myelin concentration, but the variability in myelin within an ROI ([ΔM]), that is important for determining the r1-r2* relaxivity (eq. 10). However, estimating the variability in myelin within ROIs *in vivo* based on the myelin marker MTV, we find that it only explains ∼30% of the variation in the *in vivo* r1-r2* relaxivity measurements across the brain (**Sup. Figure 27**). In the simulations, setting both the range and concentration of myelin to the ones typical for WM or GM (as estimated by MTV, **Sup. Figure 22**), slightly changed the r1-r2* relaxivity (0.002, **Sup. Figure 28**). Importantly, changing the transferrin-ferritin fraction between the physiological values of 0.1-0.2 led to a change of 0.026 in the r1-r2* relaxivity (**Sup. Figure 24**). Therefore, the simulated changes related to the molecular iron environment were one order of magnitude bigger (**Sup. Figure 24**).

We further tested what are the myelin properties that would generate similar r1-r2* relaxivity effect as the effect observed when changing the transferrin-ferritin fraction (a change of 0.026 in the r1-r2* relaxivity, **Sup. Figure 24**). As changing the myelin concentration does not change the r1-r2* relaxivity (**Sup. Figure 26**), we changed the variability in myelin concentration within the ROI ([ΔM], **Sup. Figure 28**). We found that in order to generate a change of 0.026 in the r1-r2* relaxivity only through myelin-related changes (when the iron-related properties are fixed), the variability in MTV within the simulated ROI should be in the order of 0.187 [fraction] (**Sup. Figure 29**). Evaluating the *in vivo* variability in MTV within WM, GM and subcortical ROIs across 21 young subjects, the typical variability is ∼ 0.09 [fraction], and the most extreme variability that was measured was 0.13 [fraction] (in the WM, **Sup. Figure 22**). Even this atypical value is still much lower than that required to generate a change of 0.026 in the r1-r2* relaxivity (0.187 [fraction] in MTV). Thus, while the variability in myelin within ROIs can affect the r1-r2* relaxivity, there are no physiological myelin properties that would fully explain the r1-r2* relaxivity effect measured in the brain.

The iron and myelin contents of brain tissue are tightly related, as iron is required for the formation of myelin^4^. To test how this affects the r1-r2* relaxivity measurement, we simulated a case where iron and myelin are completely correlated. Importantly, even in this extreme case, different transferrin-ferritin fractions exhibited different r1-r2* relaxivity (**Sup. Figure 30**).

To conclude, we simulated a brain-like environment in order to test the biological sources affecting the r1-r2* relaxivity measurement. We found that physiological changes in the molecular iron environment, represented in the simulations by the ferritin-transferrin fraction, led to considerable changes in the r1-r2* relaxivity (0.026). This effect could not be attributed to the absolute concentrations of iron compounds, only to the ratio between them. To estimate whether the effect of changing the molecular iron environment is measurable *in vivo*, we assessed the detection limit of the r1-r2* relaxivity measurement using scan-rescan experiments (**Sup. Figure 9**). We found that the changes in the r1-r2* relaxivity simulated by different physiological transferrin-ferritin ratios are well above the detection limit of this MRI measurement *in vivo* (MEA∼0.0035). Importantly, our arguments regarding the sensitivity of the r1-r2* relaxivity to the molecular iron environment are demonstrated in the simulations based on the example of the transferrin-ferritin ratio. However, it is evident from our theoretical formulation (S10) that the variability in iron concentration within an ROI contributes to the r1-r2* relaxivity as well. Therefore, while the transferrin-ferritin ratio was used as an example, other features of the iron environment such as the spatial variability of iron compounds, their binding capacities and aggregate sizes, could affect the r1-r2* relaxivity as well. Next, we confirmed that changes in the myelin concentration affect the measurements of R1 and R2*, but not the r1-r2* relaxivity. The myelin variability within an ROI can affect the r1-r2* relaxivity. However, we found that *in vivo* estimates of this myelin characteristic explains only 30% of the variation in the *in vivo* r1-r2* relaxivity measurement across the brain. In the simulation, setting the myelin variability within an ROI to typical GM and WM values led to a slight change in the r1-r2* relaxivity. However, we found that unrealistic myelin variability is required in order to produce the r1-r2* relaxivity effect observed for realistic changes in the transferrin-ferritin fraction. Therefore, while the myelin substantially affects the measurements of R1 and R2*, it is not the main component governing the measurement of the r1-r2* relaxivity, and under physiological conditions it cannot by itself explain the measured variability in the r1-r2* relaxivity across the brain.

**Sup. Figure 22:**
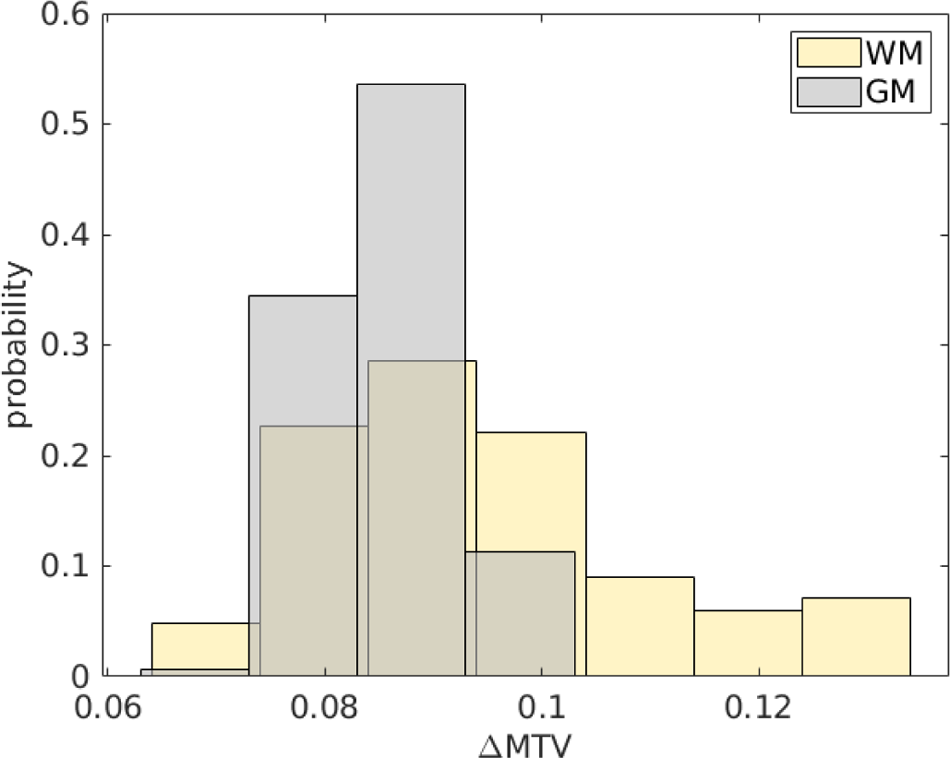
Change in MTV values ([ΔMTV]) within white-matter (WM) or gray-matter (GM) regions. ΔMTV values for gray matter (GM) and white matter (WM) are presented across 16 ROIs in the brains of 21 young subjects. For each ROI, we extracted the MTV values from all voxels and pooled them into 36 bins spaced equally between 0.05 and 0.40 [fraction]. We removed any bins in which the number of voxels was smaller than 4% of the total voxel count in the ROI. This was done so that the calculation will not be heavily affected by outlier voxels with extreme values. The median MTV of each bin was computed, and the difference between the highest and lowest binned MTV values was set as ΔMTV ([fraction]) in the ROI.

**Sup. Figure 23:**
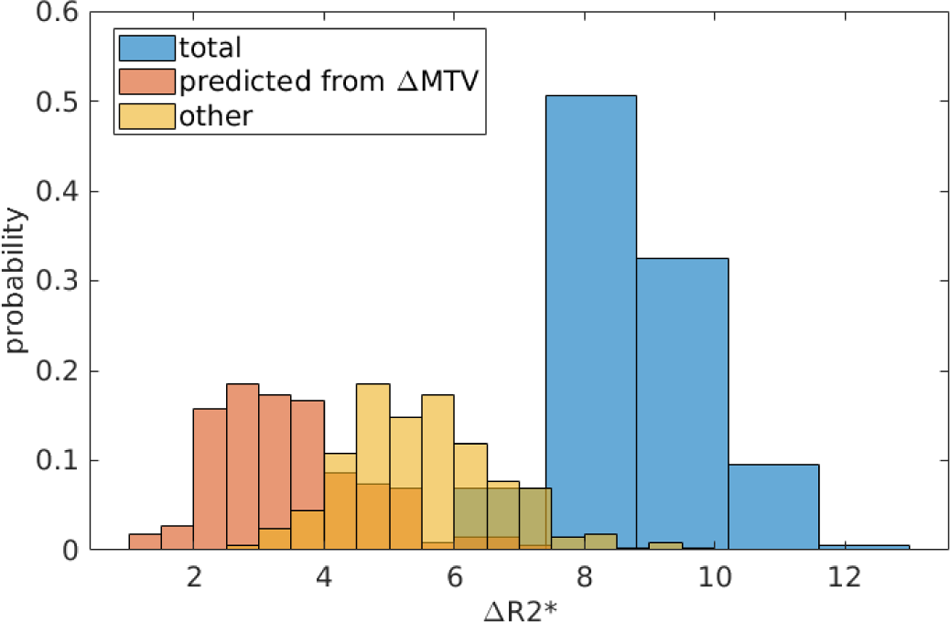
Myelin- and iron-related changes in R2* within brain regions. Total change in R2* values ([ΔR2 ∗]) within brain regions (blue histogram) is presented across 16 ROIs in the brains of 21 young subjects. For each ROI, we extracted the R2* values from all voxels and pooled them into 36 bins spaced equally between 0 and 50. We removed any bins in which the number of voxels was smaller than 4% of the total voxel count in the ROI. This was done so that the calculation will not be heavily affected by outlier voxels with extreme values. The median R2* of each bin was computed, and the difference between the highest and lowest binned R2* values was set as ΔR2* in the ROI (in [1/sec]). R2* changes related to myelin (orange histogram) were estimated based on MTV; the change in R2* predicted from the change in MTV (ΔMTV) within each ROI was calculated as the linear dependency of R2* on MTV in the ROI multiplied by ΔMTV in the ROI (r_(2,M)_[ΔM]). R2* changes related to iron (yellow histogram) were estimated as the change in R2* not explained by the change in MTV.

**Sup. Table 2:**
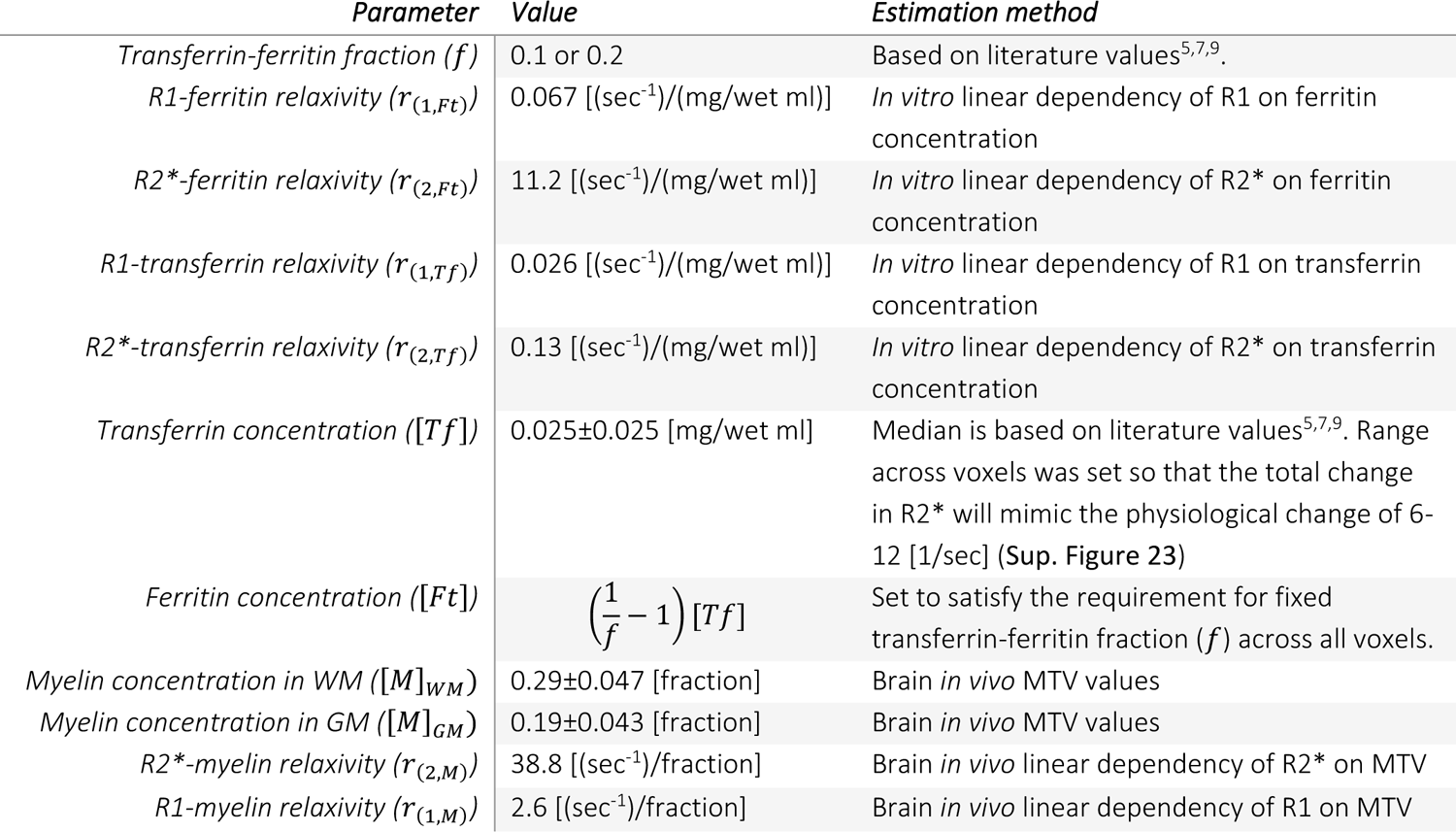
simulation parameters.

**Sup. Figure 24:**
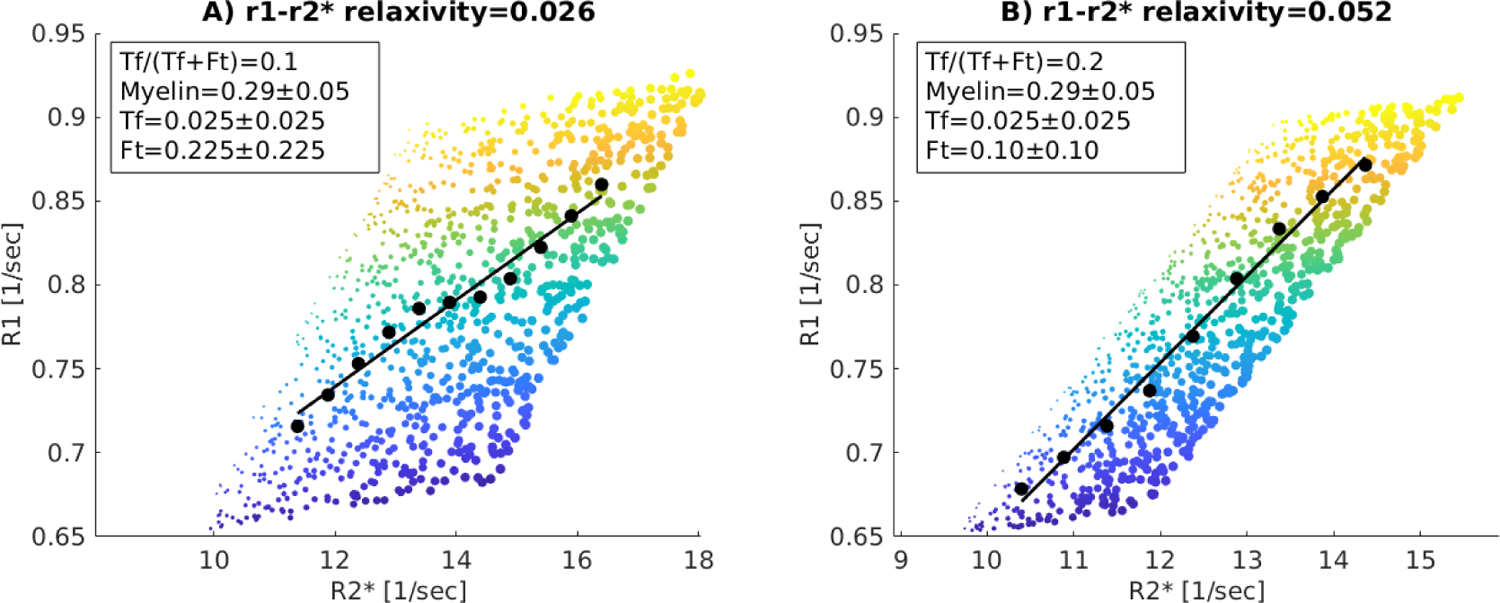
The r1-r2* relaxivity in two simulated ROIs with different transferrin-ferritin fractions (Tf/(Tf+Ft)); (A) the transferrin-ferritin fraction is 0.1; (B) the transferrin-ferritin fraction is 0.2. Each figure shows the dependency of R1 on R2* for 1,000 representative simulated voxels. The colors of the data points indicate the variability in myelin concentration across voxels, and their sizes indicate the variability in iron compounds concentration across voxels (the simulated concentrations are shown in the text box, myelin is in units of [fraction] as MTV, transferrin and ferritin are in units of [mg/ml]). As in our in vivo pipeline, R2* and R1 values were binned (black data points represent the bins’ median), and a linear fit was calculated (black line). The slopes of the linear fit (shown in the title) represent the dependency of R1 on R2* (r1-r2* relaxivity) and vary with the transferrin-ferritin fraction.

**Sup. Figure 25:**
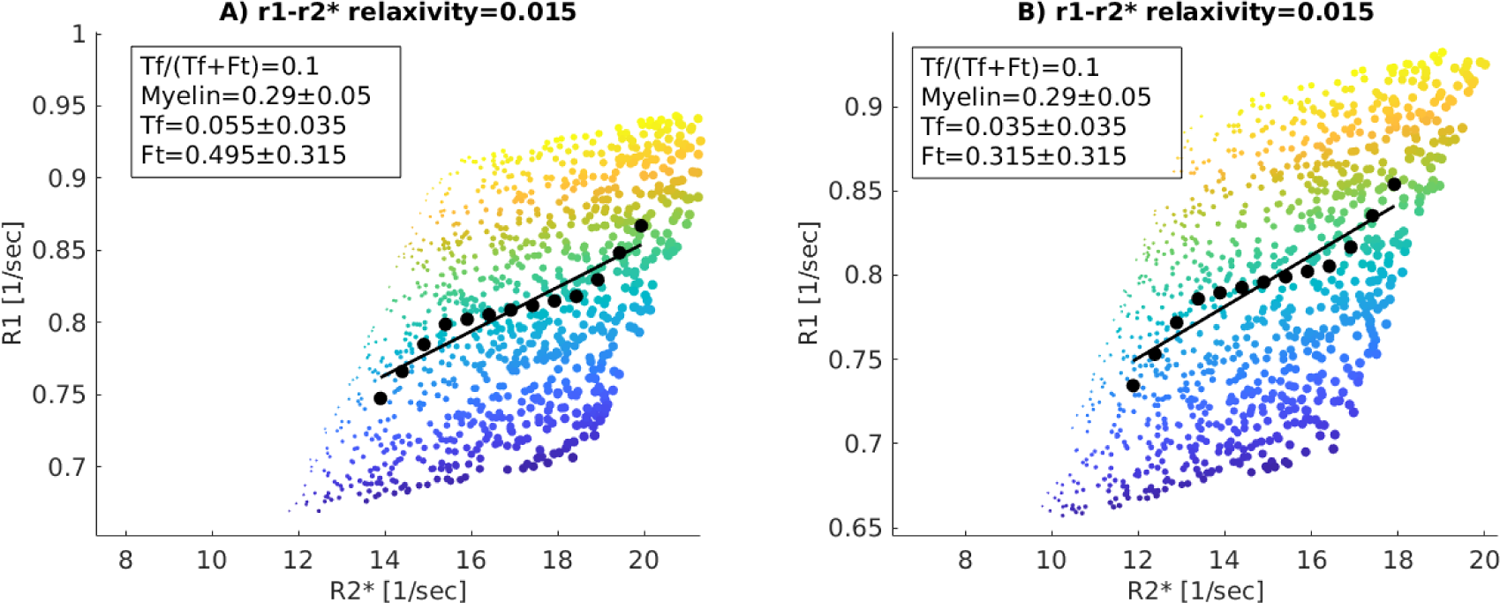
The r1-r2* relaxivity in two simulated ROIs with different transferrin and ferritin concentrations and a similar transferrin-ferritin fraction (Tf/(Tf+Ft)); (A) a higher transferrin and ferritin concentrations and a transferrin-ferritin fraction of 0.1; (B) a lower transferrin and ferritin concentrations and a transferrin-ferritin fraction of 0.1. Each figure shows the dependency of R1 on R2* for 1,000 representative simulated voxels. The colors of the data points indicate the variability in myelin concentration across voxels, and their sizes indicate the variability in iron compounds concentration across voxels (the simulated concentrations are shown in the text box, myelin is in units of [fraction] as MTV, transferrin and ferritin are in units of [mg/ml]). As in our in vivo pipeline, R2* and R1 values were binned (black data points represent the bins’ median), and a linear fit was calculated (black line). The slopes of the linear fit (shown in the title) represent the dependency of R1 on R2* (r1-r2* relaxivity) and do vary with the change in transferrin and ferritin concentrations.

**Sup. Figure 26:**
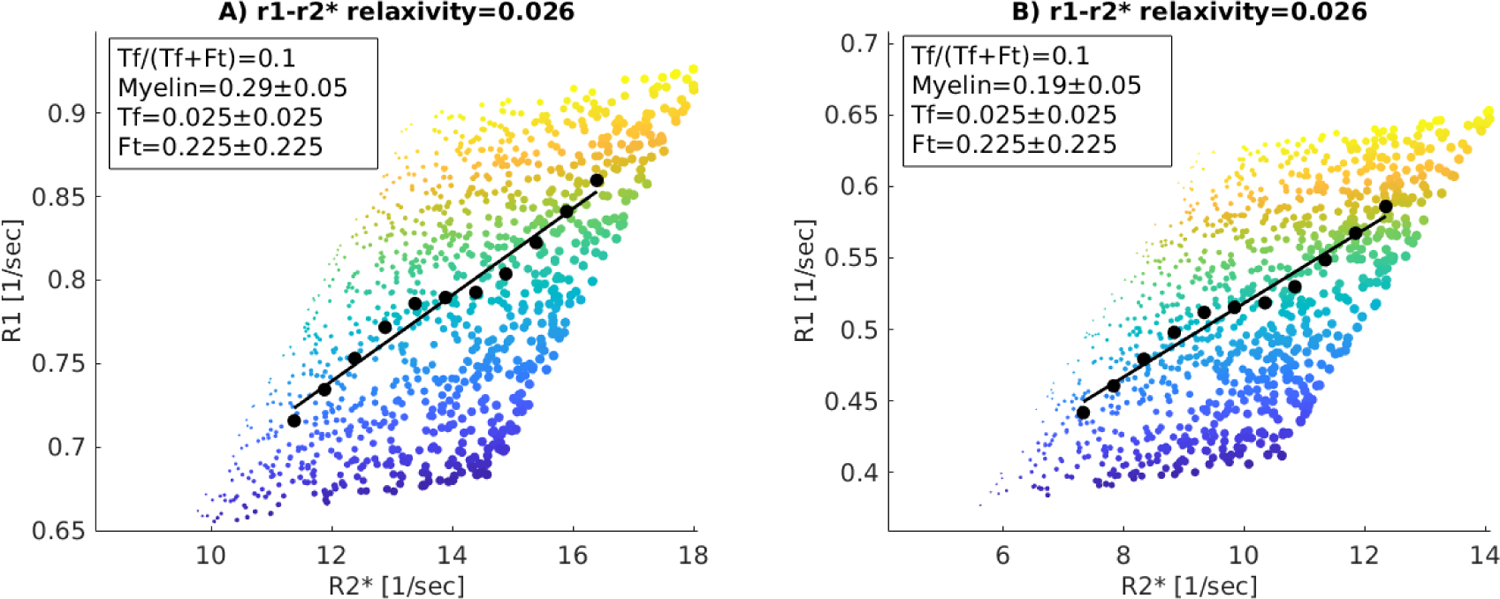
The r1-r2* relaxivity in two simulated ROIs with different myelin concentrations and similar transferrin-ferritin fractions (Tf/(Tf+Ft)); (A) a higher myelin concentration and transferrin-ferritin fraction of 0.1; (B) a lower myelin concentration and transferrin-ferritin fraction of 0.1. Each figure shows the dependency of R1 on R2* for 1,000 representative simulated voxels. The colors of the data points indicate the variability in myelin concentration across voxels, and their sizes indicate the variability in iron compounds concentration across voxels (the simulated concentrations are shown in the text box, myelin is in units of [fraction] as MTV, transferrin and ferritin are in units of [mg/ml]). As in our in vivo pipeline, R2* and R1 values were binned (black data points represent the bins’ median), and a linear fit was calculated (black line). The slopes of the linear fit (shown in the title) represent the dependency of R1 on R2* (r1-r2* relaxivity) and does not vary with the change in myelin concentration.

**Sup. Figure 27:**
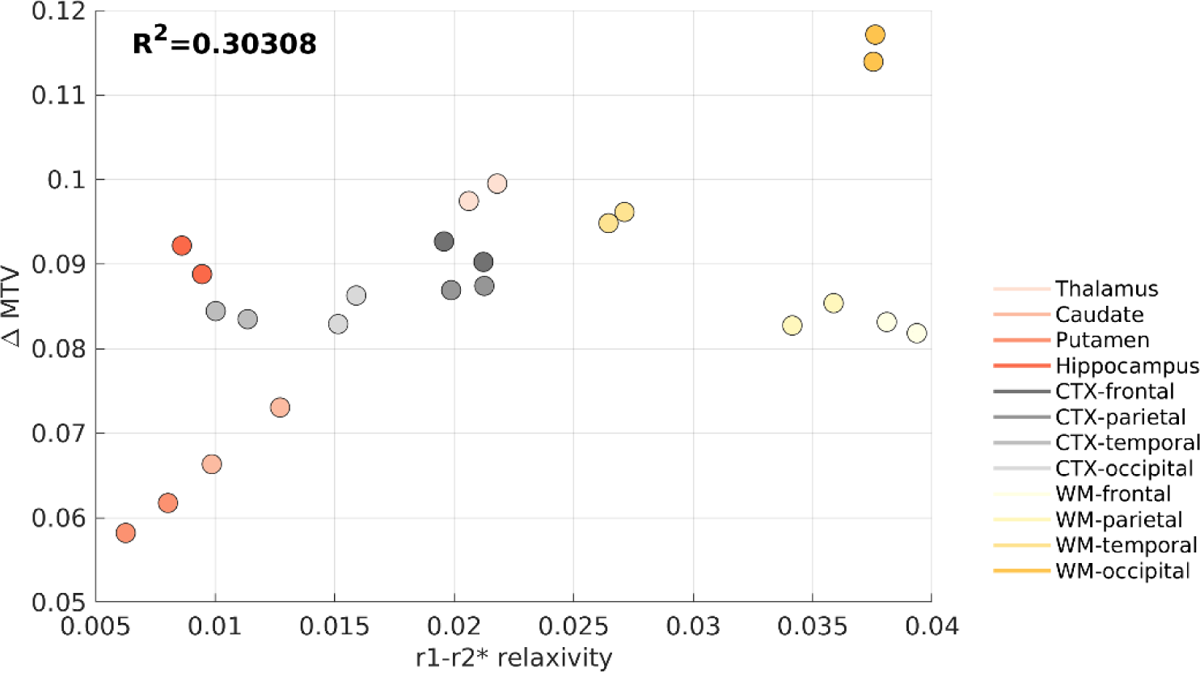
The in vivo estimates of the variability in myelin within ROIs vs. the in vivo r1-r2* relaxivity. The average estimated myelin variability within ROIs, computed across the brains of 21 young subjects, vs. the average r1-r2* relaxivity for the same subjects. Brain regions are presented in different colors (each ROI has left and right hemisphere estimates). The variability in myelin within brain regions was estimated based on the variability in MTV across voxels of each ROI (ΔMTV, y-axis). For each ROI, we extracted the MTV values from all voxels and pooled them into 36 bins spaced equally between 0.05 and 0.40 [fraction]. We removed any bins in which the number of voxels was smaller than 4% of the total voxel count in the ROI. This was done so that the calculation will not be heavily affected by outlier voxels with extreme values. The median MTV of each bin was computed, and the difference between the highest and lowest binned MTV values was set as ΔMTV ([fraction]) in the ROI.

**Sup. Figure 28:**
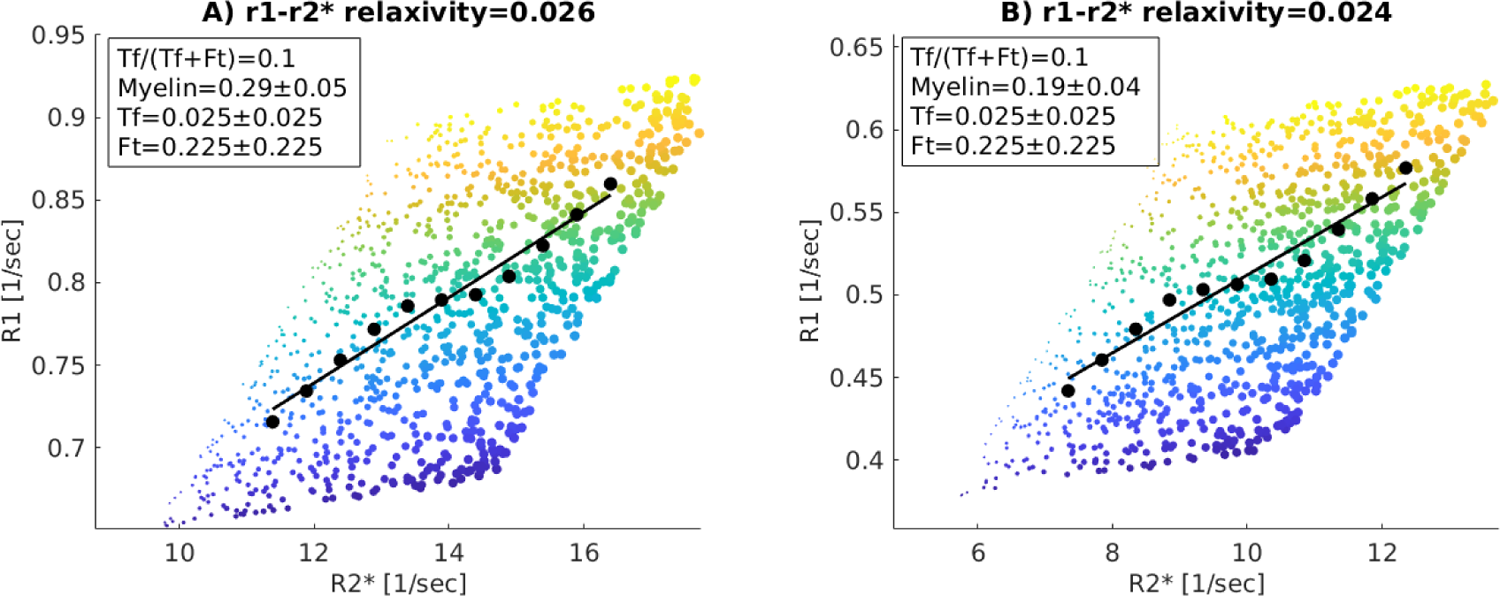
The r1-r2* relaxivity in two simulated ROIs with different myelin concentrations, different ranges of myelin concentrations ([ΔM]), and similar transferrin-ferritin fractions (Tf/(Tf+Ft)); (A) WM; a higher myelin concentration and a larger range of myelin variability, the transferrin-ferritin fraction is 0.1; (B) GM; a lower myelin concentration and a lower range of myelin variability, the transferrin-ferritin fraction is 0.1. Each figure shows the dependency of R1 on R2* for 1,000 representative simulated voxels. The colors of the data points indicate the variability in myelin concentration across voxels, and their sizes indicate the variability in iron compounds concentration across voxels (the simulated concentrations are shown in the text box, myelin is in units of [fraction] as MTV, transferrin and ferritin are in units of [mg/ml]). As in our in vivo pipeline, R2* and R1 values were binned (black data points represent the bins’ median), and a linear fit was calculated (black line). The slopes of the linear fit (shown in the title) represent the dependency of R1 on R2* (r1-r2* relaxivity) and vary slightly with the range of myelin variability ([ΔM]).

**Sup. Figure 29:**
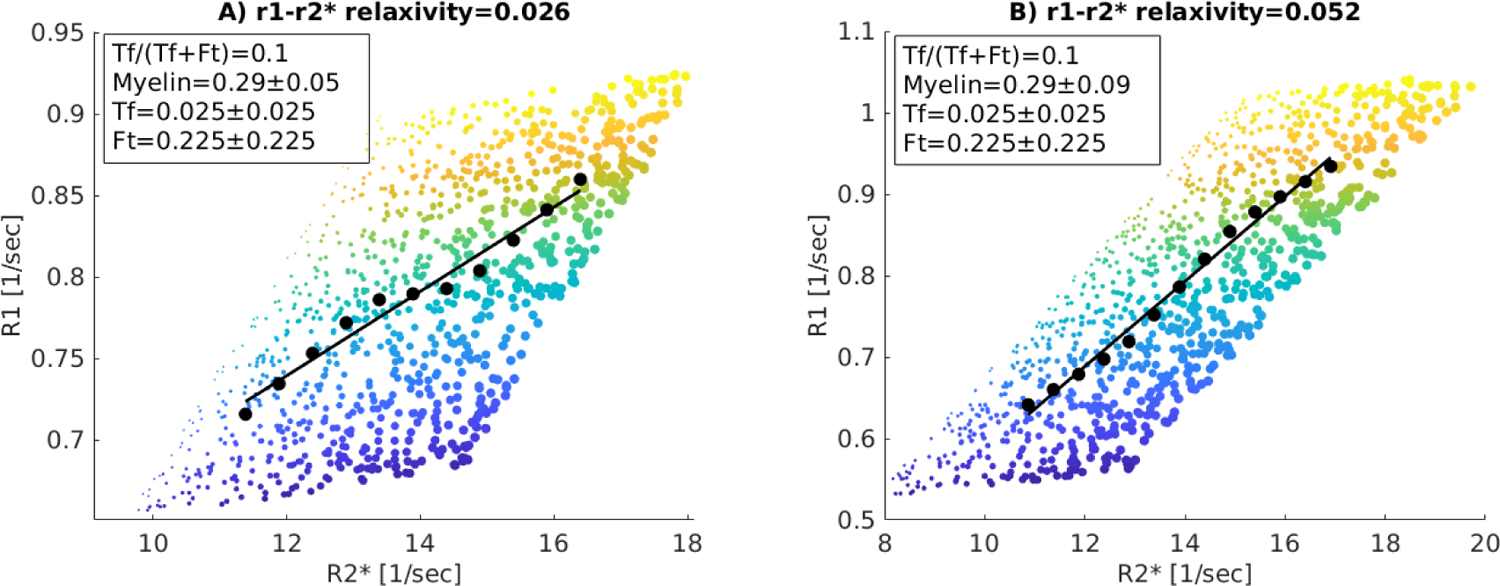
The r1-r2* relaxivity in two simulated ROIs with different extreme ranges of myelin concentrations ([ΔM]) and similar transferrin-ferritin fractions (Tf/(Tf+Ft)); (A) a physiological range of myelin variability, and a transferrin-ferritin fraction of 0.1; (B) the range of myelin variability is almost doubled, and the transferrin-ferritin fraction is 0.1. Each figure shows the dependency of R1 on R2* for 1,000 representative simulated voxels. The colors of the data points indicate the variability in myelin concentration across voxels, and their sizes indicate the variability in iron compounds concentration across voxels (the simulated concentrations are shown in the text box, myelin is in units of [fraction] as MTV, transferrin and ferritin are in units of [mg/ml]). As in our in vivo pipeline, R2* and R1 values were binned (black data points represent the bins’ median), and a linear fit was calculated (black line). The slopes of the linear fit (shown in the title) represent the dependency of R1 on R2* (r1-r2* relaxivity) and change considerably under this condition of extreme variability in myelin concentration ([ΔM]).

**Sup. Figure 30:**
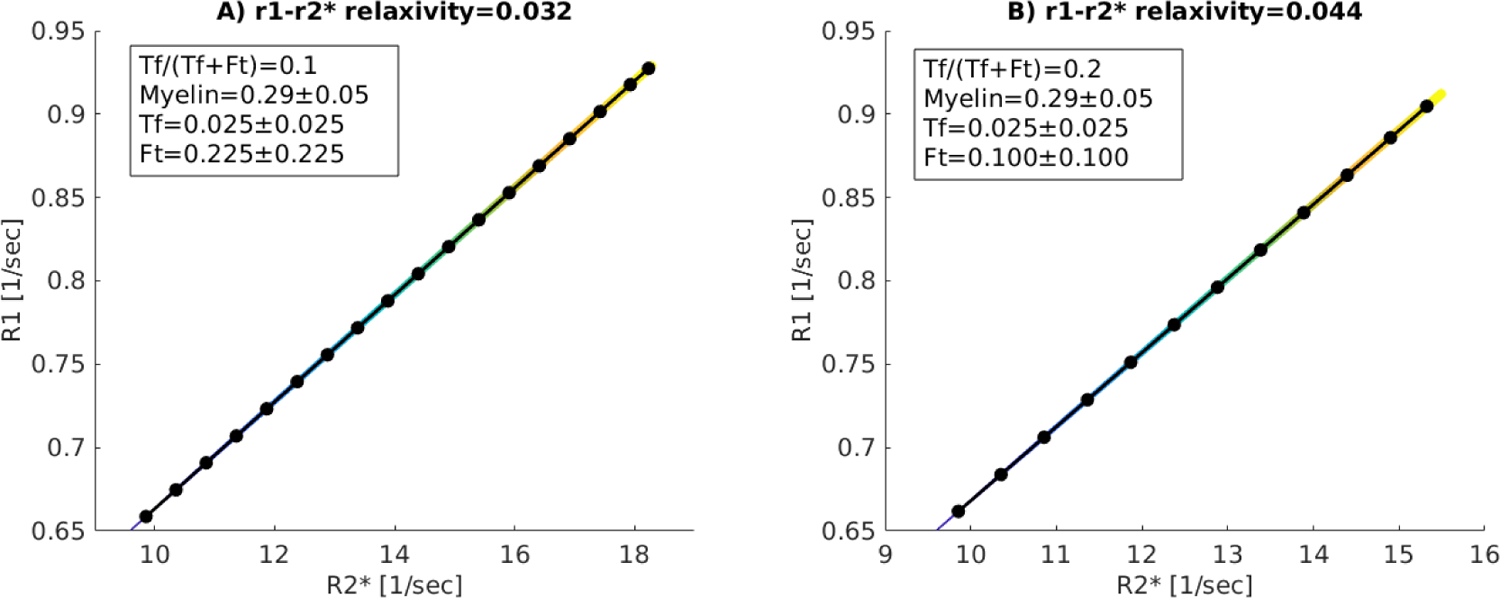
The r1-r2* relaxivity in two simulated ROIs with different transferrin-ferritin fractions (Tf/(Tf+Ft)) and a correlation between iron and myelin concentrations across voxels; (A) The transferrin-ferritin fraction is 0.1; (B) the transferrin-ferritin fraction is 0.2. In both A and B iron and myelin are correlated. Each figure shows the dependency of R1 on R2* for 1,000 representative simulated voxels. The colors of the data points indicate the variability in myelin concentration across voxels, and their sizes indicate the variability in iron compounds concentration across voxels (the simulated concentrations are shown in the text box, myelin is in units of [fraction] as MTV, transferrin and ferritin are in units of [mg/ml]). As in our in vivo pipeline, R2* and R1 values were binned (black data points represent the bins’ median), and a linear fit was calculated (black line). The slopes of the linear fit (shown in the title) represent the dependency of R1 on R2* (r1-r2* relaxivity) and change considerably with the transferrin-ferritin fraction even when iron and myelin are correlated.

## Supplementary Section 5: The r1-r2* relaxivity in the pallidum

The pallidum is unique in terms of its paramagnetic properties: it is highly rich in iron, but also contains iron oxides and metal depositions^3,89, 90^ which might affect the measurement of the r1-r2* relaxivity.

For young subjects, R1 and R2* values in the pallidum were the highest among all regions tested, and the r1-r2* relaxivity was the lowest (Figure 3). To make sure our results are not driven by the outlier values in the pallidum, we tried to exclude it from the comparisons between MRI and iron histology we show in Figure 3b-c. The correlation of the r1-r2* relaxivity with the iron concentration did not survive after excluding the pallidum (**Sup. Figure 31**). On the contrary, the correlations of R2* with the iron concentration and of the r1-r2* relaxivity with the iron mobilization remained significant even when the pallidum was excluded from the analysis (**Sup. Figure 31**).

Therefore, the correlation of the r1-r2* relaxivity with the iron concentration was driven mostly by the distinct behavior of the pallidum, while its correlation with the iron mobilization was stronger and more stable. These analyses demonstrate the enhanced sensitivity of the r1-r2* relaxivity to the iron homeostasis rather than to the absolute iron concentration.

**Sup. Figure 31:**
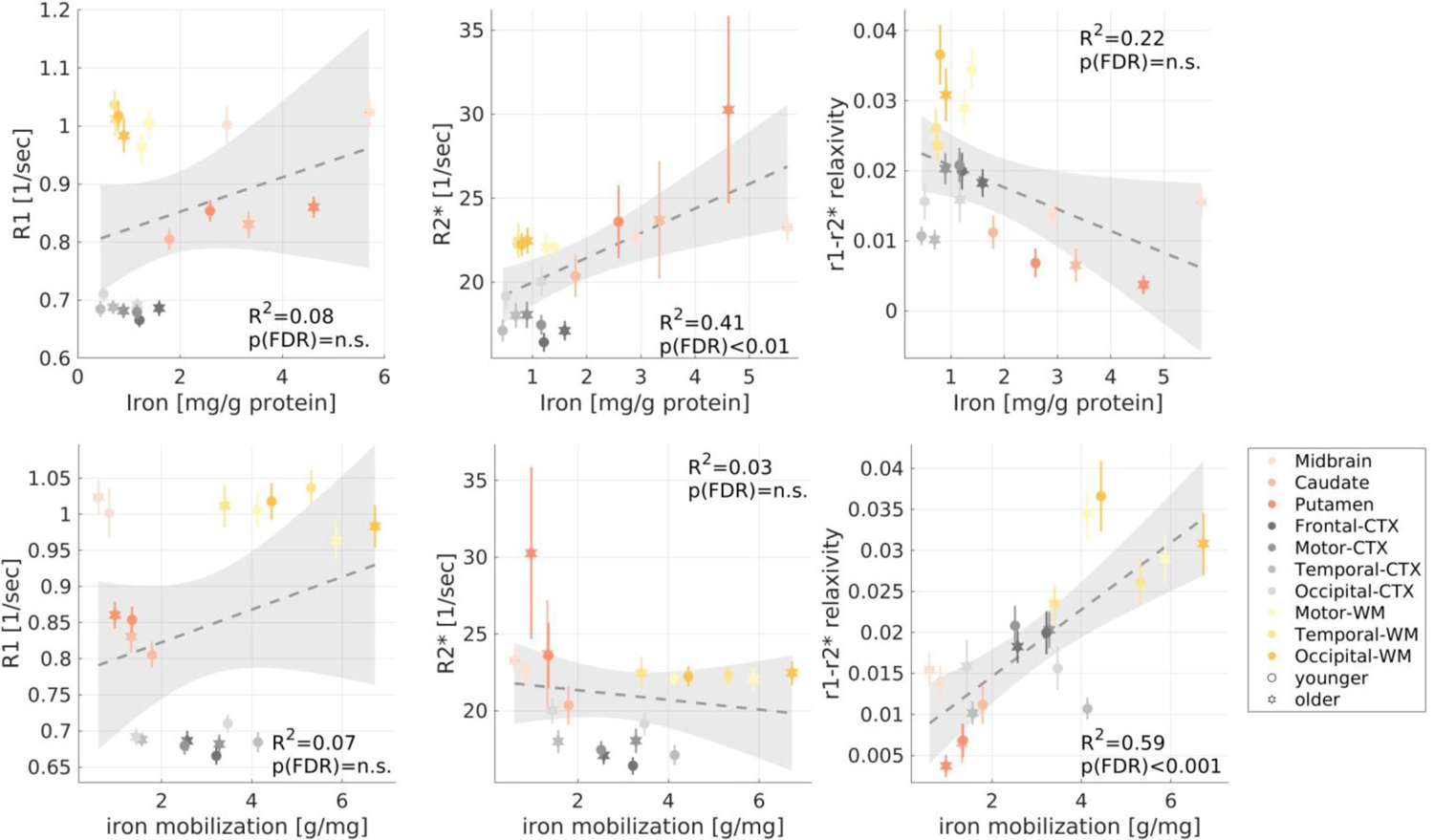
The correlations of MRI parameters with the iron environment when excluding the pallidum. **(a)** Replication of *Figure 3b-c* excluding the globus pallidus. The iron concentration and the iron mobilization capacity (transferrin/iron ratio) estimate postmortem (from the literature^5,7,9^) in different brain regions of younger (aged 27-64 years, N>=7) and older (aged 65-88 years, N>=8) subjects vs. R1, R2* and the r1-r2* relaxivity measured in vivo across younger (aged 23-63 years, N =26) and older (aged 65-77 years, N=13) subjects (different marker shapes) in 10 brain regions (different colors), excluding the pallidum. The correlation of the r1-r2* relaxivity with the iron concentration did not survive after excluding the pallidum. On the contrary, the correlations of R2* with the iron concentration and of the r1-r2* relaxivity with the iron mobilization remained significant.

## Supplementary methods

### R1-MTV dependency computation for phantoms

We computed the linear dependency of R1 on MTV across samples with varying iron-binding protein concentrations and liposomal fractions^76^. This process was implemented in MATLAB. We extracted the MTV values from all voxels and pooled them into 12 bins spaced equally between 0.05 and 0.40. This was done so that the linear fit would not be heavily affected by the density of the samples in different MTV regimes. The median MTV of each bin was computed, along with the median R1. We fitted the following linear model across samples:

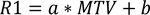

The slope of this linear model (a) represents the R1-MTV dependency. b is constant.

