## Supplemental information for "Non-invasive assessment of normal and impaired iron homeostasis in living human brains"

#### Supplementary Figure 1

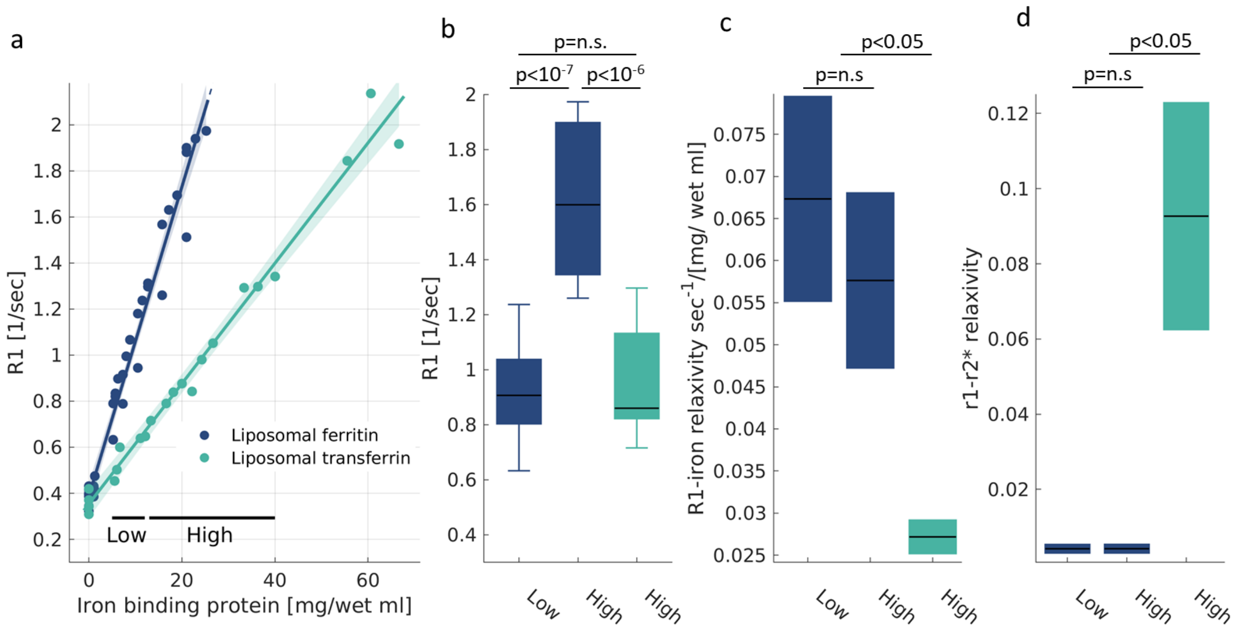

***Sup. Figure 1: The effect of iron concentration on different MR estimations. (a)*** *The dependency of R1 on the iron-binding protein concentration for liposomal ferritin and liposomal transferrin. Data points represent liposomal samples with varying iron-binding protein concentrations relative to the water fraction ([mg/wet ml]). The linear relationships between relaxation rates and iron-binding protein concentrations are marked by lines. The slopes of these lines are the iron relaxivities. Shaded areas represent the 95% confidence bounds.* ***(b)*** *The ambiguity in R1; R1 changes as a function of both iron environment and iron concentration. This is shown by calculating the median R1 value over samples with high and low iron-binding protein concentrations (marked in (a); concentration ranges were chosen so that the number of data points in each range is similar). We find that R1 is greater for a higher ferritin concentration than for a lower ferritin concentration, but also find that R1 is greater for ferritin than for transferrin. For each box, the  central line marks the median, the box extends vertically between the 25th and 75th percentiles, and the whiskers extend to the most extreme data points.* ***(c-d)*** *The ambiguity in R1 is resolved by the R1-iron relaxivity (c) and the r1-r2* (d), which are consistent when computed over higher or lower ferritin concentrations, and are consistently different from the iron relaxivity of transferrin regardless the concentration. For each box, the central lines marks the iron relaxivity, and the box shows the 95% confidence bounds of the linear fit. p-values are for the ANCOVA test corrected for multiple comparisons.*

#### Supplementary Section 1: The dependency of R1 and R2* on the iron concentration.

In ***Figure 1*** we computed the iron relaxivity as the dependency of R1 and R2* on the concentration of iron-binding proteins. We showed that different iron environments have different relaxivities. However, different proteins bind different amounts of iron. For example, ferritin binds three orders of magnitude more iron ions than does transferrin[3]. Therefore, we wanted to exclude the possibility that the different iron ion concentrations drive the different relaxivities of ferritin and transferrin.

We verified that the relaxivity changes according to the molecular type of iron, even when accounting for discrepancies in iron loading. We estimated the iron ion concentrations for ferritin and transferrin (see Methods section "Estimation of total iron content in phantoms") and tested whether those values can explain their different iron relaxivities. Importantly, after computing the iron relaxivity as the dependency of relaxation rates on the iron ion concentrations (rather than the concentration of iron-binding proteins), we still find that different iron environments have distinct relaxivities (**Sup. Figure 2**, $p(ANCOVA)<10^{-39}$).

To further stress the sensitivity of the r1-r2* relaxivity to the iron environment, we compared the relaxivity of liposomal ferrous iron ($Fe^{2+}$) and iron bound to liposomal transferrin (**Sup. Figure 3**). Unlike ferritin and transferrin, these two iron compounds have relatively similar iron ion concentrations. Yet we find that they produce different iron relaxivities ($p(ANCOVA)<10^{-4}$). The r1-r2* relaxivities of these two iron environments are different as well ($p(ANCOVA )<0.05$). Therefore, the iron relaxivity and the r1-r2* relaxivity are changing as a function of the molecular iron environment, even when accounting for the differences in iron binding.

#### Supplementary Figure 2

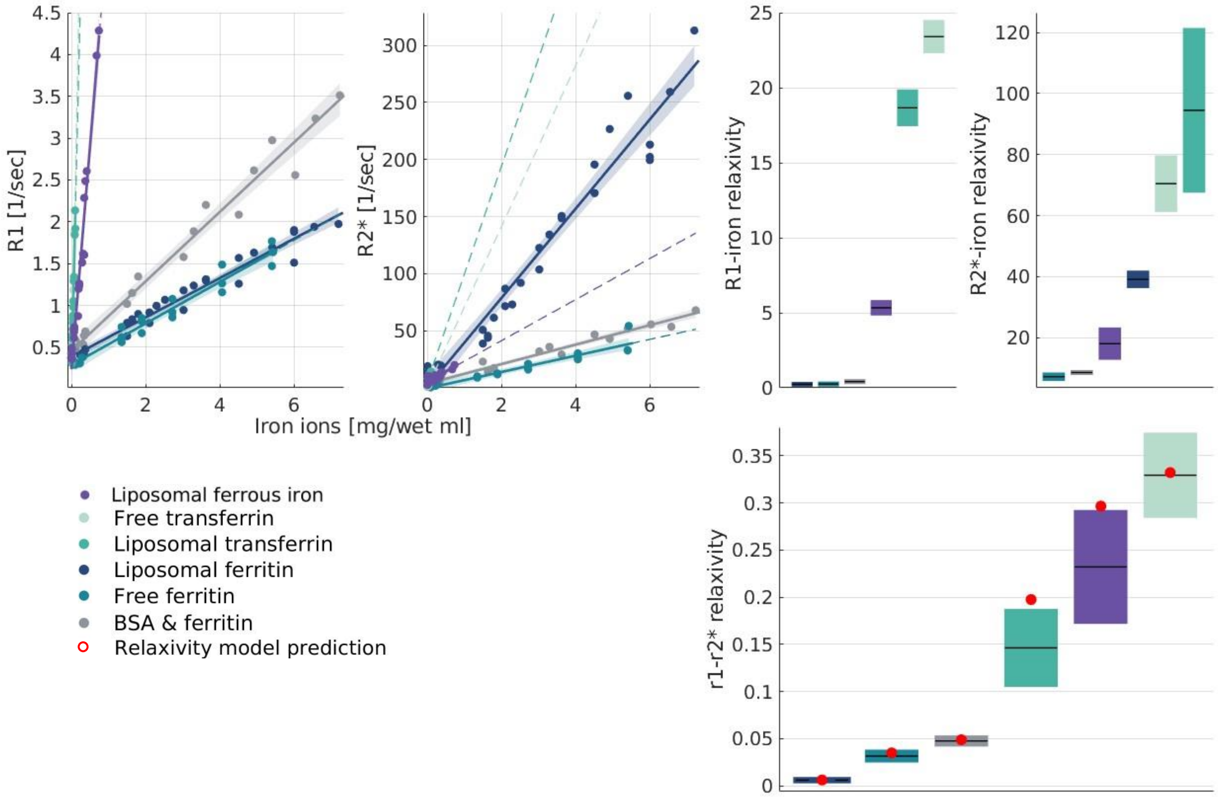

***Sup. Figure 2: The iron relaxivity and the r1-r2* relaxivity are sensitive to the molecular type of iron regardless of the differences in iron-binding. (a-b)*** *The dependency of R1 and R2* on the estimated iron concentration (see method section "Estimation of total iron content in phantoms") for six different iron compounds: free ferritin, liposomal-ferritin, Bovine Serum Albumin (BSA)-ferritin mixture, free transferrin, liposomal transferrin and liposomal ferrous iron. Data points represent samples with varied estimated iron ion concentrations relative to the water fraction ([mg/wet ml]). The linear relationships between relaxation rates and iron concentration are marked by lines. The slopes of these lines are the iron relaxivities. Dashed lines represent extrapolations the linear fits, and shaded areas represent the 95% confidence bounds.* ***(c)*** *The iron relaxivities of R1 and R2* are different for different iron environments (p(ANCOVA)<$10^{39}$). Iron relaxivity is calculated here based on the estimated iron concentration (and not iron-binding proteins concentrations, as in* Figure 1*). To do so, we use the slope of the linear relationships shown in (a,b), expressed in [sec-1/(mg/wet ml)]. For each box the central lines marks the iron relaxivity, and the box shows the 95% confidence bounds of the linear fit.* ***(d)*** *The theoretical model successfully predicts the r1-r2* relaxivity even when it is based on the estimated iron ions concentration (and not iron compound concentrations, as in* Figure 1*). The model's prediction is based on the ratio between the iron relaxivities of R1 and R2* as shown in (c). For each box the central line marks the r1-r2* relaxivity, and the box shows the 95% confidence bounds of the linear fit. Red dots represent the prediction of the theoretical model.*

#### Supplementary Figure 3

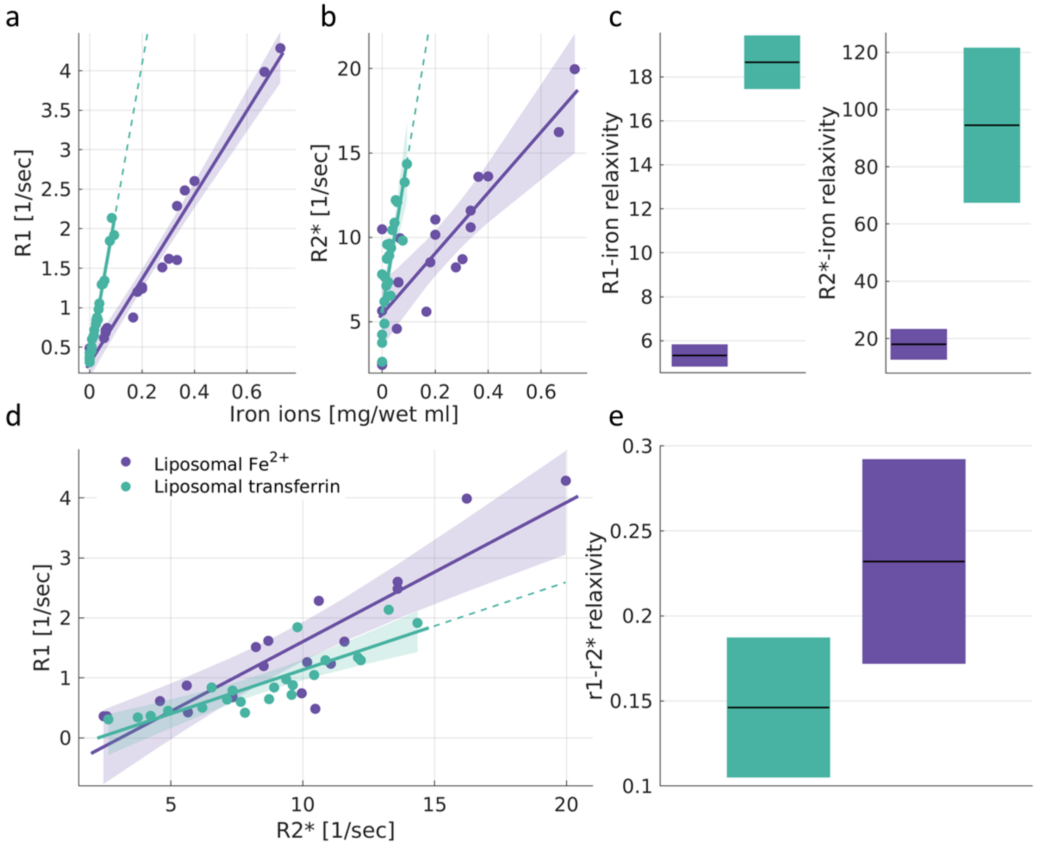

***Sup. Figure 3: The iron relaxivity and the r1-r2* relaxivity are sensitive to the molecular iron environment even when the iron concentration is similar. (a-b)*** *The dependency of R1 and R2* on the estimated iron concentration for two different iron environments: liposomal transferrin (purple) and liposomal $Fe^{2+}$ (green). Data points represent liposomal samples with varying iron ion concentrations relative to the water fraction ([mg/wet ml]). The linear relationships between relaxation rates and iron ion concentration are marked by lines. The slopes of these lines are the iron relaxivities. Dashed lines represent extrapolations of the linear fits. Shaded areas represent the 95% confidence bounds.* ***(c)*** *The iron relaxivity of R1 and R2* is different for different iron environments ($p(ANCOVA)<10^{-4}$). Iron relaxivity is calculated by taking the slope of the linear relationships shown in (a,b), and is measured in [sec-1/(mg/wet ml)]. For each box, the central line marks the iron relaxivity, and the box shows the 95% confidence bounds of the linear fit.* ***(d)*** *The dependency of R1 on R2* for different iron environments. Data points represent samples with varying concentrations. The linear relationships between R1 and R2* are marked by lines. The slopes of these lines are the r1-r2* relaxivities. Dashed lines represent extrapolations of the linear fits. Shaded areas represent the 95% confidence bounds.* ***(e)*** *The r1-r2* relaxivity is different for different iron environments (p(ANCOVA )<0.05). For each box, the central line marks the r1-r2* relaxivity, and the box shows the 95% confidence bounds of the linear fit.*

### Supplementary Figure 4

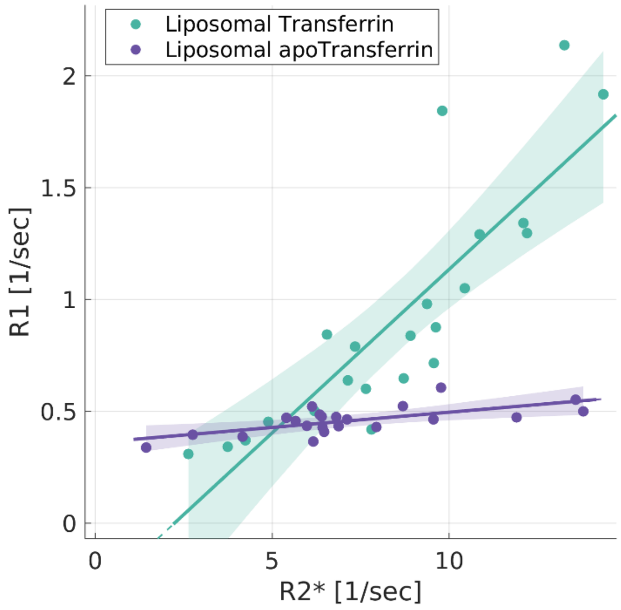

***Sup. Figure 4: Validating the sensitivity of the the r1-r2* relaxivity to the paramagnetic properties of transferrin.*** *Data points represent liposomal samples with varying concentrations of transferrin (green) and apo-transferrin (transferrin which is not bound to iron, in purple). The linear relationships between R1 and R2* are marked by lines. The slopes of these lines are the r1-r2* relaxivities. Shaded areas represent the 95% confidence bounds. Apo-transferrin with no iron has lower r1-r2* relaxivity compared to iron-bound transferrin (p(ANCOVA)<$10^{-8}$). Therefore, the r1-r2* relaxivity is sensitive to the paramagnetic properties of iron-binding proteins and not to the proteins themselves.*

#### Supplementary Figure 5

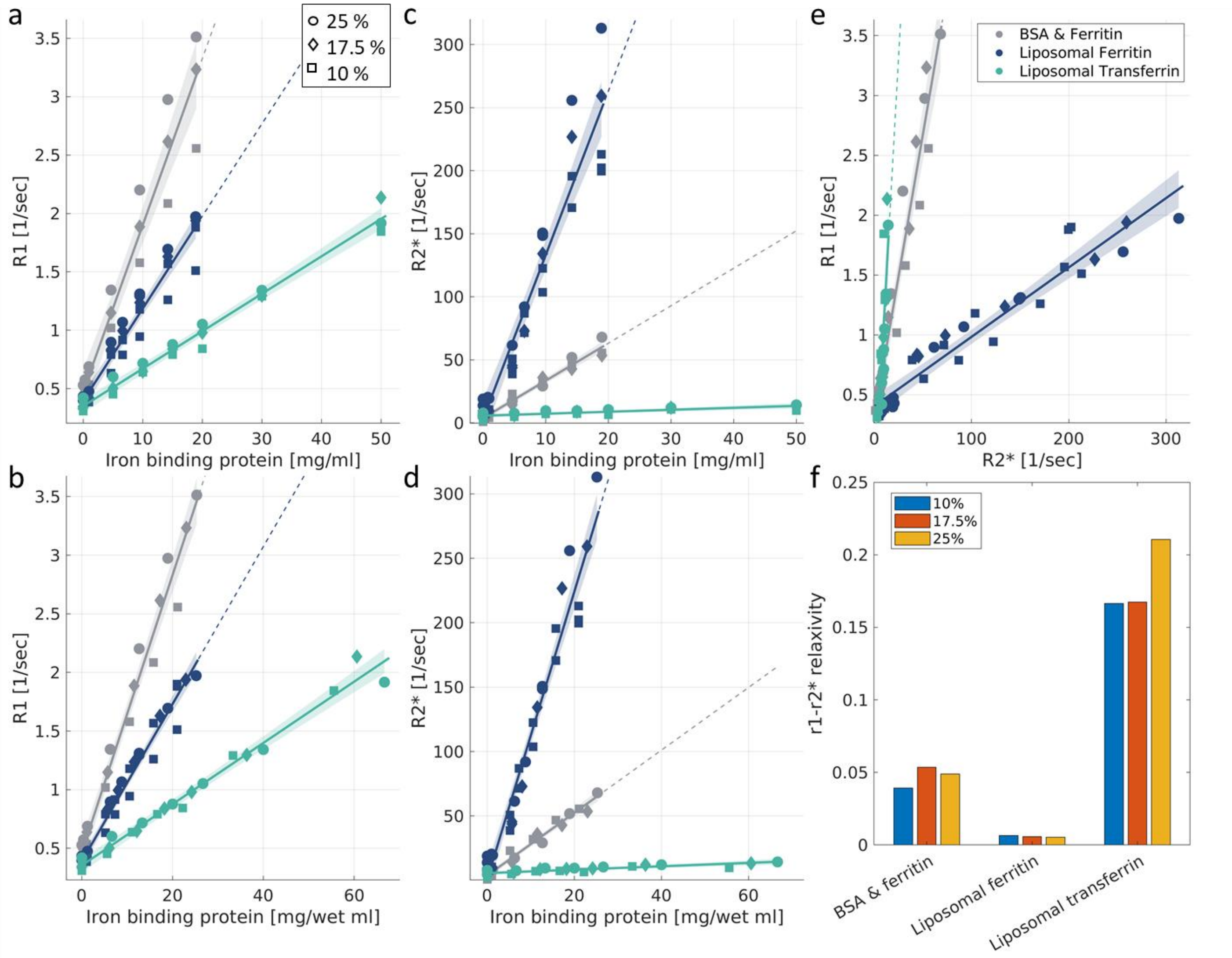

***Sup. Figure 5: Relaxivities are stable across liposomal or BSA fractions. (a)*** *The dependency of R1 on the iron-binding protein concentration for different liposomal (or BSA) fractions (different symbols) and different iron environments (different colors). The x-axis represents the absolute concentration of iron-binding proteins (not relative to the water concentration, as in b). The linear relationships between relaxation rates and iron-binding protein concentration are marked by lines. The slopes of these lines are defined as the iron relaxivities. R1 values are affected by the variable liposomal (or BSA) fractions, but the iron relaxivities of different iron environments are still distinct, regardless of this manipulation. Dashed lines represent extrapolations of the linear fits. Shaded areas represent the 95% confidence bounds.* ***(b)*** *The dependency of R1 on the iron-binding protein concentration for different liposomal (or BSA) fractions (different symbols) and different iron environments (different colors). Here the x-axis represents the concentration of iron-binding proteins relative to the water fraction (which varies with the liposomal or BSA fraction). This estimation, in units of [mg/wet ml], further eliminates the effect of the liposomal (or BSA) fraction on the iron relaxivities. This is evident by the alignment of the data points with different liposomal (or BSA) fractions (different symbols) along the iron relaxivity linear fit.* ***(c-d)*** *A similar analysis for the R2*-iron relaxivity. The effect of the different liposomal (or BSA) fractions on the*

*R2*-iron relaxivity is eliminated by the calculation of the iron-binding proteins concentration relative to the water fraction ([mg/wet ml]).* ***(e)*** *The dependency of R1 on R2* for different liposomal (or BSA) fractions (different symbols) and different iron environments (different colors). The r1-r2* relaxivities of different iron environments are distinct even when calculated across liposomal (or BSA) fractions.* ***(f)*** *The r1-r2* relaxivity (y-axis) for different compounds of iron (liposomal ferritin, liposomal transferrin and BSA-ferritin mixture) in three different liposomal (or BSA) fractions (colors). The differences in the r1-r2* relaxivity between iron environments are greater than the differences within each iron environment for the variable liposomal (or BSA) fractions.*

#### Supplementary Figure 6

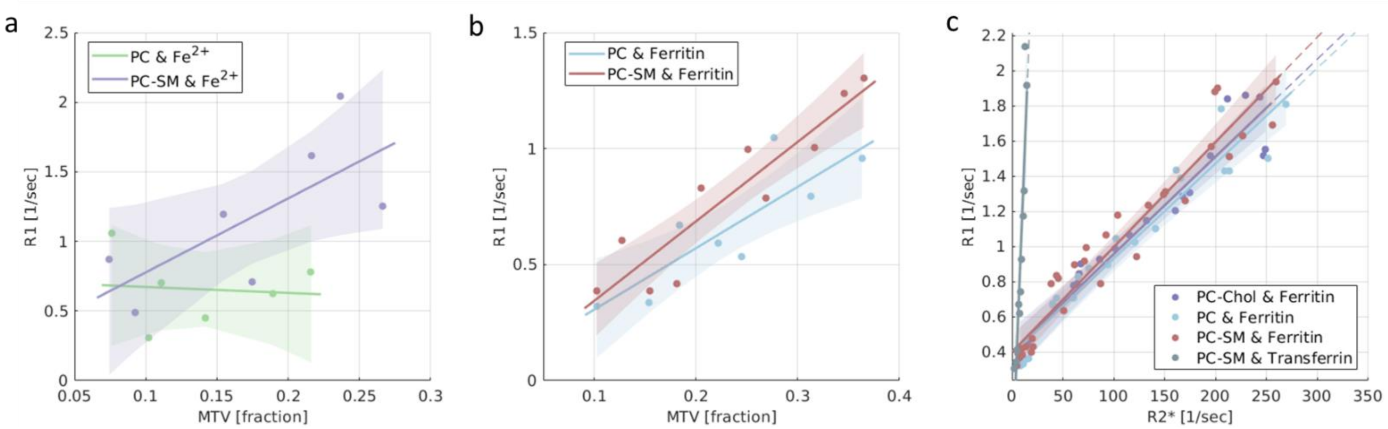

***Sup. Figure 6: The r1-r2* relaxivity is stable for different types of lipids, while the R1-MTV dependency is sensitive to the lipid type. (a)*** *The dependency of R1 on MTV for an iron ion compound ($Fe^{2+}$) mixed with two different lipids: phosphatidylcholine (PC, green) and a mixture of PC-sphingomyelin (PC-SM, blue). This result replicates the sensitivity of the MTV dependencies to lipid types[76] in $Fe^{2+}$-containing phantoms.* ***(b)*** *The dependency of R1 on MTV for a second iron compound (ferritin) mixed with the same two lipids (PC and PC-SM).* ***(c)*** *The dependency of R1 on R2* (r1-r2* relaxivity) for four different iron-lipid mixtures: ferritin-PC ferritin-PC-SM, transferrin-PC-SM and ferritin-PC-cholesterol (PC-Chol, blue). The r1-r2* relaxivity is similar for the different lipid types mixed with ferritin, and the main difference is between the iron binding proteins; i.e., transferrin sample and the ferritin samples.*

### Supplementary Figure 7

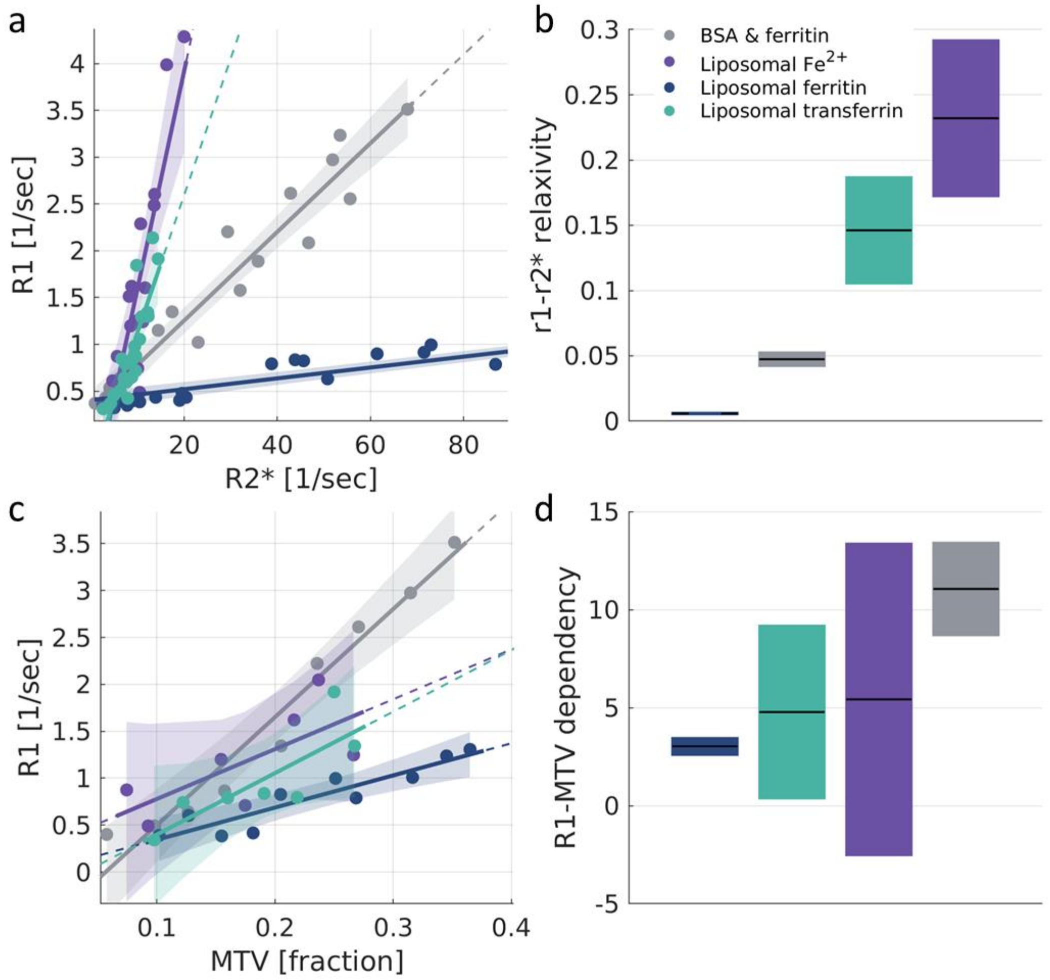

***Sup. Figure 7: Iron environments are less distinguishable with MTV dependencies than with the r1-r2* relaxivity.*** ***(a)*** *The dependency of R1 on R2* (r1-r2* relaxivity) for different iron environments: liposomal-ferritin, BSA-ferritin mixture, liposomal transferrin and liposomal $Fe^{2+}$. Liposomal samples are based on PC-sphingomyelin. Data points represent samples with varying iron compounds concentrations relative to the water fraction. The linear relationships between relaxation rates are marked by lines, whose slopes represent the r1-r2* relaxivities. Dashed lines represent extrapolations of the linear fits. Shaded areas represent the 95% confidence bounds. The x-axis presents only partial range of R2* values, similar to Figure 1d (for the entire R2* range, see the inset of Figure 1d).* ***(b)*** *The r1-r2* relaxivities are different for these four iron environments. For each box, the central line marks the r1-r2* relaxivity, and the box shows the 95% confidence bounds of the linear fit.* ***(c)*** *The dependency of R1 on MTV for these four iron environments. Data points represent samples with varying iron compounds concentrations relative to the water fraction. The linear relationships between R1 and MTV are marked by lines, whose slopes represent the R1-MTV dependencies. Dashed lines represent extrapolations of the linear fits. Shaded areas represent the 95% confidence bounds.* ***(d)*** *The R1-MTV dependencies for the four iron environments. For each box, the central line marks the R1-MTV dependency, and the box showes the 95% confidence bounds of the linear fit.*

#### Supplementary Figure 8

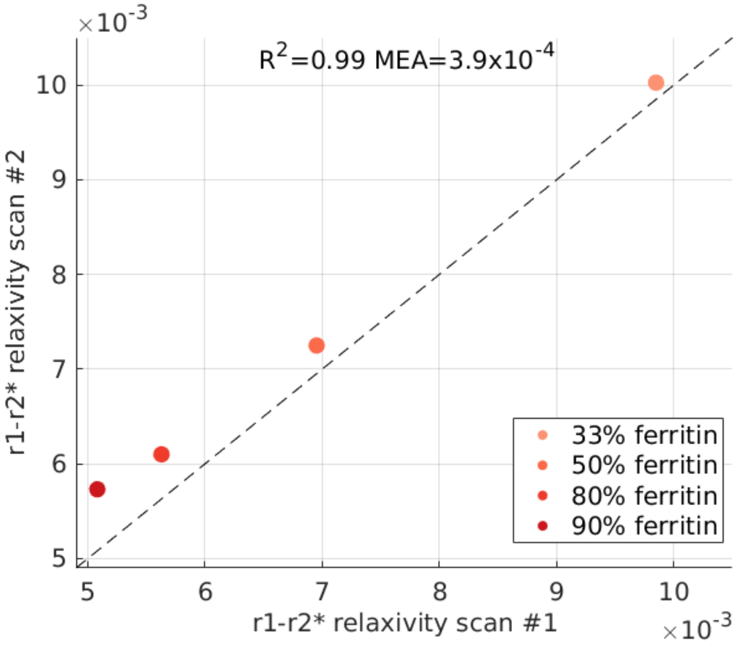

***Sup. Figure 8: The reproducibility of the r1-r2* relaxivity measurement in vitro.*** *The reproducibility of the r1-r2* relaxivity measurement for in vitro ferritin-transferrin mixtures was estimated based on scan-rescan experiments. Four different transferrin-ferritin mixtures were scanned twice (on different days). Each mixture experiment had a different transferrin-ferritin fraction (different colors, legend shows the percentage of ferritin in the mixture). The r1-r2* relaxivity of each mixture experiment was calculated over samples with the same transferrin-ferritin fraction but varying total iron-binding protein concentrations. Figure shows the r1-r2* relaxivity values measured in the first scan (x-axis) vs. the r1-r2* relaxivity values measured in the second scan (y-axis) for each in vitro experiment. Dashed line is the identity line. The measured scan-rescan mean absolute error (MAE) represents an experimental estimate of the detection limit of the r1-r2* relaxivity.*

#### Supplementary Figure 9

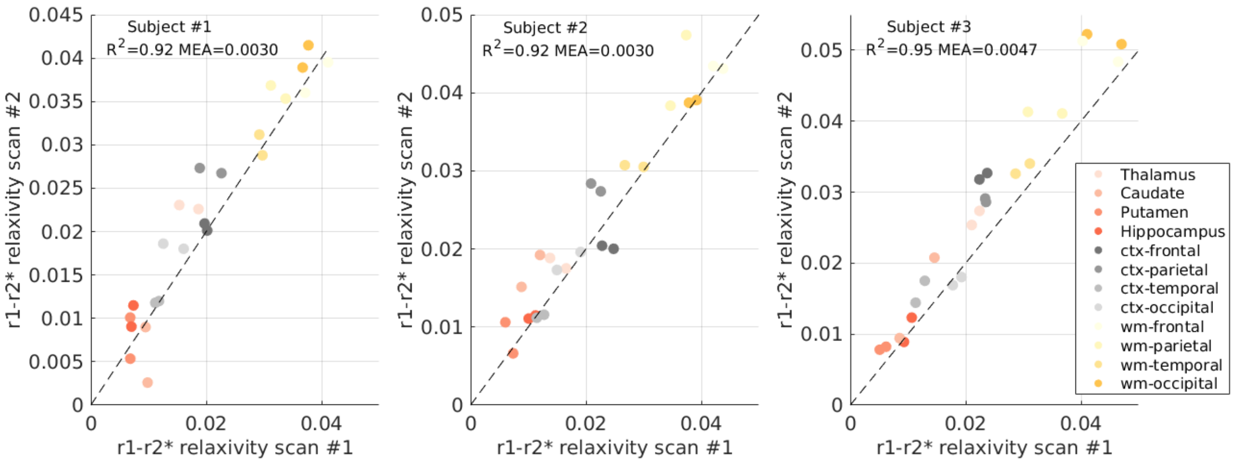

***Sup. Figure 9: The reproducibility of the r1-r2* relaxivity measurement in the in vivo brain.*** *The reproducibility of the r1-r2* relaxivity measurement in the in vivo brain was estimated based on scan-rescan experiments in three human subjects. Each subject was scanned twice in the MRI (on different days). The r1-r2* relaxivity was calculated for each scan in 12 different brain regions (different colors) in both hemispheres. Panels show the r1-r2* relaxivity values measured in the first scan (x-axis) vs. the r1-r2* relaxivity values measured in the scanned scan for each subject. Dashed line is the identity line. The measured scan-rescan mean absolute error (MAE) represents an experimental estimate of the detection limit of the r1-r2* relaxivity.*

#### Supplementary Figure 10

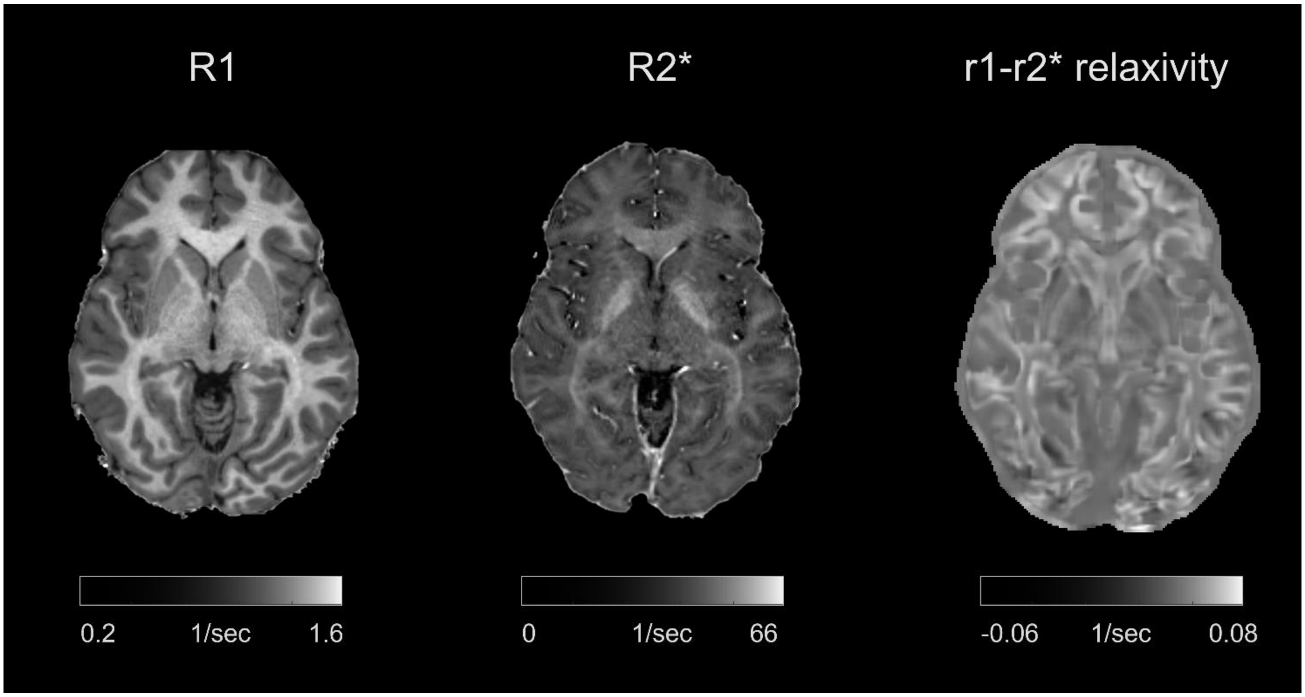

***Sup. Figure 10: Voxel-wise comparison of the r1-r2* relaxivity map to R1 and R2* maps in the in vivo healthy brain.*** *The maps of R1 (left) and R2* (middle) are compared to the local r1-r2* relaxivity visualization (right) on a representative young healthy subject. The voxel-wise visualization of the r1-r2* relaxivity in the brain was generated based on the local linear dependency of R1 on R2* using a moving-window approach (for more details see "Generating voxel-wise r1-r2* relaxivity visualizations" in Methods).*

#### Supplementary Figure 11

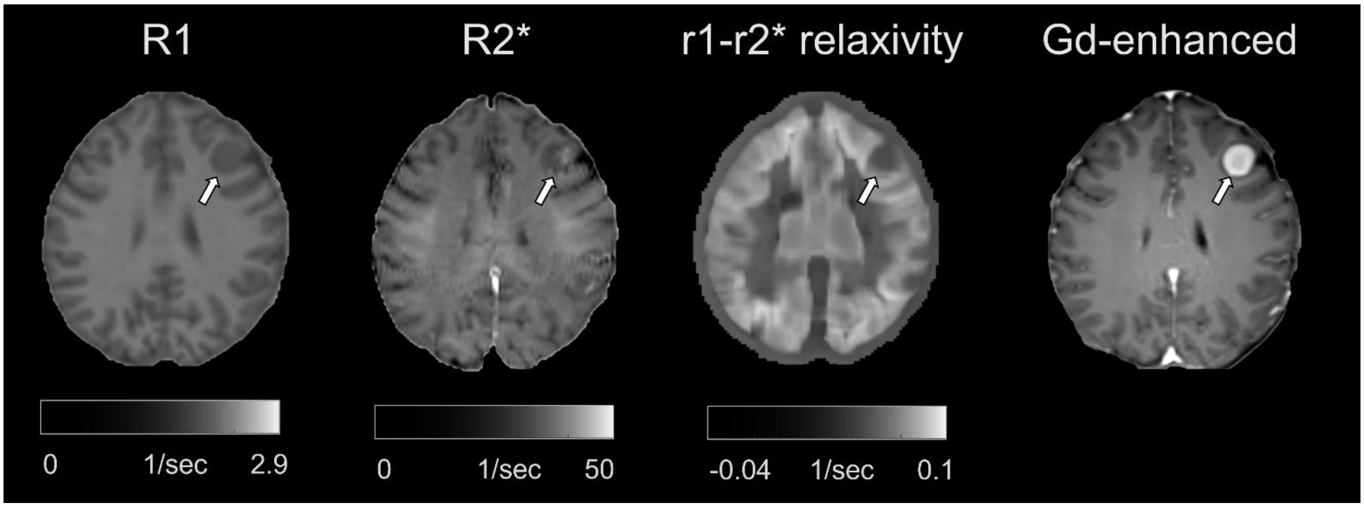

***Sup. Figure 11: Voxel-wise comparison of the r1-r2* relaxivity map to R1 and R2* maps and to the Gd-enhanced contrast in the in vivo brain of a meningioma patient.*** *Representative visualization of R1, R2*, the voxel-wise r1-r2* relaxivity map and the Gd-enhanced contrast for a meningioma patient. Tumors are marked with arrows. The voxel-wise r1-r2* relaxivity map was generated based on the local linear dependency of R1 on R2* using a moving-window approach (for more details see "Generating voxel-wise r1-r2* relaxivity visualizations" in Methods).*

#### Supplementary Figure 12

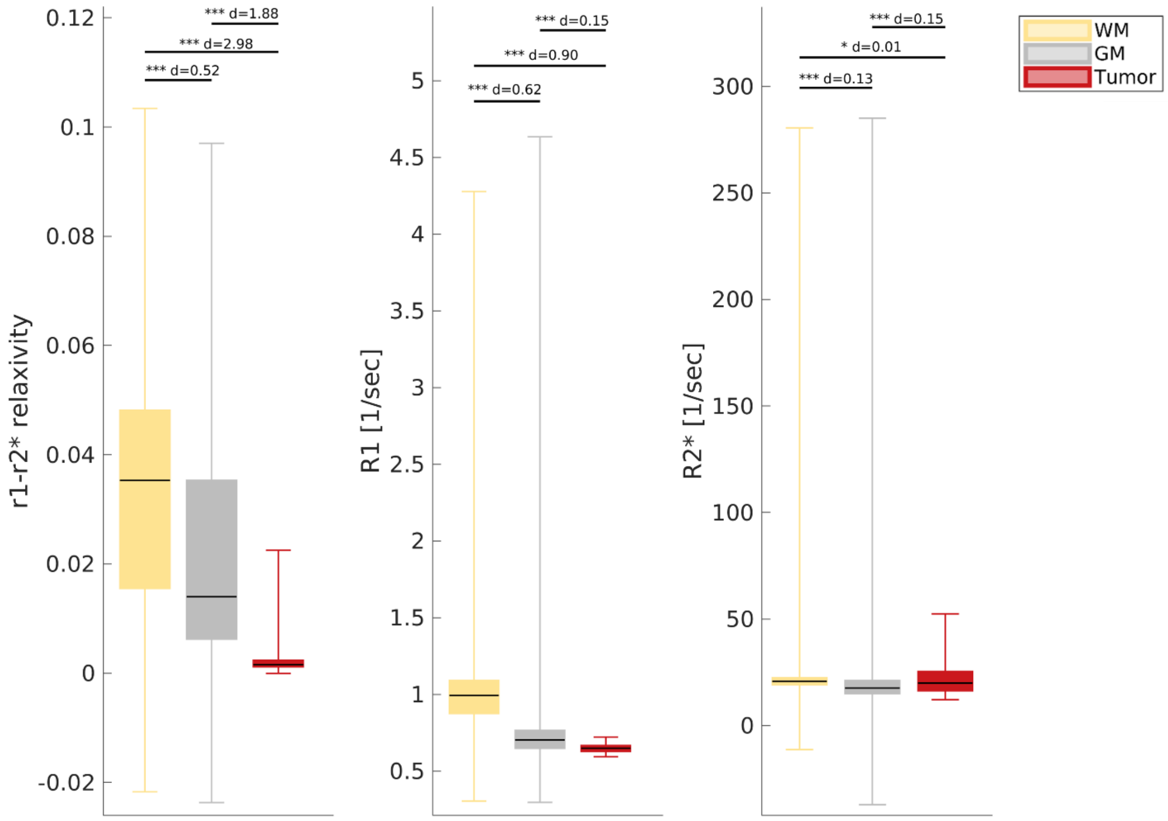

***Sup. Figure 12: The voxel-wise r1-r2* relaxivity enhances the contrast between tumor tissue and non-pathological tissue on a representative meningioma patient.*** *Replication of the ROI-based results presented in figure 4d-f on the voxel-wise level. The contrast between the white matter (WM), gray matter (GM) and tumor tissues is presented for R1, R2* and the voxel-wise r1-r2* relaxivity. The variation in each box is calculated across voxels in a representative meningioma patient (Sup. Figure 11). The 25th, 50th and 75th percentiles and extreme data points are shown. The d-values represent the effect size (Cohen's d) of the differences between tissue types, and the significance level is based on a t-test. Across voxels, the r1-r2* relaxivity allows to distinguish between tumor tissue and non-pathological tissue better than R1 and R2*. Estimates in non-pathological tissues are for the tumor-free hemisphere.* *$p < 0.05$; **$p < 0.01$; ***$p < 0.001$*

#### Supplementary Figure 13

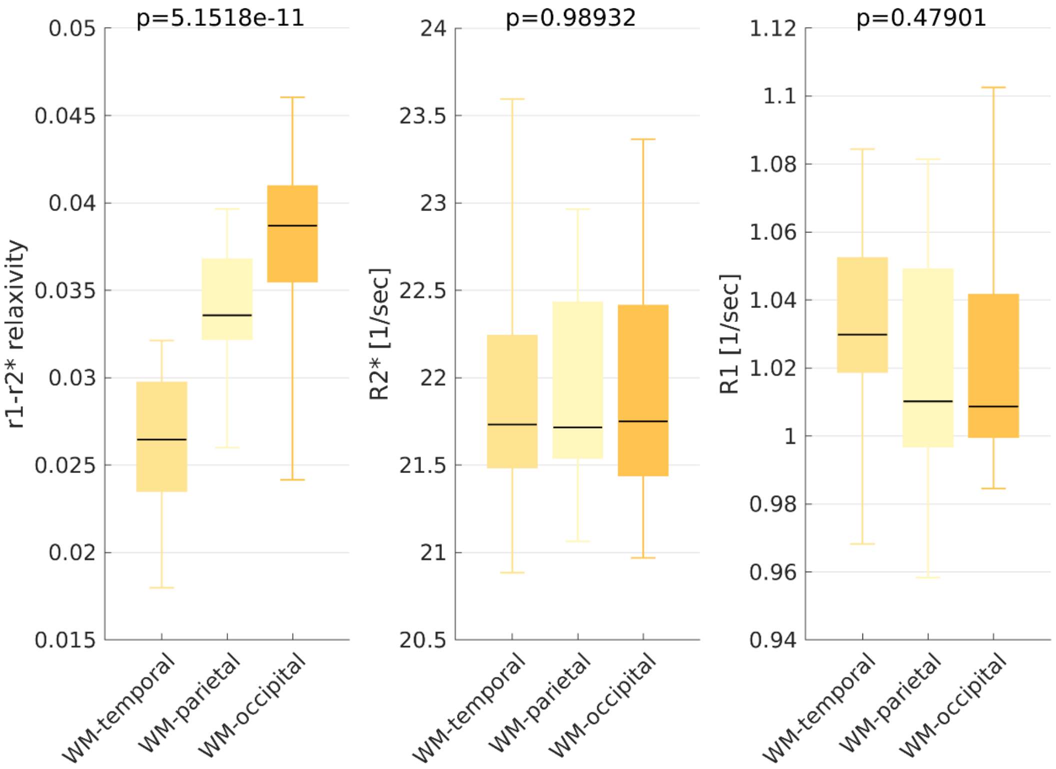

***Sup. Figure 13: The new contrast of the r1-r2* relaxivity in white matter.*** *From left to right, r1-r2* relaxivity, R2* and R1 for three different white-matter (WM) ROIs (temporal, parietal and occipital). We can separate these three WM ROIs with the r1-r2* relaxivity but not with either R2* or R1. Boxes represent the variation in the MRI parameters across normal subjects (age 27±2, N = 21). The 50$^{th}$ percentile (horizontal black lines) 25th and 75th percentiles (box edges) and extreme data points (whiskers) are shown for each box. p-values are for the ANOVA test.*

#### Supplementary Figure 14

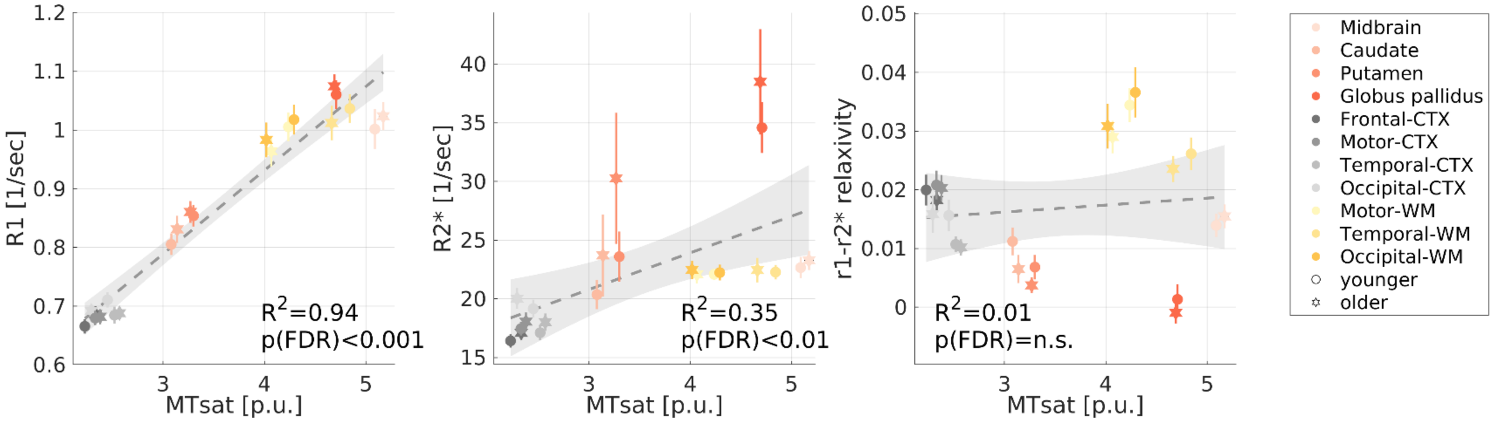

***Sup. Figure 14: The correlations of MRI parameters with MTsat.*** *The qMRI measurement of the magnetization transfer saturation (MTsat) vs. R1, R2* and the r1-r2* relaxivity measured in vivo across younger (aged 23-63 years, N =26) and older (aged 65-77 years, N=13) subjects (different marker shapes) in 10 brain regions (different colors). Unlike R1 and R2*, the r1-r2* relaxivity is not significantly correlated with MTsat.*

#### Supplementary Figure 15

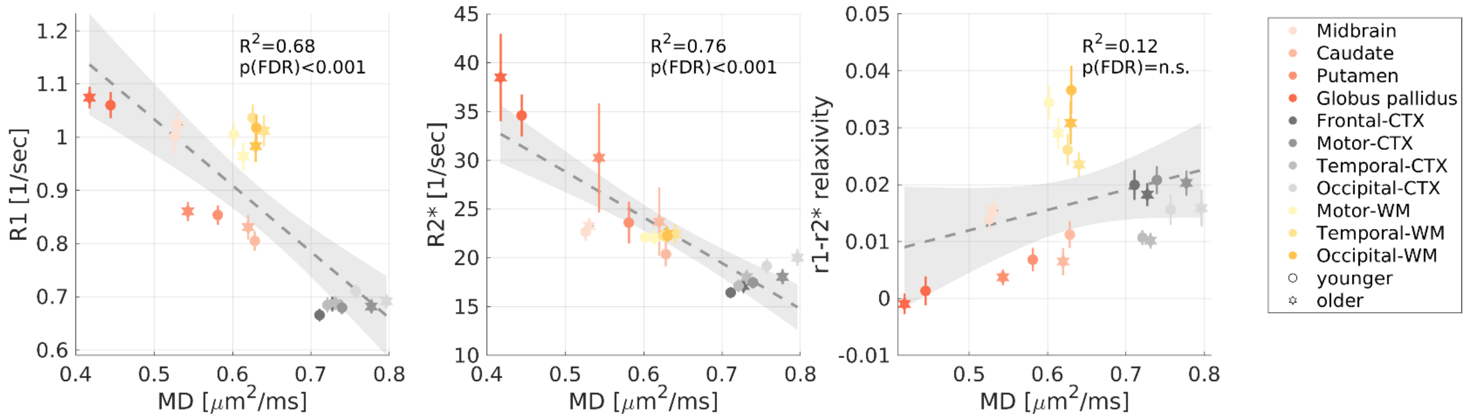

***Sup. Figure 15: The correlations of MRI parameters with MD.*** *The qMRI measurement of the mean diffusivity (MD) measured in vivo across younger (aged 23-63 years, N =25) and older (aged 65-77 years, N=12) subjects vs. R1, R2* and the r1-r2* relaxivity measured in vivo across younger (aged 23-63 years, N =26) and older (aged 65-77 years, N=13) subjects (different marker shapes) in 10 brain regions (different colors). Unlike R1 and R2*, the r1-r2* relaxivity is not significantly correlated with MD.*

#### Supplementary Figure 16

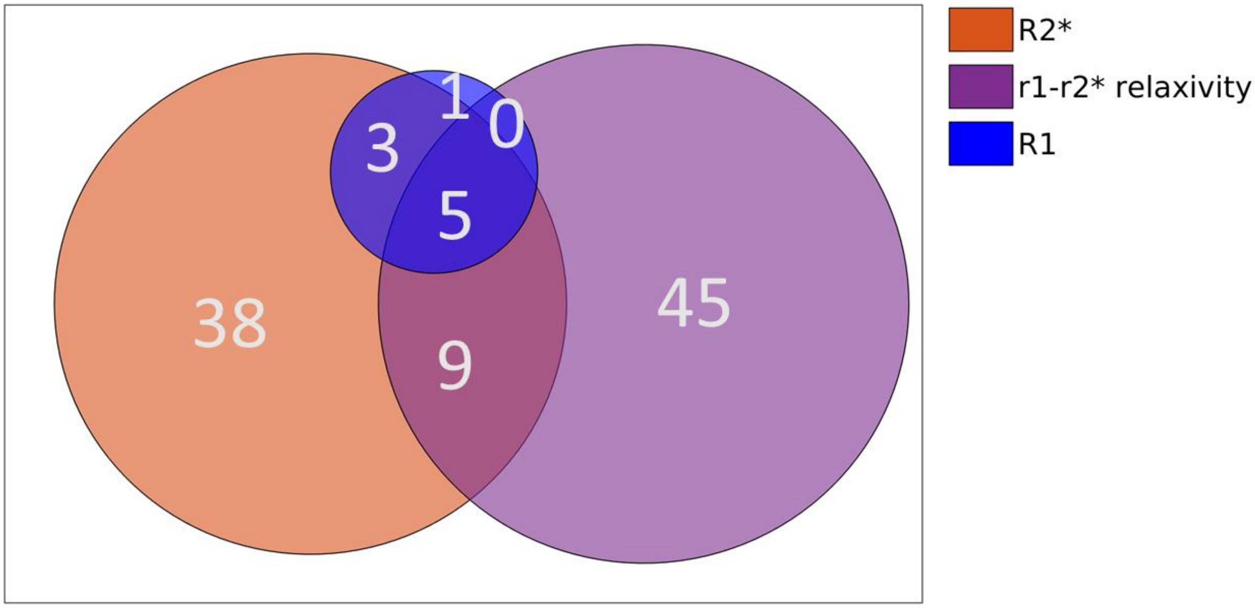

***Sup. Figure 16: The number of significantly enriched pathways associated with each qMRI parameter.*** *The Venn diagram shows the number of significantly enriched pathways (p(FWER)<0.01) for each qMRI parameter (R2*, R1 and the r1-r2* relaxivity). Almost half of the significantly enriched pathways are exclusive for the r1-r2* relaxivity. See also supplementary table 1.*

### Supplementary Table 1:

| pathway | R1 NES | R1 pval | R2* NES | R2* pval | r1-r2* relaxivity NES | r1-r2* relaxivity pval |
|---|---|---|---|---|---|---|
| GO ALPHA BETA T CELL ACTIVATION | 1.15 | 1 | 2.43 | 0.001 | -1.12 | 1 |
| GO ALPHA BETA T CELL DIFFERENTIATION | 1.26 | 1 | 2.44 | 0.001 | -1.04 | 1 |
| GO ANTIGEN BINDING | 0.79 | 1 | 2.23 | 0.024 | -2.86 | 0 |
| GO B CELL MEDIATED IMMUNITY | -0.64 | 1 | 1.48 | 1 | -2.56 | 0 |
| GO B CELL RECEPTOR SIGNALING PATHWAY | 0.97 | 1 | 2.21 | 0.038 | -2.50 | 0 |
| GO CD4 POSITIVE ALPHA BETA T CELL ACTIVATION | 0.99 | 1 | 2.32 | 0.005 | -1.22 | 1 |
| GO CD4 POSITIVE ALPHA BETA T CELL DIFFERENTIATION | 1.22 | 1 | 2.38 | 0.003 | -1.20 | 1 |
| GO COMPLEMENT ACTIVATION | -0.86 | 1 | 1.33 | 1 | -2.90 | 0 |
| GO CONDENSED NUCLEAR CHROMOSOME CENTROMERIC REGION | -2.33 | 0.004 | -1.64 | 1 | -1.46 | 1 |
| GO COTRANSLATIONAL PROTEIN TARGETING TO MEMBRANE | 2.55 | 0.001 | 2.44 | 0.001 | 3.18 | 0 |
| GO CYTOSOLIC LARGE RIBOSOMAL SUBUNIT | 2.31 | 0.019 | 2.27 | 0.012 | 2.95 | 0 |
| GO CYTOSOLIC RIBOSOME | 2.21 | 0.112 | 2.57 | 0 | 3.06 | 0 |
| GO CYTOSOLIC SMALL RIBOSOMAL SUBUNIT | 1.73 | 1 | 2.38 | 0.003 | 2.44 | 0.003 |
| GO DEFENSE RESPONSE TO BACTERIUM | 0.78 | 1 | 1.76 | 1 | -2.27 | 0.006 |
| GO ESTABLISHMENT OF PROTEIN LOCALIZATION TO ENDOPLASMIC RETICULUM | 2.11 | 0.317 | 2.08 | 0.272 | 3.14 | 0 |
| GO FC RECEPTOR MEDIATED STIMULATORY SIGNALING PATHWAY | 1.29 | 1 | 1.65 | 1 | -2.29 | 0.004 |
| GO HUMORAL IMMUNE RESPONSE | 0.72 | 1 | 1.72 | 1 | -2.43 | 0 |
| GO HUMORAL IMMUNE RESPONSE MEDIATED BY CIRCULATING IMMUNOGLOBULIN | -0.52 | 1 | 1.61 | 1 | -3.10 | 0 |
| GO IMMUNE RECEPTOR ACTIVITY | 1.27 | 1 | 2.46 | 0 | 0.71 | 1 |
| GO IMMUNOGLOBULIN COMPLEX | -1.14 | 1 | 2.52 | 0 | -3.62 | 0 |
| GO IMMUNOGLOBULIN COMPLEX CIRCULATING | 0.94 | 1 | 2.32 | 0.005 | -3.14 | 0 |
| GO IMMUNOGLOBULIN RECEPTOR BINDING | 0.72 | 1 | 2.30 | 0.008 | -3.06 | 0 |
| GO KINETOCHORE | -2.00 | 0.594 | -2.40 | 0.002 | -0.94 | 1 |
| GO LARGE RIBOSOMAL SUBUNIT | 0.91 | 1 | 1.13 | 1 | 2.59 | 0 |
| GO METAPHASE ANAPHASE TRANSITION OF CELL CYCLE | -2.10 | 0.219 | -2.55 | 0 | -0.81 | 1 |
| GO MITOCHONDRIAL GENE EXPRESSION | -2.18 | 0.067 | -2.45 | 0 | 1.57 | 1 |
| GO MITOCHONDRIAL TRANSLATION | -2.08 | 0.266 | -2.44 | 0 | 1.82 | 0.998 |
| GO MITOCHONDRIAL TRANSLATIONAL TERMINATION | -2.05 | 0.388 | -2.58 | 0 | 1.73 | 1 |
| GO MITOTIC METAPHASE PLATE CONGRESSION | -1.80 | 1 | -2.41 | 0.002 | 1.05 | 1 |
| GO MITOTIC NUCLEAR DIVISION | -2.12 | 0.156 | -2.32 | 0.009 | -0.85 | 1 |
| GO MITOTIC SISTER CHROMATID SEGREGATION | -2.19 | 0.059 | -2.36 | 0.005 | 0.85 | 1 |
| GO NEGATIVE REGULATION OF CHROMOSOME SEGREGATION | -2.01 | 0.548 | -2.35 | 0.007 | -0.84 | 1 |
| GO NEGATIVE REGULATION OF METAPHASE ANAPHASE TRANSITION OF CELL CYCLE | -2.13 | 0.155 | -2.46 | 0 | -0.98 | 1 |
| GO NUCLEAR TRANSCRIBED MRNA CATABOLIC PROCESS NONSENSE MEDIATED DECAY | 2.25 | 0.059 | 2.36 | 0.003 | 2.49 | 0.001 |
| GO PHAGOCYTOSIS RECOGNITION | -0.69 | 1 | 1.60 | 1 | -2.89 | 0 |
| GO POSITIVE REGULATION OF B CELL ACTIVATION | -0.51 | 1 | 1.50 | 1 | -2.35 | 0.001 |
| GO POSITIVE REGULATION OF LEUKOCYTE CELL CELL ADHESION | 0.91 | 1 | 2.32 | 0.003 | -1.06 | 1 |
| GO POSITIVE T CELL SELECTION | 1.14 | 1 | 2.45 | 0.001 | 0.92 | 1 |
| GO PROTEIN LOCALIZATION TO ENDOPLASMIC RETICULUM | 2.16 | 0.195 | 1.98 | 0.711 | 2.93 | 0 |
| GO PROTEIN TARGETING TO MEMBRANE | 1.83 | 0.998 | 1.71 | 1 | 2.50 | 0.001 |
| GO REGULATION OF B CELL ACTIVATION | 0.68 | 1 | 1.84 | 0.995 | -2.29 | 0.004 |
| GO REGULATION OF CHROMOSOME SEPARATION | -2.06 | 0.34 | -2.49 | 0 | -0.84 | 1 |
| GO REGULATION OF HUMORAL IMMUNE RESPONSE | -1.03 | 1 | 1.24 | 1 | -2.59 | 0 |
| GO REGULATION OF SISTER CHROMATID SEGREGATION | -1.94 | 0.878 | -2.43 | 0 | -0.80 | 1 |
| GO RIBOSOMAL SUBUNIT | 0.93 | 1 | 1.41 | 1 | 2.79 | 0 |
| GO RIBOSOME | -0.80 | 1 | 1.25 | 1 | 2.64 | 0 |
| GO SMALL RIBOSOMAL SUBUNIT | -0.91 | 1 | 1.43 | 1 | 2.40 | 0.007 |
| GO STRUCTURAL CONSTITUENT OF RIBOSOME | 1.07 | 1 | 1.49 | 1 | 2.91 | 0 |
| GO TRANSLATIONAL INITIATION | 1.93 | 0.951 | 2.05 | 0.401 | 2.47 | 0.001 |
| GO TRANSLATIONAL TERMINATION | -1.98 | 0.68 | -2.51 | 0 | 1.83 | 0.997 |
| GO T CELL ACTIVATION INVOLVED IN IMMUNE RESPONSE | 1.27 | 1 | 2.46 | 0 | -1.22 | 1 |
| GO T CELL RECEPTOR COMPLEX | -0.60 | 1 | 2.35 | 0.003 | 0.65 | 1 |
| GO T CELL SELECTION | 1.21 | 1 | 2.58 | 0 | 1.06 | 1 |
| GO UNFOLDED PROTEIN BINDING | -1.08 | 1 | -2.34 | 0.007 | 1.36 | 1 |
| KEGG LEISHMANIA INFECTION | 1.70 | 1 | 2.44 | 0.001 | 1.01 | 1 |
| KEGG RIBOSOME | 2.53 | 0.001 | 2.68 | 0 | 3.10 | 0 |
| PID IL12 2PATHWAY | 0.99 | 1 | 2.35 | 0.003 | -1.04 | 1 |
| PID PLK1 PATHWAY | -2.61 | 0 | -2.38 | 0.003 | -1.27 | 1 |
| PID TCR PATHWAY | 0.81 | 1 | 2.37 | 0.003 | -0.94 | 1 |
| REACTOME ACTIVATION OF THE MRNA UPON BINDING OF THE CAP BINDING COMPLEX AND EIFS AND SUBSEQUENT BINDING TO | 2.26 | 0.056 | 2.23 | 0.022 | 2.75 | 0 |
| REACTOME ANTIGEN ACTIVATES B CELL RECEPTOR BCR LEADING TO GENERATION OF SECOND MESSENGERS | 0.80 | 1 | 1.91 | 0.947 | -3.01 | 0 |
| REACTOME BINDING AND UPTAKE OF LIGANDS BY SCAVENGER RECEPTORS | -0.75 | 1 | 1.58 | 1 | -2.74 | 0 |
| REACTOME CD22 MEDIATED BCR REGULATION | -1.40 | 1 | 2.18 | 0.073 | -3.25 | 0 |
| REACTOME CELL CYCLE CHECKPOINTS | -2.17 | 0.073 | -2.54 | 0 | -1.00 | 1 |
| REACTOME COMPLEMENT CASCADE | -1.04 | 1 | 0.94 | 1 | -2.78 | 0 |
| REACTOME CREATION OF C4 AND C2 ACTIVATORS | -0.99 | 1 | 1.52 | 1 | -3.07 | 0 |
| REACTOME CYCLIN A B1 B2 ASSOCIATED EVENTS DURING G2 M TRANSITION | -2.34 | 0.004 | -2.45 | 0 | -0.97 | 1 |
| REACTOME EUKARYOTIC TRANSLATION ELONGATION | 2.67 | 0 | 2.80 | 0 | 3.19 | 0 |
| REACTOME EUKARYOTIC TRANSLATION INITIATION | 2.56 | 0.001 | 2.60 | 0 | 3.23 | 0 |
| REACTOME FCERI MEDIATED CA 2 MOBILIZATION | 1.18 | 1 | 2.11 | 0.172 | -2.91 | 0 |
| REACTOME FCERI MEDIATED MAPK ACTIVATION | 0.89 | 1 | 1.78 | 1 | -2.88 | 0 |
| REACTOME FCGAMMA RECEPTOR FCGR DEPENDENT PHAGOCYTOSIS | 1.40 | 1 | 1.61 | 1 | -2.41 | 0 |
| REACTOME FCGR3A MEDIATED IL10 SYNTHESIS | 0.76 | 1 | 1.93 | 0.891 | -2.86 | 0 |
| REACTOME FCGR ACTIVATION | 0.97 | 1 | 2.01 | 0.558 | -3.27 | 0 |
| REACTOME GENERATION OF SECOND MESSENGER MOLECULES | 1.34 | 1 | 2.57 | 0 | 1.17 | 1 |
| REACTOME IMMUNOREGULATORY INTERACTIONS BETWEEN A LYMPHOID AND A NON LYMPHOID CELL | 0.52 | 1 | 2.39 | 0.003 | -2.21 | 0.02 |
| REACTOME INFLUENZA INFECTION | 1.42 | 1 | 1.79 | 1 | 2.70 | 0 |
| REACTOME INITIAL TRIGGERING OF COMPLEMENT | -0.86 | 1 | 1.42 | 1 | -3.05 | 0 |
| REACTOME INTERLEUKIN 10 SIGNALING | 0.72 | 1 | 1.92 | 0.929 | -2.45 | 0 |
| REACTOME MITOCHONDRIAL TRANSLATION | -2.06 | 0.339 | -2.48 | 0 | 1.80 | 1 |
| REACTOME MITOTIC METAPHASE AND ANAPHASE | -1.99 | 0.624 | -2.46 | 0 | 1.20 | 1 |
| REACTOME MITOTIC PROMETAPHASE | -1.89 | 0.969 | -2.34 | 0.007 | -0.83 | 1 |
| REACTOME MITOTIC SPINDLE CHECKPOINT | -2.21 | 0.042 | -2.69 | 0 | -0.71 | 1 |
| REACTOME NONSENSE MEDIATED DECAY NMD | 2.28 | 0.039 | 2.42 | 0.001 | 2.66 | 0 |
| REACTOME PARASITE INFECTION | 0.90 | 1 | 1.39 | 1 | -2.59 | 0 |
| REACTOME REGULATION OF EXPRESSION OF SLITS AND ROBOS | 1.49 | 1 | 1.83 | 0.999 | 2.52 | 0.001 |
| REACTOME RESOLUTION OF D LOOP STRUCTURES THROUGH SYNTHESIS DEPENDENT STRAND ANNEALING SDSA | -2.15 | 0.101 | -2.43 | 0 | -1.38 | 1 |
| REACTOME RESOLUTION OF SISTER CHROMATID COHESION | -2.31 | 0.005 | -2.71 | 0 | -0.72 | 1 |
| REACTOME RESPONSE OF EIF2AK4 GCN2 TO AMINO ACID DEFICIENCY | 2.21 | 0.111 | 2.55 | 0 | 2.73 | 0 |
| REACTOME ROLE OF LAT2 NTAL LAB ON CALCIUM MOBILIZATION | 1.12 | 1 | 1.94 | 0.852 | -3.13 | 0 |
| REACTOME ROLE OF PHOSPHOLIPIDS IN PHAGOCYTOSIS | 1.41 | 1 | 2.25 | 0.016 | -3.04 | 0 |
| REACTOME RRNA PROCESSING | 0.91 | 1 | 1.32 | 1 | 2.69 | 0 |
| REACTOME SCAVENGING OF HEME FROM PLASMA | 1.37 | 1 | 2.20 | 0.038 | -3.27 | 0 |
| REACTOME SELENOAMINO ACID METABOLISM | 1.87 | 0.993 | 2.31 | 0.008 | 2.95 | 0 |
| REACTOME SEPARATION OF SISTER CHROMATIDS | -2.05 | 0.363 | -2.54 | 0 | 1.10 | 1 |
| REACTOME SIGNALING BY ROBO RECEPTORS | 1.61 | 1 | 1.90 | 0.961 | 2.40 | 0.007 |
| REACTOME SRP DEPENDENT COTRANSLATIONAL PROTEIN TARGETING TO MEMBRANE | 2.30 | 0.027 | 2.24 | 0.021 | 3.14 | 0 |
| REACTOME TRANSLATION | -1.14 | 1 | -1.34 | 1 | 2.93 | 0 |
| WP CYTOPLASMIC RIBOSOMAL PROTEINS | 2.71 | 0 | 2.77 | 0 | 3.17 | 0 |
| WP MICROGLIA PATHOGEN PHAGOCYTOSIS PATHWAY | 1.79 | 0.999 | 2.61 | 0 | 1.63 | 1 |
| WP TYROBP CAUSAL NETWORK | 2.30 | 0.024 | 2.83 | 0 | 1.16 | 1 |

***Sup. Table 1: Significantly enriched pathways for R1, R2* and the r1-r2* relaxivity.*** *For each of the 101 gene sets, we show the normalized enrichments score (NES) for each of the three qMRI parameters, along with the FWER-corrected p-value. This table was used for the clustering shown in Figure 5b.*

### Supplementary Figure 17

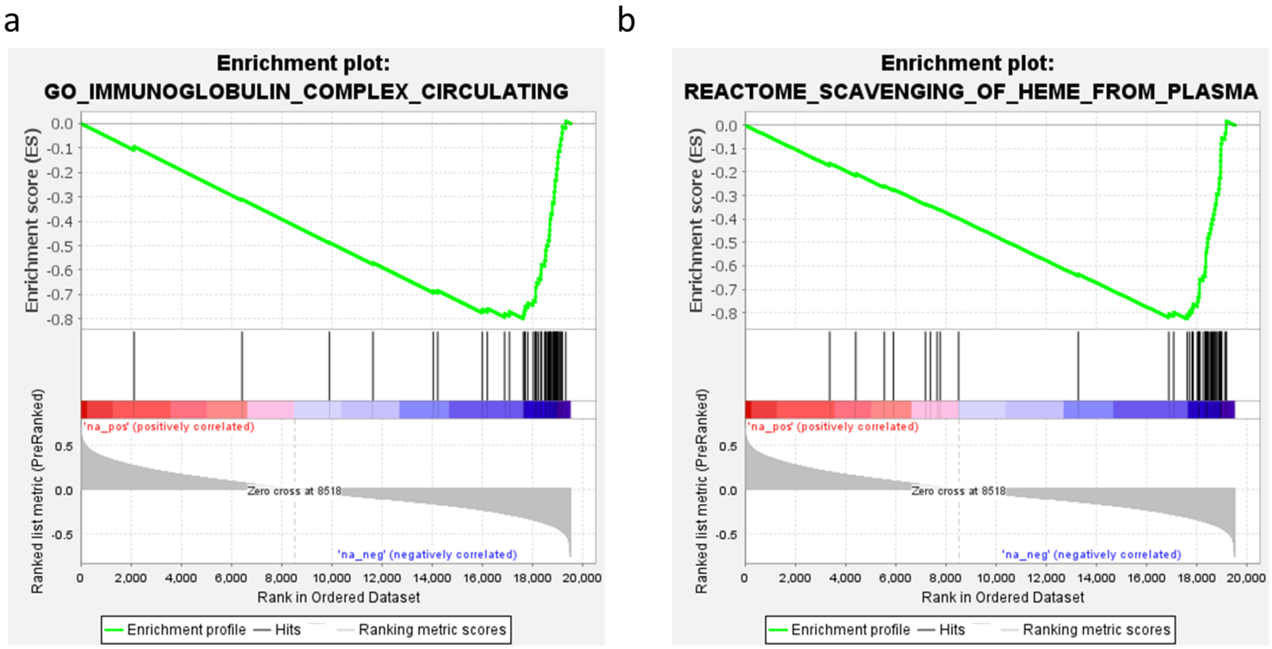

***Sup. Figure 17: Enrichment plots of the two most enriched pathways for the r1-r2* relaxivity.*** *Gene enrichment plots of the two most enriched pathways for the r1-r2* relaxivity "Immunoglobulin complex"* ***(a)*** *and "scavenging of heme from plasma"* ***(b)****. The top portion of each panel shows the running enrichment score for the gene set as the analysis goes over the ranked list of genes. The list is based on the genes' correlation with the r1-r2* relaxivity. The middle portion of each panel shows where the members of the gene set appear in the ranked list of genes. The bottom portion of each panel shows the r value of the correlation between genes and the r1-r2* relaxivity. The two gene sets preferentially fall toward the negative end of the correlation spectrum, indicating their significant association with the r1-r2* relaxivity.*

#### Supplementary Figure 18

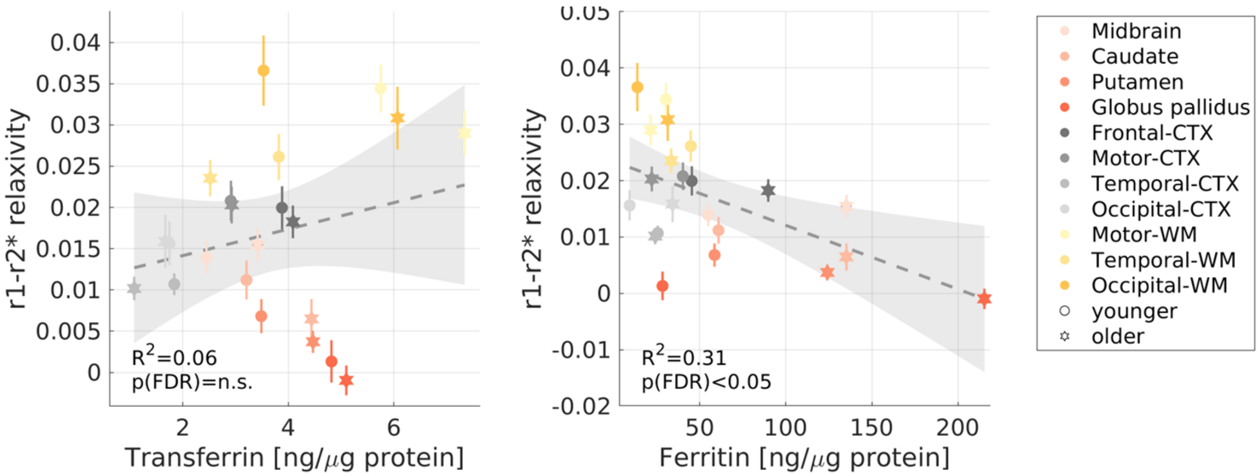

***Sup. Figure 18: The correlations of the r1-r2* relaxivity with the transferrin and ferritin concentrations.*** *The transferrin and ferritin concentrations (postmortem, from the literature[5,7,9]) in different brain regions of younger (aged 27-64 years, N>=7) and older (aged 65-88 years, N>=8) subjects vs. the r1-r2* relaxivity measured in vivo across younger (aged 23-63 years, N =26) and older (aged 65-77 years, N=13) subjects (different marker shapes) in 10 brain regions (different colors).*

### Supplementary Section 4: Exploring the biophysical sources of the r1-r2* relaxivity.

#### Supplementary Section 4.1: The theoretical basis for the r1-r2* relaxivity of brain tissue.

Brain tissue contains a complex milieu of iron compounds with variable iron binding capacities and aggregation states and includes myelin. Here we will expand the biophysical model of the r1-r2* relaxivity ("*In vivo* iron relaxivity model" in Methods) for the case of a heterogenous iron environment and in the presence of myelin.

Assuming the iron environment of a myelinated tissue contains N different iron compounds, and under the assumption that water can freely diffuse, the MR relaxation rates can be expressed as[87,88]:

$$\text{S1)} \qquad R_1 = \sum_N r_{(1,i)}[i] + r_{(1,M)}[M] + C_1$$

$$\text{S2)} \qquad R_2^* = \sum_N r_{(2,i)}[i] + r_{(2,M)}[M] + C_2$$

Where $[i]$ is the concentration of the i'th iron compound, and $[M]$ is the myelin concentration. $r_{(1,i)}$ and $r_{(2,i)}$ are the R1- and R2*- relaxivities of the i'th iron compound. $r_{(1,M)}$ and $r_{(2,M)}$ are the R1- and R2*-relaxivities of myelin. $\mathrm{C}_1$ and $\mathrm{C}_2$ are constants which capture other non-iron and non-myelin contributions.

The r1-r2* relaxivity measurement is defined as the linear dependency of R1 on R2* within an ROI in the brain (or across *in vitro* samples). This is equivalent to the total change in R1 relative to the total change in R2* ($\frac{\Delta R_1}{\Delta R_2^*}$):

$$\text{S3)} \qquad \frac{\Delta R_1}{\Delta R_2^*} = \frac{\sum_N r_{(1,i)}[\Delta i] + r_{(1,M)}[\Delta M]}{\sum_N r_{(2,i)}[\Delta i] + r_{(2,M)}[\Delta M]}$$

Where $[\Delta i]$ and $[\Delta M]$ are the changes in the iron compounds and myelin concentrations within an ROI respectively.

#### Supplementary Section 4.2: The r1-r2* relaxivity of ferritin and transferrin mixtures *in vitro*.

First, we tested the theoretical formulation presented in eq. S1-S3 in an artificial environment of multiple iron compounds. For this aim, we constructed phantom experiments containing both ferritin and transferrin in a liposomal environment. In this synthetic toy model, the transferrin-ferritin fraction ($f = \frac{[Tf]}{[Ft]+[Tf]}$) represents an example for a feature of the molecular iron environment which should affect the r1-r2* relaxivity. To test this assumption, the biophysical model shown in Supp. Section 4.1 can be further developed under the conditions of these phantom experiments:

When ferritin, transferrin and myelin are present, eq. S*1*-S2 can be expressed as:

$$\text{S4)} \qquad R_1 = r_{(1,Ft)}[Ft] + r_{(1,Tf)}[Tf] + r_{(1,M)}[M]$$

$$\text{S5)} \qquad R_2^* = r_{(2,Ft)}[Ft] + r_{(2,Tf)}[Tf] + r_{(2,M)}[M]$$

Where $[Ft]$, $[Tf]$ and $[M]$ are the ferritin, transferrin and myelin concentrations respectively. $r_{(1,Ft)}$, $r_{(1,Tf)}$ and $r_{(1,M)}$ are the R1-relaxivities of ferritin, transferrin and myelin respectively. $r_{(2,Ft)}$, $r_{(2,Tf)}$ and $r_{(2,M)}$ are the R2*-relaxivities of ferritin, transferrin and myelin respectively.

According to eq. S3, the r1-r2* relaxivity measurement is equivalent to the total change in R1 relative to the total change in R2* ($\frac{\Delta R_1}{\Delta R_2^*}$):

$$\text{S6)} \qquad \frac{\Delta R_1}{\Delta R_2^*} = \frac{r_{(1,Ft)}[\Delta Ft] + r_{(1,Tf)}[\Delta Tf] + r_{(1,M)}[\Delta M]}{r_{(2,Ft)}[\Delta Ft] + r_{(2,Tf)}[\Delta Tf] + r_{(2,M)}[\Delta M]}$$

Where $[\Delta Ft]$, $[\Delta Tf]$, and $[\Delta M]$ are the changes in the ferritin, transferrin and myelin concentrations across *in vitro* samples, respectively.

In the ferritin-transferrin mixtures experiments we tested four different transferrin-ferritin fractions ($f = \frac{[Tf]}{[Ft]+[Tf]}$). For each fraction, we prepared samples with varying ferritin and transferrin concentrations, while keeping the fixed fraction between them (**Sup. Figure 19**). This allowed us to fit the linear relationship between R1 and R2* (the r1-r2* relaxivity) for each transferrin-ferritin fraction (**Sup. Figure 20**).

Assuming the transferrin-ferritin fraction ($f$) remains fixed across the *in vitro* samples over which the r1-r2* relaxivity is calculated:

$$f = \frac{Tf_1}{Tf_1 + Ft_1} = \frac{Tf_0}{Tf_0 + Ft_0}$$

(defining $[\Delta Tf] = Tf_1 - Tf_0$ and $[\Delta Ft] = Ft_1 - Ft_0$).

Under this condition:

$$\text{S7)} \qquad f = \frac{[\Delta Tf]}{[\Delta Ft]+[\Delta Tf]} = \frac{[Tf]}{[Ft]+[Tf]}$$

And therefore eq. S6 can be expressed as:

S8) $$\frac{\Delta R_1}{\Delta R_2^*} = \frac{(f*r_{(1,Tf)}+(1-f)r_{(1,Ft)})[\Delta iron]+r_{(1,M)}[\Delta M]}{(f*r_{(2,Tf)}+(1-f)r_{(2,Ft)})[\Delta iron]+r_{(2,M)}[\Delta M]}$$

Where $[\Delta Ft] + [\Delta Tf] = [\Delta iron]$.

In the ferritin-transferrin mixtures experiments, the liposomal fraction, which mimics the effect of myelin, was fixed at 17.5%. Therefore, in this case $[\Delta M] = 0$ and eq. S8 reduces to:

S9) $$\frac{\Delta R_1}{\Delta R_2^*} = \frac{(f*r_{(1,Tf)}+(1-f)r_{(1,Ft)})[\Delta iron]}{(f*r_{(2,Tf)}+(1-f)r_{(2,Ft)})[\Delta iron]} = \frac{(f*r_{(1,Tf)}+(1-f)r_{(1,Ft)})}{(f*r_{(2,Tf)}+(1-f)r_{(2,Ft)})}$$

Using the ferritin and transferrin relaxivities ($r_{(1/2,Tf/Ft)}$) measured for each liposomal iron compound individually (Figure 1a-c), we could test the prediction of this model. Notably, the theoretical r1-r2* relaxivities calculated with eq. $S9$ were in agreement with the experimental r1-r2* relaxivities (**Sup. Figure 21**), validating the presented biophysical framework *in vitro*. Moreover, the differences in the r1-r2* relaxivities measured for different transferrin-ferritin fractions were above the detection limit of this MRI measurement as estimated in a scan-rescan experiment (MAE=3.9*$10^{-4}$, **Sup. Figure 8**). Therefore, this toy example demonstrates that the heterogeneity in the iron environment can produce measurable changes in the r1-r2* relaxivity, which are in agreement with the biophysical modeling of this measurement.

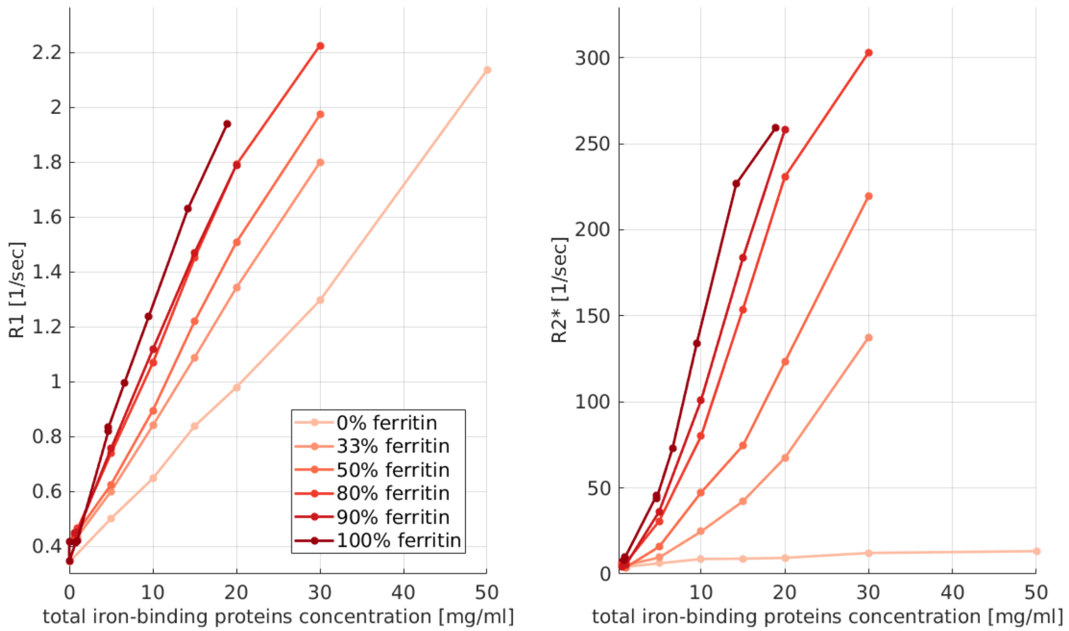

***Sup. Figure 19: The dependency of R1 (left) and R2* (right) on the total iron-binding proteins concentration for four transferrin-ferritin mixtures.*** *Each mixture has a different transferrin-ferritin fraction (different colors, legend shows the percentage of ferritin in the mixture). Data points are different transferrin-ferritin samples, line connect between samples with the same transferrin-ferritin fraction and varying total iron-binding protein concentrations.*

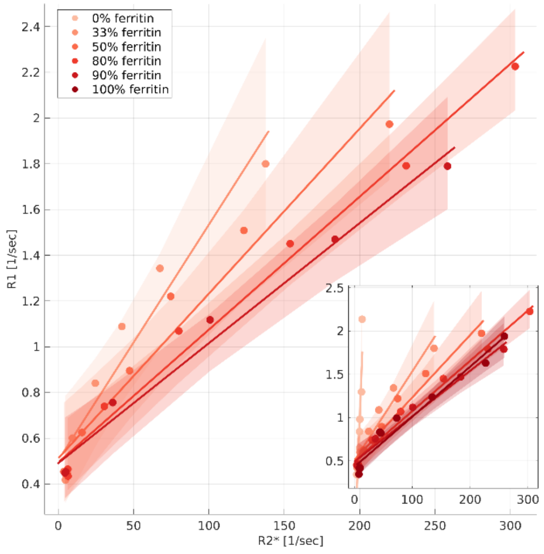

***Sup. Figure 20: The dependency of R1 on R2* for four transferrin-ferritin mixtures.*** *Each mixture has a different transferrin-ferritin fraction (different colors, legend shows the percentage of ferritin in the mixture). Data points represent samples with varying total iron-binding proteins concentrations. The linear relationships of R1 and R2* are marked by lines. The slopes of these lines are the r1-r2* relaxivities. Shaded areas represent the 95% confidence bounds. Inset shows the pure ferritin (100% ferritin) and pure transferrin (0% ferritin) samples as well.*

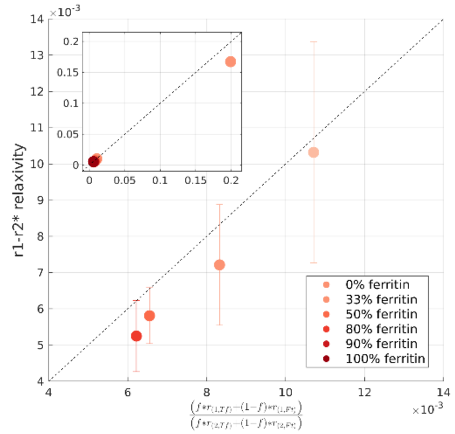

***Sup. Figure 21: The theoretical r1-r2* relaxivities calculated with eq. S9 are in agreement with the experimental r1-r2* relaxivities.*** *The y-axis shows the r1-r2* relaxivity calculated for four transferrin-ferritin mixtures with different transferrin-ferritin fractions (different colors, Sup. Figure 20). Errorbars show the 95% confidence bounds. The x-axis shows the prediction for the r1-r2* relaxivity based on eq. S9. The ferritin and transferrin relaxivities in the equation were plugged in based on our experimental results for liposomal ferritin and liposomal transferrin samples (*Figure 1a-c*). f represents the transferrin-ferritin fractions and varies between data points. Dashed line is the identity line. Inset shows the data points for pure ferritin (100% ferritin) and pure transferrin (0% ferritin) samples as well. Note that for these pure samples the prediction is already presented in figure 1e.*

#### Supplementary Section 4.3: Numerical simulations of the r1-r2* relaxivity.

The phantom experiments of ferritin and transferrin mixtures allowed us to establish a theoretical framework for the r1-r2* relaxivity in an *in vitro* environment where only the iron concentration changes, and the liposomal fraction mimicking the myelin is fixed ($[\Delta M] = 0$). In the brain, we estimate the r1-r2* relaxivity across all voxels of an anatomically-defined ROI. Within brain tissue ROIs, both the iron and the myelin concentration may vary[45].

Importantly, rearranging eq. S8 we find that the strength of the myelin effect on the r1-r2* relaxivity depends on how variable is the myelin content within an ROI relative to how variable is the iron content ($\frac{[\Delta M]}{[\Delta iron]}$):

$$S10) \qquad \frac{\Delta R_1}{\Delta R_2^*} = \frac{\left(f * r_{(1,Tf)} + (1-f) r_{(1,Ft)}\right) + r_{(1,M)} \frac{[\Delta M]}{[\Delta iron]}}{\left(f * r_{(2,Tf)} + (1-f) r_{(2,Ft)}\right) + r_{(2,M)} \frac{[\Delta M]}{[\Delta iron]}}$$

In order to evaluate how the r1-r2* relaxivity is modulated by the molecular iron environment and by the myelin and iron variabilities within an ROI in the brain, we performed a set of numerical simulations. In these simulations, similar to the *in vitro* mixtures experiments, the transferrin-ferritin fraction ($f =$

$$\Delta(iron)R_2^* = \Delta(total)R_2^* - r_{(2,M)}[\Delta M]$$

We found that the total change in R2* within ROIs in the human brain is on average $\Delta(total)R_2^*$=9.0 1/sec, from which about 61% ($\Delta(iron)R_2^*$=5.6 1/sec) could be related to changes in iron concentration. Therefore, the simulated variably in the ferritin and transferrin concentrations were set to satisfy this requirement.

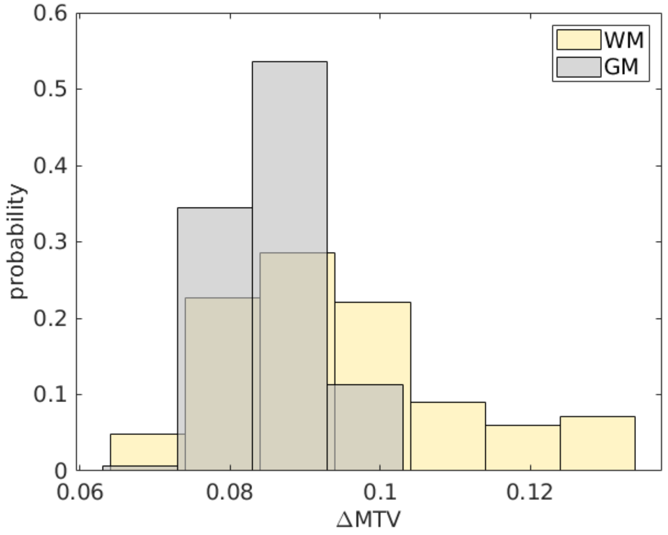

***Sup. Figure 22: Change in MTV values ($[\Delta MTV]$) within white-matter (WM) or gray-matter (GM) regions.*** *ΔMTV values for gray matter (GM) and white matter (WM) are presented across 16 ROIs in the brains of 21 young subjects. For each ROI, we extracted the MTV values from all voxels and pooled them into 36 bins spaced equally between 0.05 and 0.40 [fraction]. We removed any bins in which the number of voxels was smaller than 4% of the total voxel count in the ROI. This was done so that the calculation will not be heavily affected by outlier voxels with extreme values. The median MTV of each bin was computed, and the difference between the highest and lowest binned MTV values was set as ΔMTV ([fraction]) in the ROI.*

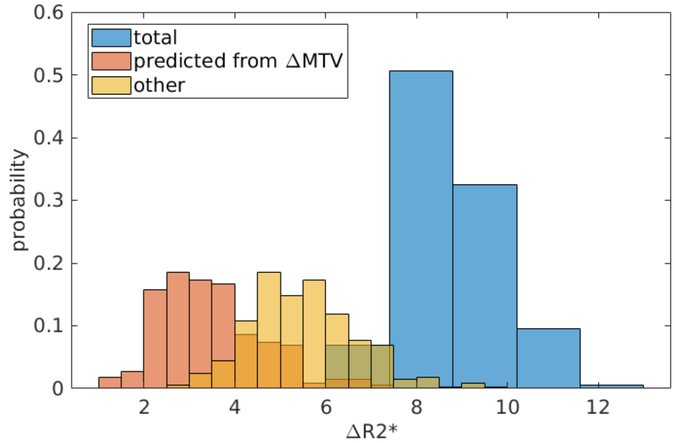

***Sup. Figure 23: Myelin- and iron- related changes in R2* within brain regions. Total change in R2* values*** *($[\Delta R2 *]$)* ***within brain regions (blue histogram)*** *is presented across 16 ROIs in the brains of 21 young subjects. For each ROI, we extracted the R2* values from all voxels and pooled them into 36 bins spaced equally between 0 and 50. We removed any bins in which the number of voxels was smaller than 4% of the total voxel count in the ROI. This was done so that the calculation will not be heavily affected by outlier voxels with extreme values. The median R2* of each bin was computed, and the difference between the highest and lowest binned R2* values was set as ΔR2* in the ROI (in [1/sec]).* ***R2* changes related to myelin (orange histogram)*** *were estimated based on MTV; the change in R2* predicted from the change in MTV (ΔMTV) within each ROI was calculated as the linear dependency of R2* on MTV in the ROI multiplied by ΔMTV in the ROI ($r_{(2,M)}[\Delta M]$).* ***R2* changes related to iron (yellow histogram)*** *were estimated as the change in R2* not explained by the change in MTV.*

| *Parameter* | *Value* | *Estimation method* |
|---|---|---|
| *Transferrin-ferritin fraction* ($f$) | 0.1 or 0.2 | Based on literature values[5,7,9]. |
| *R1-ferritin relaxivity* ($r_{(1,Ft)}$) | 0.067 [($sec^{-1}$)/(mg/wet ml)] | *In vitro* linear dependency of R1 on ferritin concentration |
| *R2*-ferritin relaxivity* ($r_{(2,Ft)}$) | 11.2 [($sec^{-1}$)/(mg/wet ml)] | *In vitro* linear dependency of R2* on ferritin concentration |
| *R1-transferrin relaxivity* ($r_{(1,Tf)}$) | 0.026 [($sec^{-1}$)/(mg/wet ml)] | *In vitro* linear dependency of R1 on transferrin concentration |
| *R2*-transferrin relaxivity* ($r_{(2,Tf)}$) | 0.13 [($sec^{-1}$)/(mg/wet ml)] | *In vitro* linear dependency of R2* on transferrin concentration |
| *Transferrin concentration* ($[Tf]$) | 0.025±0.025 [mg/wet ml] | Median is based on literature values[5,7,9]. Range across voxels was set so that the total change in R2* will mimic the physiological change of 6-12 [1/sec] (**Sup. Figure 23**) |
| *Ferritin concentration* ($[Ft]$) | $\left(\frac{1}{f}-1\right)[Tf]$ | Set to satisfy the requirement for fixed transferrin-ferritin fraction ($f$) across all voxels. |
| *Myelin concentration in WM* ($[M]_{WM}$) | 0.29±0.047 [fraction] | Brain *in vivo* MTV values |
| *Myelin concentration in GM* ($[M]_{GM}$) | 0.19±0.043 [fraction] | Brain *in vivo* MTV values |
| *R2*-myelin relaxivity* ($r_{(2,M)}$) | 38.8 [($sec^{-1}$)/fraction] | Brain *in vivo* linear dependency of R2* on MTV |
| *R1-myelin relaxivity* ($r_{(1,M)}$) | 2.6 [($sec^{-1}$)/fraction] | Brain *in vivo* linear dependency of R1 on MTV |

***Sup. Table 2: simulation parameters.***

An example result of the numerical simulation is presented in **Sup. Figure 24A**. It is evident that both the changes in myelin concentration across voxels of the simulated ROI (represented by different colors) and the changes in ferritin and transferrin concentrations across voxels (represented by the symbols size) affect the measured r1-r2* relaxivity. In our analysis pipeline we first bin the R1 and R2* values within the ROI and next calculate r1-r2* relaxivity over the binned values. Therefore, the variability in R1 for a given R2* bin is collapsed to an average R1 value (black data points). We assume that this approach eliminates some of the variability related to myelin. We hypothesized that the r1-r2* relaxivity is sensitive to the iron environment. Indeed, in our simulations we find that by setting different physiological transferrin-ferritin fractions and leaving the myelin parameters constant, the r1-r2* relaxivity changes considerably (**Sup. Figure 24**).

In addition, we hypothesized that the r1-r2* relaxivity less sensitive to the iron concentration and is more sensitive to the iron environment. To test this, we run two numerical simulations with the same transferrin-ferritin fraction but with different transferrin and ferritin concentrations. As expected, R1 and R2* values changed with increased ferritin and transferrin concentrations but the r1-r2* relaxivity did not change (**Sup. Figure 25**). This indicates that the r1-r2* relaxivity measurement is less

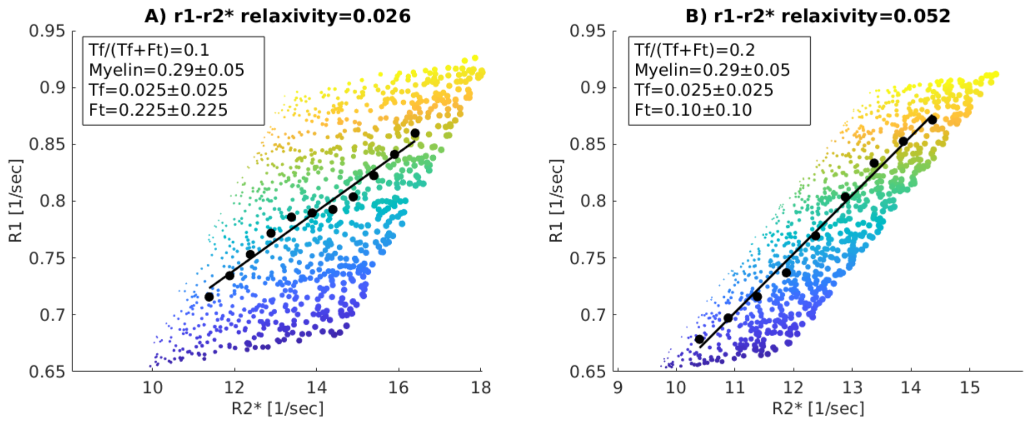

***Sup. Figure 24: The r1-r2* relaxivity in two simulated ROIs with different transferrin-ferritin fractions (Tf/(Tf+Ft))****; (A) the transferrin-ferritin fraction is 0.1; (B) the transferrin-ferritin fraction is 0.2. Each figure shows the dependency of R1 on R2* for 1,000 representative simulated voxels. The colors of the data points indicate the variability in myelin concentration across voxels, and their sizes indicate the variability in iron compounds concentration across voxels (the simulated concentrations are shown in the text box, myelin is in units of [fraction] as MTV, transferrin and ferritin are in units of [mg/ml]). As in our in vivo pipeline, R2* and R1 values were binned (black data points represent the bins' median), and a linear fit was calculated (black line). The slopes of the linear fit (shown in the title) represent the dependency of R1 on R2* (r1-r2* relaxivity) and vary with the transferrin-ferritin fraction.*

sensitive to absolute changes in the iron concentration and is sensitive to the interplay between iron compounds, reflected in the simulations by the transferrin-ferritin fraction. In the brain, changes in the myelin content between GM and WM are known to substantially affect the measurements of R1 and R2*[4,19,25,29,40–44]. To test the potential contribution of the myelin to the r1-r2* relaxivity, we changed the myelin concentration in our simulation while keeping the rest of the parameters fixed. Setting the myelin concentration to that typical for GM or WM (as estimated *in vivo* by MTV) led to considerable changes in R1 and R2*, but did not produce any change in the r1-r2* relaxivity (**Sup. Figure 26**).

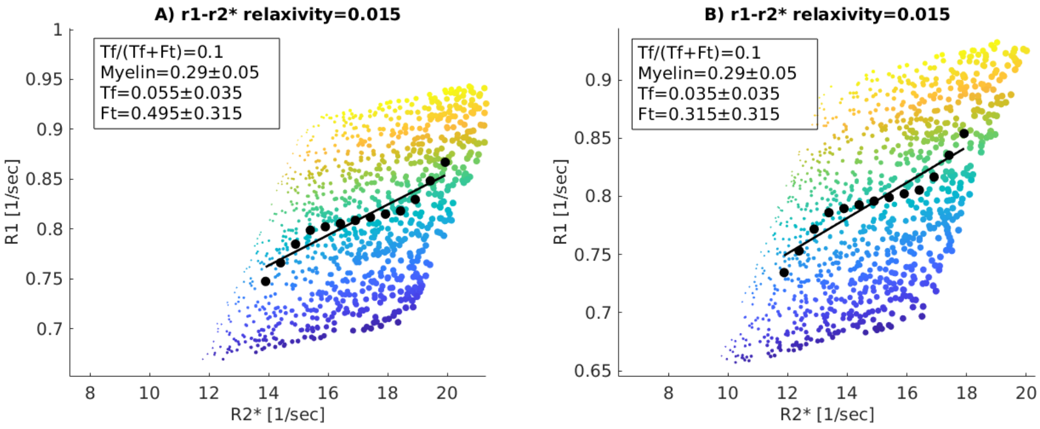

***Sup. Figure 25: The r1-r2* relaxivity in two simulated ROIs with different transferrin and ferritin concentrations and a similar transferrin-ferritin fraction (Tf/(Tf+Ft));*** *(A) a higher transferrin and ferritin*

*concentrations and a transferrin-ferritin fraction of 0.1; (B) a lower transferrin and ferritin concentrations and a transferrin-ferritin fraction of 0.1. Each figure shows the dependency of R1 on R2* for 1,000 representative simulated voxels. The colors of the data points indicate the variability in myelin concentration across voxels, and their sizes indicate the variability in iron compounds concentration across voxels (the simulated concentrations are shown in the text box, myelin is in units of [fraction] as MTV, transferrin and ferritin are in units of [mg/ml]). As in our in vivo pipeline, R2* and R1 values were binned (black data points represent the bins' median), and a linear fit was calculated (black line). The slopes of the linear fit (shown in the title) represent the dependency of R1 on R2* (r1-r2* relaxivity) and do vary with the change in transferrin and ferritin concentrations.*

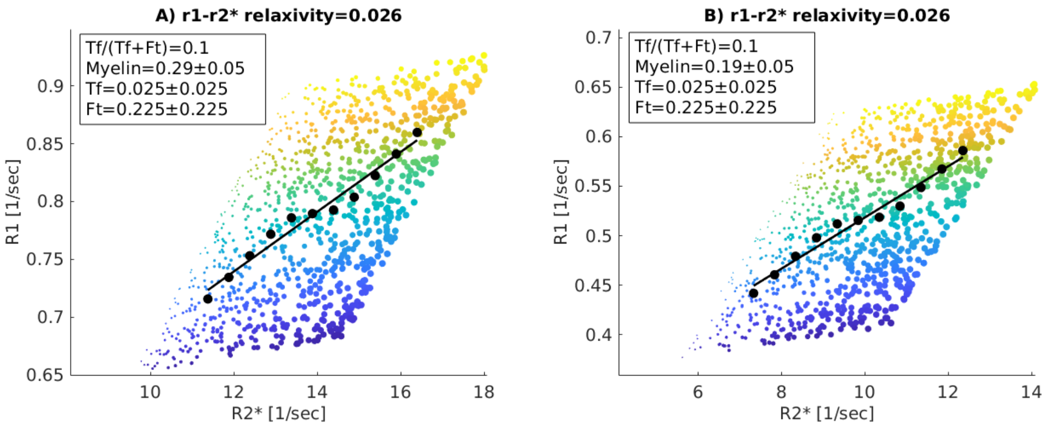

***Sup. Figure 26: The r1-r2* relaxivity in two simulated ROIs with different myelin concentrations and similar transferrin-ferritin fractions (Tf/(Tf+Ft))****; (A) a higher myelin concentration and transferrin-ferritin fraction of 0.1; (B) a lower myelin concentration and transferrin-ferritin fraction of 0.1. Each figure shows the dependency of R1 on R2* for 1,000 representative simulated voxels. The colors of the data points indicate the variability in myelin concentration across voxels, and their sizes indicate the variability in iron compounds concentration across voxels (the simulated concentrations are shown in the text box, myelin is in units of [fraction] as MTV, transferrin and ferritin are in units of [mg/ml]). As in our in vivo pipeline, R2* and R1 values were binned (black data points represent the bins' median), and a linear fit was calculated (black line). The slopes of the linear fit (shown in the title) represent the dependency of R1 on R2* (r1-r2* relaxivity) and does not vary with the change in myelin concentration.*

The theoretical formulation presented here indicates that it is not the myelin concentration, but the variability in myelin within an ROI ($[\Delta M]$), that is important for determining the r1-r2* relaxivity (eq. S10). However, estimating the variability in myelin within ROIs *in vivo* based on the myelin marker MTV, we find that it only explains ~30% of the variation in the *in vivo* r1-r2* relaxivity measurements across the brain (**Sup. Figure 27**). In the simulations, setting both the range and concentration of myelin to the ones typical for WM or GM (as estimated by MTV, **Sup. Figure 22**), slightly changed the r1-r2* relaxivity (0.002, **Sup. Figure 28**). Importantly, changing the transferrin-ferritin fraction between the physiological values of 0.1-0.2 led to a change of 0.026 in the r1-r2* relaxivity (**Sup. Figure 24**). Therefore, the simulated changes related to the molecular iron environment were one order of magnitude bigger (**Sup. Figure 24**).

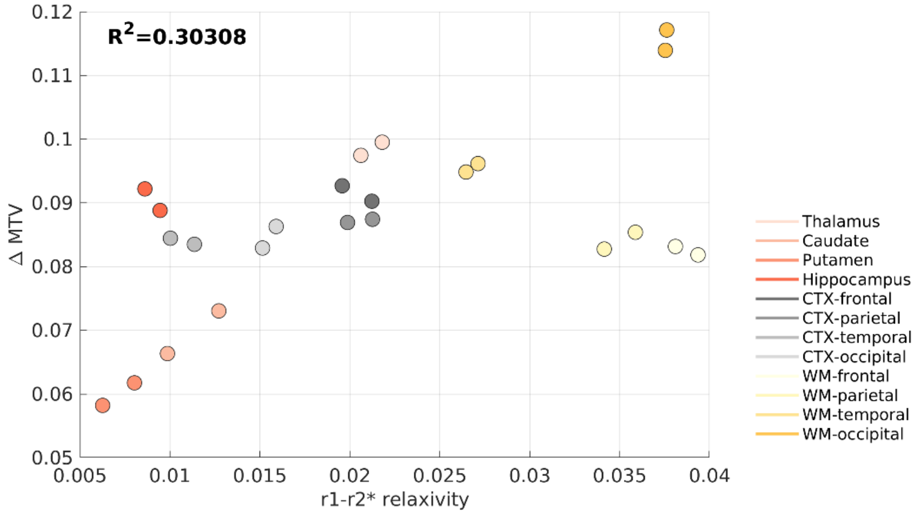

***Sup. Figure 27: The in vivo estimates of the variability in myelin within ROIs vs. the in vivo r1-r2* relaxivity.*** *The average estimated myelin variability within ROIs, computed across the brains of 21 young subjects, vs. the average r1-r2* relaxivity for the same subjects. Brain regions are presented in different colors (each ROI has left and right hemisphere estimates). The variability in myelin within brain regions was estimated based on the variability in MTV across voxels of each ROI (ΔMTV, y-axis). For each ROI, we extracted the MTV values from all voxels and pooled them into 36 bins spaced equally between 0.05 and 0.40 [fraction]. We removed any bins in which the number of voxels was smaller than 4% of the total voxel count in the ROI. This was done so that the calculation will not be heavily affected by outlier voxels with extreme values. The median MTV of each bin was computed, and the difference between the highest and lowest binned MTV values was set as ΔMTV ([fraction]) in the ROI.*

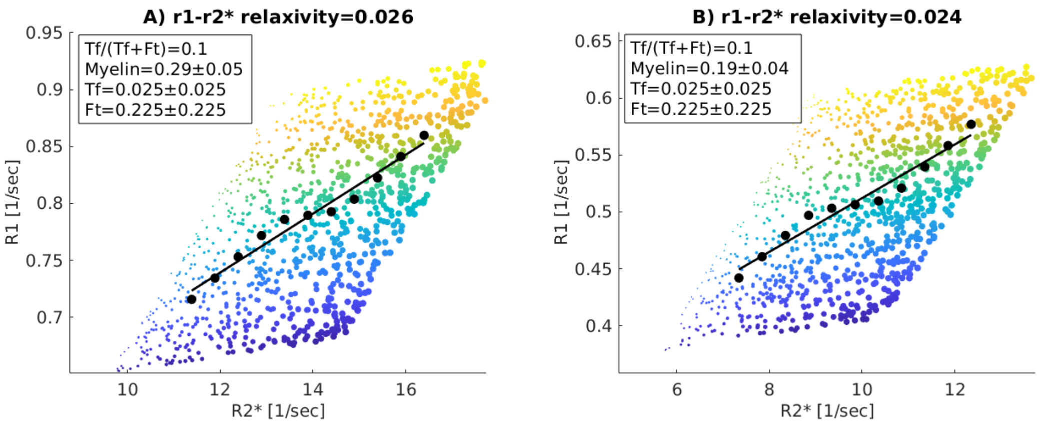

***Sup. Figure 28: The r1-r2* relaxivity in two simulated ROIs with different myelin concentrations, different ranges of myelin concentrations ($[\Delta M]$), and similar transferrin-ferritin fractions (Tf/(Tf+Ft));*** *(A) WM; a higher myelin concentration and a larger range of myelin variability, the transferrin-ferritin fraction is 0.1; (B) GM; a lower myelin concentration and a lower range of myelin variability, the transferrin-ferritin fraction is 0.1. Each figure shows the dependency of R1 on R2* for 1,000 representative simulated voxels. The colors of the data points indicate the variability in myelin concentration across voxels, and their sizes indicate the variability in iron compounds concentration*

*across voxels (the simulated concentrations are shown in the text box, myelin is in units of [fraction] as MTV, transferrin and ferritin are in units of [mg/ml]). As in our in vivo pipeline, R2* and R1 values were binned (black data points represent the bins' median), and a linear fit was calculated (black line). The slopes of the linear fit (shown in the title) represent the dependency of R1 on R2* (r1-r2* relaxivity) and vary slightly with the range of myelin variability ($[\Delta M]$).*

We further tested what are the myelin properties that would generate similar r1-r2* relaxivity effect as the effect observed when changing the transferrin-ferritin fraction (a change of 0.026 in the r1-r2* relaxivity, **Sup. Figure 24**). As changing the myelin concentration does not change the r1-r2* relaxivity (**Sup. Figure 26**), we changed the variability in myelin concentration within the ROI ($[\Delta M]$, **Sup. Figure 28**). We found that in order to generate a change of 0.026 in the r1-r2* relaxivity only through myelin-related changes (when the iron-related properties are fixed), the variability in MTV within the simulated ROI should be in the order of 0.187 [fraction] (**Sup. Figure 29**). Evaluating the *in vivo* variability in MTV within WM, GM and subcortical ROIs across 21 young subjects, the typical variability is ~ 0.09 [fraction], and the most extreme variability that was measured was 0.13 [fraction] (in the WM, **Sup. Figure 22**). Even this atypical value is still much lower than that required to generate a change of 0.026 in the r1-r2* relaxivity (0.187 [fraction] in MTV). Thus, while the variability in myelin within ROIs can affect the r1-r2* relaxivity, there are no physiological myelin properties that would fully explain the r1-r2* relaxivity effect measured in the brain.

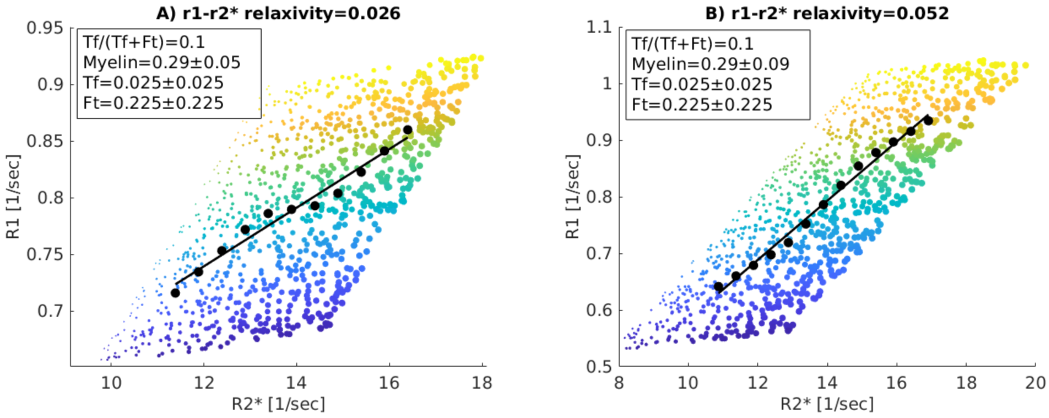

***Sup. Figure 29: The r1-r2* relaxivity in two simulated ROIs with different extreme ranges of myelin concentrations ($[\Delta M]$) and similar transferrin-ferritin fractions (Tf/(Tf+Ft));*** *(A) a physiological range of myelin variability, and a transferrin-ferritin fraction of 0.1; (B) the range of myelin variability is almost doubled, and the transferrin-ferritin fraction is 0.1. Each figure shows the dependency of R1 on R2* for 1,000 representative simulated voxels. The colors of the data points indicate the variability in myelin concentration across voxels, and their sizes indicate the variability in iron compounds concentration across voxels (the simulated concentrations are shown in the text box, myelin is in units of [fraction] as MTV, transferrin and ferritin are in units of [mg/ml]). As in our in vivo pipeline, R2* and R1 values were binned (black data points represent the bins' median), and a linear fit was calculated (black line). The*

*slopes of the linear fit (shown in the title) represent the dependency of R1 on R2* (r1-r2* relaxivity) and change considerably under this condition of extreme variability in myelin concentration ($[\Delta M]$).*

The iron and myelin contents of brain tissue are tightly related, as iron is required for the formation of myelin[4]. To test how this affects the r1-r2* relaxivity measurement, we simulated a case where iron and myelin are completely correlated. Importantly, even in this extreme case, different transferrin-ferritin fractions exhibited different r1-r2* relaxivity (**Sup. Figure 30** ).

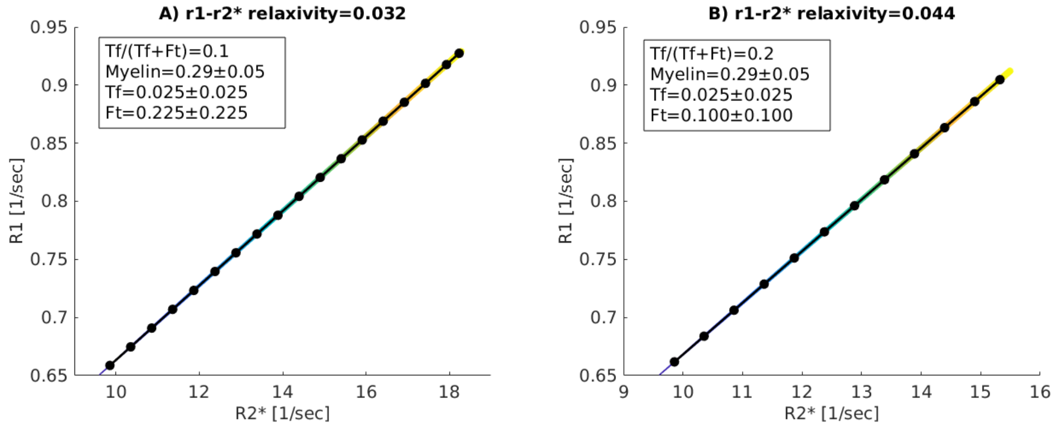

***Sup. Figure 30 : The r1-r2* relaxivity in two simulated ROIs with different transferrin-ferritin fractions (Tf/(Tf+Ft)) and a correlation between iron and myelin concentrations across voxels****; (A) The transferrin-ferritin fraction is 0.1; (B) the transferrin-ferritin fraction is 0.2. In both A and B iron and myelin are correlated. Each figure shows the dependency of R1 on R2* for 1,000 representative simulated voxels. The colors of the data points indicate the variability in myelin concentration across voxels, and their sizes indicate the variability in iron compounds concentration across voxels (the simulated concentrations are shown in the text box, myelin is in units of [fraction] as MTV, transferrin and ferritin are in units of [mg/ml]). As in our in vivo pipeline, R2* and R1 values were binned (black data points represent the bins' median), and a linear fit was calculated (black line). The slopes of the linear fit (shown in the title) represent the dependency of R1 on R2* (r1-r2* relaxivity) and change considerably with the transferrin-ferritin fraction even when iron and myelin are correlated.*

#### Supplementary Figure 31

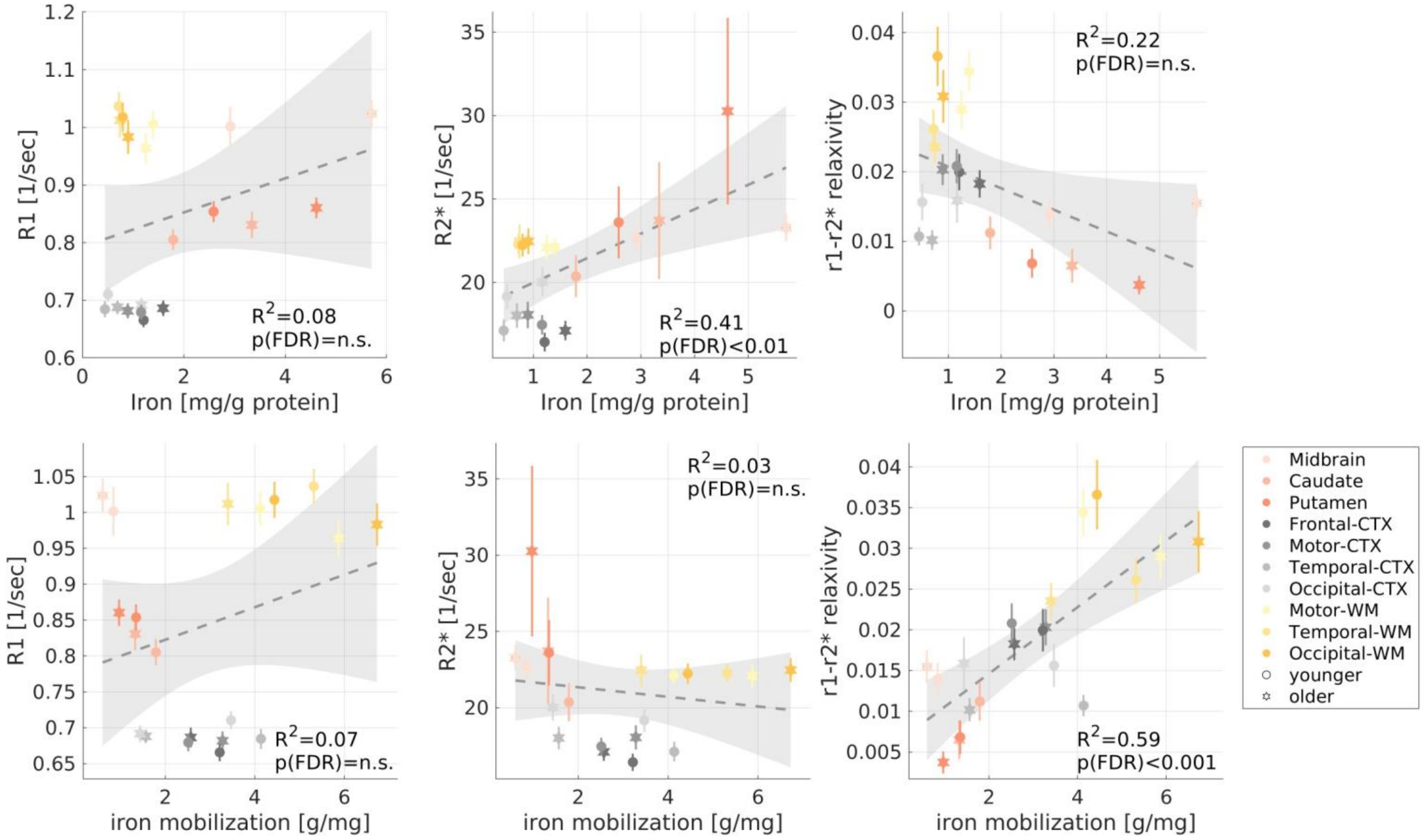

***Sup. Figure 31: The correlations of MRI parameters with the iron environment when excluding the pallidum. (a)*** *Replication of Figure* 3b-c *excluding the globus pallidus. The iron concentration and the iron mobilization capacity (transferrin/iron ratio) estimate postmortem (from the literature*[5,7,9]*) in different brain regions of younger (aged 27-64 years, N>=7) and older (aged 65-88 years, N>=8) subjects vs. R1, R2* and the r1-r2* relaxivity measured in vivo across younger (aged 23-63 years, N =26) and older (aged 65-77 years, N=13) subjects (different marker shapes) in 10 brain regions (different colors), excluding the pallidum. The correlation of the r1-r2* relaxivity with the iron concentration did not survive after excluding the pallidum. On the contrary, the correlations of R2* with the iron concentration and of the r1-r2* relaxivity with the iron mobilization remained significant.*

### Supplementary methods

##### R1-MTV dependency computation for phantoms.

We computed the linear dependency of R1 on MTV across samples with varying iron-binding protein concentrations and liposomal fractions[76]. This process was implemented in MATLAB. We extracted the MTV values from all voxels and pooled them into 12 bins spaced equally between 0.05 and 0.40. This was done so that the linear fit would not be heavily affected by the density of the samples in different MTV regimes. The median MTV of each bin was computed, along with the median R1. We fitted the following linear model across samples:

$$R1 = a * MTV + b$$

The slope of this linear model ($a$) represents the R1-MTV dependency. $b$ is constant.
